# Phase transitions of a SEC14-like condensate at Arabidopsis plasma membranes regulate root growth

**DOI:** 10.1101/2022.03.26.485938

**Authors:** Chen Liu, Andriani Mentzelopoulou, Fotini Papagavriil, Prashanth Ramachandran, Artemis Perraki, Lucas Claus, Sebastian Barg, Peter Dörmann, Yvon Jaillais, Philipp Johnen, Eugenia Russinova, Electra Gizeli, Gabriel Schaaf, Panagiotis Nikolaou Moschou

## Abstract

Protein function can be modulated by phase transitions in their material properties, which can range from liquid-to solid-like; yet the mechanisms that drive these transitions and whether they are important for physiology are still unknown. Using the model plant Arabidopsis, we show that developmental robustness is reinforced by phase transitions of the plasma membrane–bound lipid-binding protein SEC14-like. Using imaging, genetics, and *in vitro* reconstitution experiments, we show that SEC14-like undergoes liquid-like phase separation in the root stem cells. Outside the stem cell niche, SEC14-like associates with the caspase-like protease separase and conserved microtubule motors at unique polar plasma membrane interfaces. In these interfaces, SEC14-like undergoes abrupt processing by separase, which promotes its liquid-to-solid transition. The SEC14-like liquid-to-solid transition is important for root developmental robustness, as lines expressing an uncleavable SEC14-like variant or mutants of separase, and associated microtubule motors show similar developmental phenotypes. Furthermore, the processed and solidified but not the liquid form of SEC14-like interacts with the polar protein PINFORMED2 at the plasma membrane and perhaps other polar proteins of the PINFORMED family. This work demonstrates that robust development can involve abrupt liquid-to-solid transitions mediated by proteolysis at unique plasma membrane interfaces.

## Introduction

Under certain conditions, biomolecules can separate from their bulk phase through liquid–liquid phase separation (LLPS), thereby retaining liquid-like properties, such as surface tension, leading to highly circular condensates akin to droplets [1]. LLPS determines the formation of many evolutionary conserved condensates, such as nucleoli, stress granules, and processing bodies. Starting as liquids, some condensates undergo transitions in their material properties that affect their viscosity, surface tension, and degree of penetrance by other molecules. For example, in *Drosophila melanogaster, oskar* ribonucleoprotein (RNP) condensates undergo a liquid-to-solid transition, which is important for the polar distribution of some RNAs in the cell [2]. Whereas *oskar* RNP liquidity allows RNA sequestration, its solid phase precludes the incorporation of RNA while still allowing protein sequestration. Although they are not delimited by membranes, condensates can interface with them or even engulf small vesicles [3].

The past few years have experienced tremendous progress in the evolution of a molecular grammar that underpins LLPS. Molecules such as proteins and RNAs are polymers with attractive groups known as “stickers” that form non-covalent and mainly weak interactions. At certain concentrations which are determined by various factors (e.g., temperature, redox state, pH), interactions are enabled among intra- or intermolecular stickers. When reaching a system-specific threshold concentration, the whole system undergoes LLPS. The stickers promote the attraction between charged residues, dipoles, or aromatic groups that are usually provided by the low-complexity regions so-called “intrinsically disordered regions” (IDRs)[4]. Stickers are connected by “spacers” that regulate the density transitions (i.e., liquid-to-solid transitions) by orienting stickers. The IDRs lack a defined structure and thus can easily expose their stickers. Furthermore, IDRs can increase the apparent size known as hydrodynamic radius adopted by the solvated, tumbling protein molecule [5].

In the model plant, Arabidopsis (*Arabidopsis thaliana*) LLPS is involved in, for example, the internal chloroplast cargo sorting, defence, diurnal rhythms, RNA processing, and environmental sensing by forming plant-specific condensates [6–10]. Furthermore, plants form stress granules and processing bodies to sense and respond to environmental stimuli [11–13]. Condensates of the TPLATE, a plant-specific complex involved in endocytosis can likely form on the plasma membrane [14]. We have also shown that proteins of processing bodies can also form on membranes in Arabidopsis and can attain polarity (i.e., localizing asymmetrically at the plasma membrane) [11]. However, how the condensation of proteins at the plasma membrane is linked to polarity is unclear.

In plants, the few known polar plasma membrane proteins provide crucial information for the robust development [15–17]. One of the most well-studied polar proteins is the auxin efflux carrier PINFORMED2 (PIN2) in the root, which localizes at apical or basal plasma membrane (PM) domains and is required for the formation of the morphogen auxin concentration gradient [18]. Auxin signalling can be modulated by the nucleo-cytoplasmic partitioning of the prion-like transcription factors known as AUXIN RESPONSE FACTORs (ARFs). Intriguingly, during development, ARFs form condensates in the cytoplasm that cannot enter the nucleus, thereby attenuating the auxin signalling [19]. These findings suggest a link between development and condensates in plant roots.

We have previously discovered a link between development and a complex comprising the Arabidopsis caspase-like protease separase (also named EXTRA SPINDLE POLES [ESP]) and three Arabidopsis homologs in the centromeric protein-E-like Kinesin 7 (KIN7), the so-called KIN7.3-clade (KIN7.1, KIN7.3, and KIN7.5). This complex (the kinesin-separase complex [KISC]) is recruited to microtubules (MTs), with the most abundant and important kinesin being KIN7.3 [20]. ESP is an evolutionarily conserved protein responsible for sister chromatid separation and membrane fusion in both plants and animals [21, 22]. ESP binds to the KIN7 C termini, inducing conformational changes that expose the MT-avid N-terminal motor domain of KIN7s, thereby increasing KISC binding atop MTs. The KISC can also modulate polar domains of the PM, as supported by evidence suggesting that the temperature-sensitive *radially swollen 4* (*rsw4*) mutant harbouring a temperature-sensitive *ESP* variant or KIN7.3-clade mutants display reduced PINs delivery at the PM [20]. Yet, how the KISC acts upon PM polar domains to regulate development remains elusive.

Whether condensates interfacing with membranes can undergo transitions in their material properties like cytoplasmic ones and if these changes would have any biological outcome (in any organism) is unclear. Here, we discovered that a previously uncharacterized SEC14-like lipid transfer protein that we named SEC FOURTEEN-HOMOLOG8 (SFH8) recruits KISC to the PM. ESP trimmed SFH8 protein, leading to the conversion of SFH8 from a liquid to a more rigid filamentous phase that remains attached to the PM, an event that we could also reconstitute *in vitro*. This liquid-to-solid transition was associated with SFH8 polarization and robust development of roots. Remarkably, we showed how spatiotemporally confined proteolysis can yield changes in the material properties of proteins and how these properties affect robust development.

## Results

### The KISC Associates with the Lipid-Transfer Protein SFH8 at Polar PM Domains

As the KISC regulates processes that are relevant to the PM (e.g., PIN2 delivery), we aimed to survey an underlying molecular mechanism. We determined that fluorescently tagged or native ESP and KIN7.3 proteins detected by specific antibodies decorate the PM. In the distal meristem (as defined below), both proteins decorated apical domains in the outermost cell layer, the epidermis and basal domains in the adjacent layer, the cortex; the polarity in the vasculature though although evident for the two proteins was harder to determine as these tissues are harder to optically access (**S1A-F Figs**). As KISC proteins lack lipid-binding motifs, we postulated that the KISC associates with the PM via a protein tether, which we sought to identify by screening a yeast two-hybrid (Y2H) library using KIN7.3 as bait. Among the five clones identified, we focused on AT2G21520, as its encoded protein showed peripheral localization reminiscent of the PM when expressed transiently in *Nicotiana benthamiana* leaves (**S2A and B Fig**; **Supplemental Info**). The protein encoded by AT2G21520 is a SEC14-like protein (BLAST: *p* = 1e^−49^) that was ascribed the symbol SFH8, bearing a C-terminal “nodulin”-like motif punctuated with positively charged residues (**S2C Fig; File S1**) [23]. SFH8 is a genuine SEC14-like protein, as the heterologous expression of *SFH8* in budding yeast (*Saccharomyces cerevisiae*) rescued the corresponding temperature-sensitive phenotype of the *sec14-1* mutant (**S2D Fig**) [24]. We confirmed that KIN7.3-SFH8 interacted by co-immunoprecipitation (**Fig 1A**). Furthermore, we established that SFH8 associates with KIN7.3 at the PM, as evidenced by a ratiometric bimolecular fluorescence complementation (rBiFC) assay in Arabidopsis root protoplasts. Unlike conventional BiFC which lacks an internal reference marker, rBiFC can distinguish weak interactions from background fluorescence levels [25]. In rBiFC, we used as a positive control the KIN7.3 interaction with the N-terminus of ESP (aa 1-735), as described previously (**Fig 1B**)[20].

**Fig 1.**
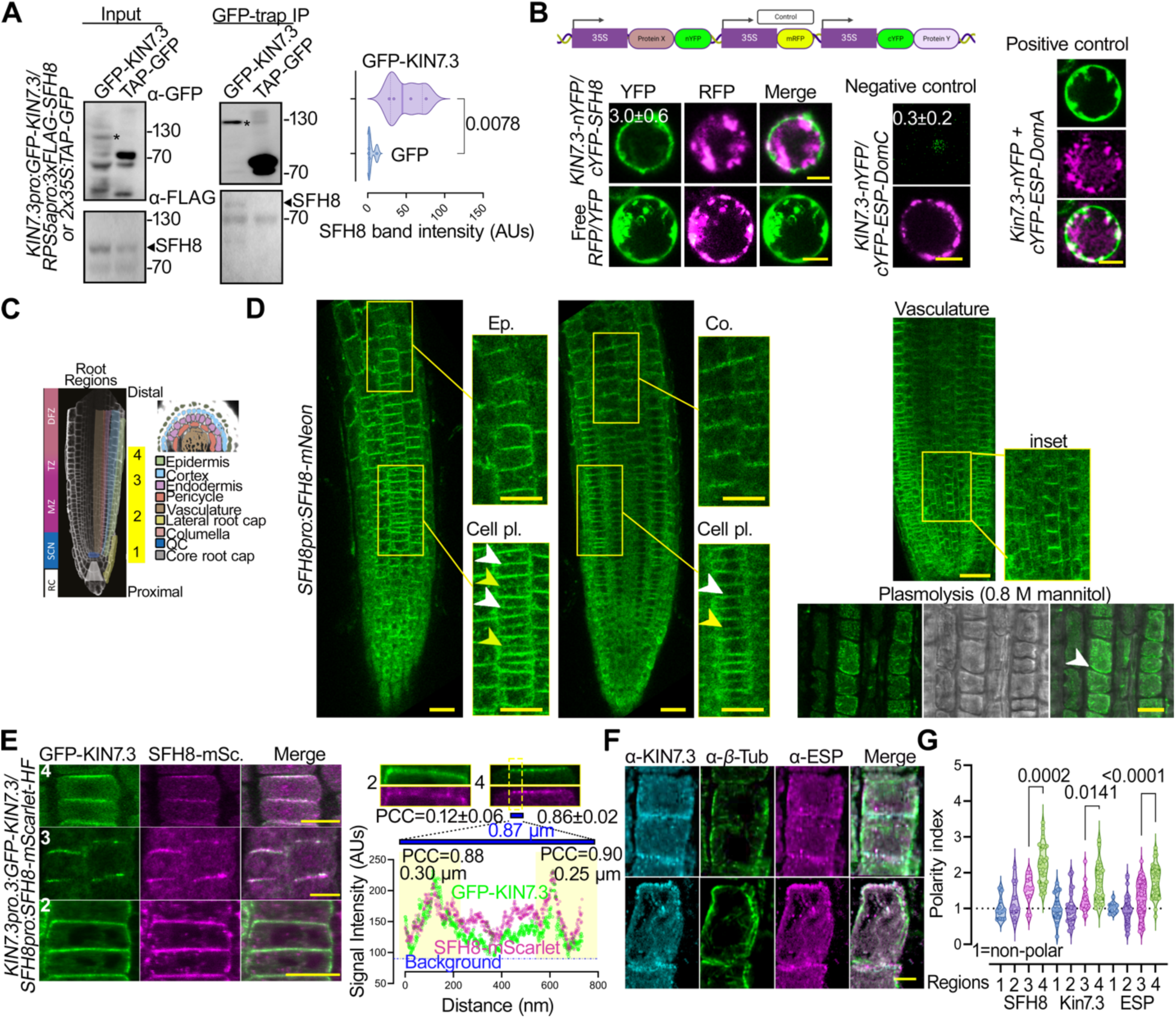
KISC associates with the lipid-transfer protein SFH8 at the PM. **A.** KIN7.3 associates with SFH8 in Arabidopsis. Co-immunoprecipitation from Arabidopsis plants co-expressing *KIN7.3pro:KIN7.3-GFP/RPS5apro:3xFLAG-SFH8* or *2x35Spro:TAP-GFP* (5 days after germination [DAG]); quantification of the interaction (chart showing the SFH8 signal intensity detected by α-FLAG that was pulled-down by GFP or GFP-KIN7.3; n=4, 2-tailed *t*-test). TAP, tandem affinity purification tag. Asterisks denote the full-length GFP-KIN7.3. Note that KIN7.3 is sensitive to proteolytic degradation in the input sample, under the conditions used. **B.** KIN7.3 interacts with SFH8 at the PM of Arabidopsis root protoplasts. Ratiometic BiFC assays showing that KIN7.3 and SFH8 interact at the PM (protoplasts collected 5 DAG; the cartoon on the top shows the construct used and the transcriptional units in the ratiometric BiFC vector). Controls: KIN7.3-nYFP with the ESP-DomA (1-735; positive control) or -DomC (negative control) as defined previously [20]. Mean YFP/RFP signal ratios±SD (n=20) indicated on images. Scale bars, 6 μm. **C.** Root micrograph showing the “4 root regions” examined herein. RC, root columella; SCN, stem cell niche; MZ, meristematic zone; TZ, transition zone; DFZ, differentiation zone; QC, quiescent center. **D.** Tissue-specific expression and subcellular localization of SFH8-mNeon (5 DAG, at the indicated root cell layers). Numbers denote regions according to the root model in **C**. White and yellow arrowheads: mature and expanding cell plates, respectively. The plasmolysis experiment confirms SFH8 signal exclusion from the cell wall (white arrowhead; root region 2). Scale bars, 20 μm. Ep., epidermis; Co., cortex; Cell pl., cell plate. **E.** KIN7.3 and SFH8 colocalize significantly in regions 3 and 4 and show similar polarization patterns in root epidermal cells. Scale bars, 10 μm. Right top: high-resolution signal of KIN7.3/SFH8 at the PM (regions 2 and 4). The overall Pearson Correlation Coefficient (PCC) values for regions 2 and 4 are shown (ROIs: whole image). Note the low PCC value for region 2. For region 4, a plot profile of signal intensity across a straight line of 0.87 μm is shown. The colocalization analyses using PCC) are also shown at the indicated regions of interest (ROIs; rectangular of ∼0.3 μm). AUs, arbitrary units (N=5, n=3, adjacent cells±SD). mScar., mScarlet. **F.** KIN7.3 and SFH8 show similar polarity indexes in root epidermal cells that become significant in regions 3 and 4. Example of α-ESP/α-KIN7.3 colocalization and polarization (counterstained with α-*β*-Tubulin; region 3). Scale bars, 5 μm. **G.** Quantification of SFH8, KIN7.3, and ESP polarity in regions 1-4 (values >1 represent polarized proteins, and calculations are described in **S1A Fig**; N=5, n≥25; Dunnett).

The localization and functions of SFH-like proteins in Arabidopsis are unknown, while genetic evidence suggests that SFH1 is essential for root hair development [23]. We expressed *SFH8- mNeon* under the *SFH8* promoter (*SFH8pro*) to explore its localization; the SFH8-mNeon signal was observed in the root meristem. To expedite our localization analyses, we defined four developmental root regions: core meristem (1; stem cell niche), meristematic zone (2; proximal meristem), meristematic/transition zone (3), and late transition zone (distal meristem; model of different root regions described in **Fig 1C**) (4). We detected SFH8-mNeon signals in all meristematic root cells at the PM, decorating apical PM domains in the epidermis and basal domains in the cortex/vasculature in distal meristem cells (**Fig 1D**), like the ones shown for KISC proteins above. Accordingly, SFH8 colocalized at the PM with KIN7.3, and this colocalization was significant in regions 3 and 4, as revealed by analysis of signal collinearity in super-resolution micrographs (120 nm) using the Pearson correlation coefficient (PCC) to quantify the levels of colocalization (**Fig 1E**, right chart and below). Furthermore, SFH8 and KISC proteins attained significant and similar polarity in regions 3 and 4, localizing to basal (in the cortex) or apical domains, but not in regions 1 and 2 (**Figs 1F and G; S1E Fig**). Later, we discuss this polarization in more detail, but altogether these results suggest the association of KISC with SFH8 at polar domains of the PM in root cells and suggest that SFH8 could be a tether for KISC.

### SFH8 Clusters Recruit the KISC Where ESP Cleaves SFH8 Creating Filaments

The interactions between KISC-SFH8 prompted us to examine whether SFH8 is tethering KISC at the PM. We thus identified two T-DNA insertion mutants in *SFH8*, designated *sfh8-1* and *sfh8-*2. We continued further analyses with the *sfh8-1* background (hereafter “*sfh8*”) because as explained later it is phenotypically similar to *sfh8-*2 (**Fig 2A**). In *sfh8*, GFP-KIN7.3 displayed both a reduced PM localization and polarity compared to that in the wild type (**Fig 2B**). While SFH8- mNeon tethering at the PM did not appear to depend on KISC, SFH8-mNeon was highly apolar in all cell types examined (**Fig 2C**). To further validate this result we used an inducible system that leads to the overaccumulation of the KIN7.3 C-terminal tail (*XVEpro>KIN7.3pro:HA- KIN7.3tail*) with the ability to disrupt KISC, as it titrates ESP out of the active KISC [20]. Thus, the transient KISC depletion led to a loss of SFH8 polarity within 2 days (**Fig 2D**). Hence, KISC and SFH8 synergistically define their localization: SFH8 tethers KISC at the PM, and in turn, KISC promotes SFH8 polarization.

**Fig 2.**
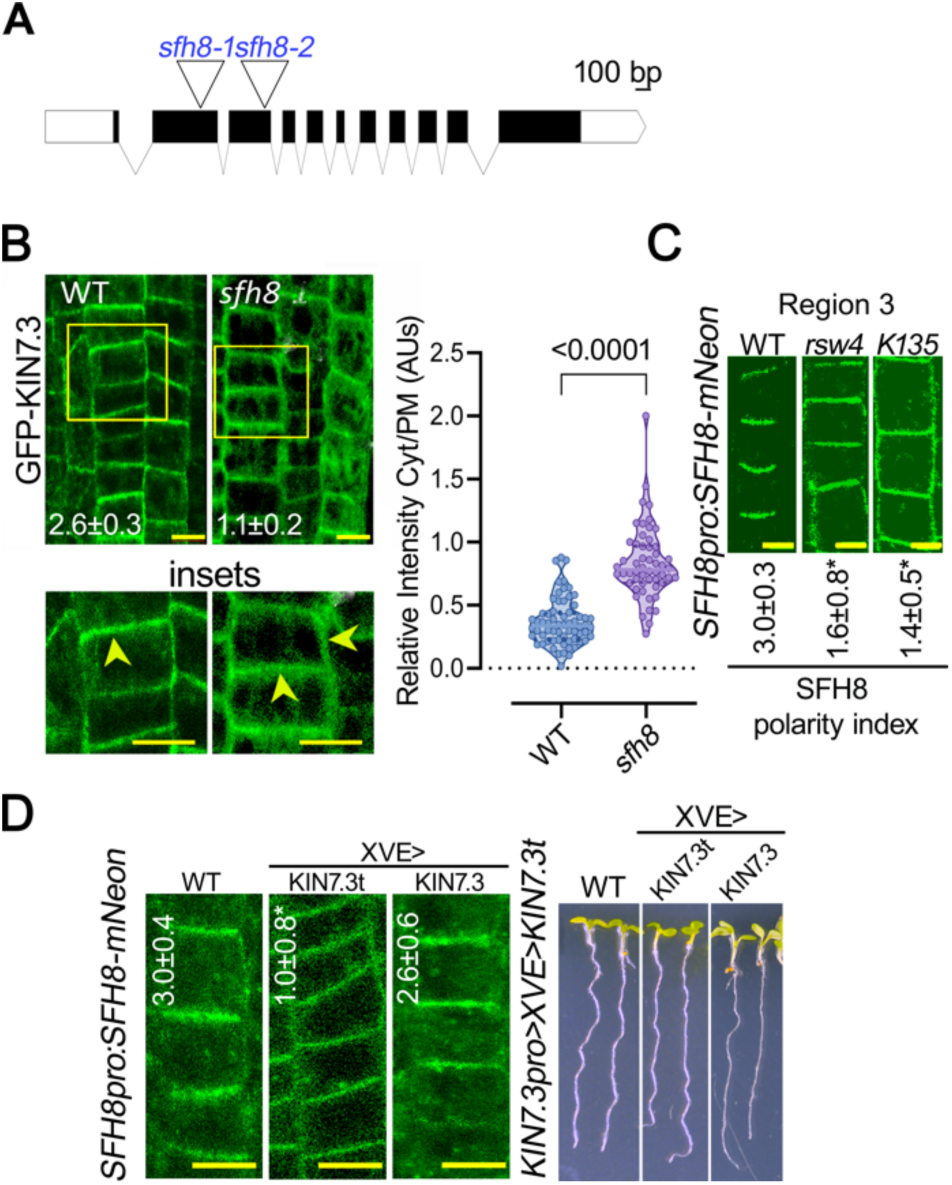
SFH8 modulates development and recruits KISC at the PM. **A.** Schematic representation of T-DNA insertion sites for *sfh8-1* and *sft8-2*. **B.** SFH8 and KIN7.3 localizations at the PM are interdependent. GFP-KIN7.3 PM localization in WT or *sfh8* (5 DAG, region 3). The polarity index of KIN7.3 is also shown on the images (three pooled experiments from region 3±SD; n=40; differences were significant at p<0.0001, Dunnett). Scale bars, 5 μm; quantification of cytoplasmic to PM signal (right; a single representative experiment replicated multiple times; n=25; 2-tailed *t*-test). Arrowheads in the insets show apical or lateral localization of KIN7.3. **C.** SFH8-mNeon localization in WT, *rsw4*, and *k135* (5 DAG, region 3). Images are representative of an experiment replicated multiple times for polarity. The polarity index of SFH8 is also shown (three pooled experiments from region 3±SD; n=40; “*”: p<0.0001 to WT, Dunnett). Scale bars, 5 μm. **D.** Perturbations of SFH8 polarity through KISC disruption. Perturbed gravitropism and growth of lines over-expressing transiently KIN7.3 tail (“t”; 24-36 h, 2 μM estradiol). Scale bars, 8 μm. Right: SFH8 polarity loss in *KIN7.3pro˃XVEpro˃KIN7.3t* lines (region 3, epidermal cells). Images are representative of an experiment replicated multiple times for polarity. Scale bars, 10 μm. SFH8 polarity index is also shown on the images (three pooled experiments from region 3±SD; n=40; “*”: p<0.0001 to WT, Dunnett).

Interestingly, in follow-up experiments aiming at studying in detail the localization of SFH8 in the loss-of-function KIN7.3-clade mutant *kin7.1 kin7.3 kin7.5* [*k135*] and *rsw4* backgrounds, we observed that the full-length SFH8 levels increased in these two mutants. Immunoblot analysis of lines expressing a construct encoding SFH8 with a hexahistidine-triple-flag (referred to as HF)- mScarlet tag at the C or N terminus (∼106 kDa) under the control of the *RPS5a* promoter showed increased SFH8 abundance in *k135* and *rsw4* backgrounds (**Fig 3A**). The *RPS5a*, a meristematic specific promoter here was used as the *SFH8pro* could not lead to a detectable signal in immunoblots. Given the increased abundance in these KISC mutants of SFH8 and considering that ESP is a protease, we decided to examine the possibility that KISC regulates SFH8 protein levels. The reduced abundance of SFH8 levels in the wild type compared to that in *k135* and *rsw4* associated with a presumptive ∼40-kDa (or ∼10 kDa excluding mScarlet) N-terminal cleavage product (**Fig 3A,** right). We, thus, speculated that in the presence of KISC, SFH8 is cleaved from ESP at its N terminus, producing this 10kDa product (hereafter, identified as “cleavage product”). We followed up this cleavage process in vivo using lines harboring the *SFH8pro:SFH8* construct with a C- or N- terminal fusion of the mNeon fluorescent tag. We observed fluorescent cytoplasmic puncta that accumulated gradually during development in *SFH8pro:mNeon-SFH8* lines in region 3 and mainly in region 4 that, as shown above, KISC shows strong colocalization with SFH8; these foci were absent from the lines harboring the *SFH8pro:SFH8-mNeon* construct encoding a C-terminal mNeon fusion with SFH8, where the SFH8 signal was mostly on the PM (**Fig 3B**; regions 3, 4), suggesting that only the N-terminus of SFH8 is cleaved and released in the cytoplasm. These results altogether suggest that SFH8 is cleaved by ESP at the N terminus part, creating a cleavage product in the form of cytoplasmic puncta, and this cleavage is progressive along the developmental root axis.

**Fig 3.**
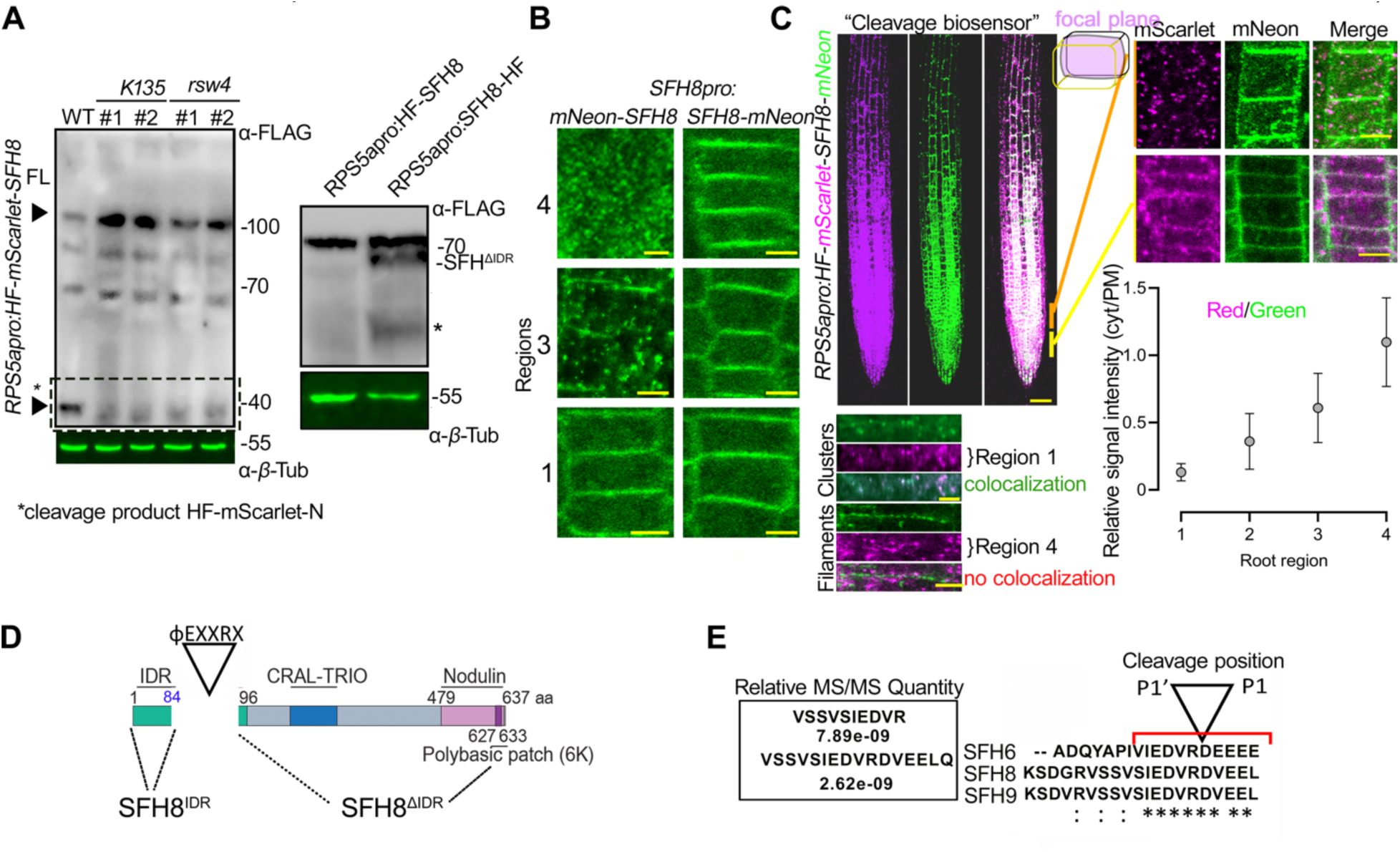
KISC trimming of SFH8 promotes the cluster-to-filamentous transition. **A.** SFH8 releases an N-terminal fragment in a KISC-dependent manner. Western blot showing the accumulation of an SFH8 N-terminal fragment (black arrowhead at ∼40 kDa; FL, full-length) in WT, *k135,* or *rsw4* (24 h at the restrictive temperature 28°C, two lines; lower) expressing *HF-mScarlet-SFH8*. Two independent lines obtained by transformation are shown (designated as 1 and 2). Note the reduced accumulation of the N-terminal fragment in *k135* and *rsw4*. Right: western blot showing the remaining C-terminal SFH8 fragment (asterisk, an additional truncated product of low abundance) in WT lines expressing *RPS5apro:HF*-tagged SFH8. Immunoblots are representative of experiments replicated multiple times. **B.** N-terminal fragment production of SFH8 is developmentally regulated and it accumulates in the cytoplasm. Micrographs from WT lines expressing *SFH8* N- or C-terminally tagged with mNeon under the *SFH8* promoter (3 regions, epidermis; 4^th^ region slightly below the transition zone in this case). Note the abrupt formation of N-terminal bodies in the cytoplasm of lines expressing *mNeon-SFH8* at region 3, and the reduction of the corresponding PM signal. Scale bars, 4 μm. **C.** An SFH8 cleavage biosensor reveals that the N-terminal fragment and the remaining C- terminus of SFH8 are apart during development. Localization and expression of the biosensor (scale bar, 50 μm), details (right, mid-plane focus of cortex, scale bars, 4 μm), and relative signal intensity of cytoplasmic versus PM signal (chart). Data are means ± SD (N=10, n=10). Super-resolution imaging (120 nm) of cluster-to-filament conversion (regions 2 to 4)(lower). Note the absence of mScarlet signal from filaments. **D.** SFH8 protein architecture (IDR, intrinsically disordered region, CRAL-TRIO: active site for SEC14 proteins, and NOD: nodulin motif). **E.** SFH8 is cleaved at a specific site. SFH8 IDR peptides identified in *pRPS5a:SFH8-mScarlet- HF* pull-down experiments coupled with LC-MS/MS. The cleavage motif of ESP on SFH proteins, φEXXR is conserved (presented here for three SFH protein paralogs, SFH6/8/9; upper left).

To dynamically follow SFH8 cleavage *in vivo*, we established a double-labelled N- terminal/C-terminal tagged fluorescent SFH8 (cleavage biosensor; *RPS5apro:mScarlet-SFH8- mNeon*), whereby cleavage would disrupt the colocalization of mNeon and mScarlet signals. Indeed, we observed a lack of mNeon/mScarlet colocalization in region 3 and mainly in 4 (**Fig 3C**). High-resolution imaging defined a more clustered form of SFH8 in region 1 at the PM (where both signals colocalize, indicative of an uncleaved SFH8 biosensor) and a more filamentous form in regions 3 and 4 of the C terminal part of the SFH8 protein (where no colocalization is observed) (**Fig 3C,** detail and graph). We thus showed that SFH8 transitions from a cluster (full-length protein) to a filament (C terminal part) upon its cleavage from ESP and this transition is detectable at region 3 onwards (**Fig 3C**).

We also aimed at defining the exact cleavage site within SFH8. Accordingly, we immunoprecipitated SFH8-mScarlet-HF using α-FLAG and quantified the abundance of SFH8 peptides via mass spectrometry (MS), resulting in the identification of a potential cleavage site right after the residue R84. The size of the predicted cleavage fragment was in good agreement with the western blots shown in **Fig 3A** (∼10 kDa). The I80EDVR84D sequence corresponded to the reported non-plant ESP cleavage consensus φEXXR [26, 27], also found in other SFH8-like proteins (**Fig 3D and E, S3 File**). By establishing an in vitro ESP cleavage assay we confirmed that immunopurified ESP, mitotically activated through co-expression with *Cyclin D* [28], can cleave recombinant glutathione *S*-transferase (GST)-SFH8 at R84; we validated our assay by showing the cleavage of mitotic cohesin (SYN4), the well-known target of ESP (**S3A-D Figs**) [29]. Hence, the transition of SFH8 to filaments depends on the cleavage at R84.

### SFH8 Forms Transient Clusters with Liquid-Like Properties that Exclude KISC

We further aimed to follow the colocalization and cleavage of KISC/SFH8 in more detail. As the KISC binds MTs, we postulated that SFH8 and the KISC might co-partition in MT filaments in proximity to the PM. Contrary to our expectations, the standard MT marker MAP4^MBD^ (the microtubule-binding domain of MICROTUBULE-ASSOCIATED PROTEIN4) or β*-*tubulin showed only partial colocalization with KIN7.3 at the PM, while amiprophos-methyl (APM) that disassembles MTs (10 nM; [20]) did not alter KIN7.3 localization at the PM (**S4A and B Figs**). In Arabidopsis roots, SFH8 filaments were short (<0.5 μm) and insensitive to APM treatment; ESP decorated similar filaments as shown in root cells expressing ESP under an inducible promoter driving expression at KIN7.3 domains (*KIN7.3pro>XVEpro>GFP-ESP/RPS5apro:SFH8- mScarlet*; **S4C Fig**). These results suggest that SFH8 and KISC at the PM do not remain attached to MTs.

As these results suggested that the KISC and SFH8 co-assemble in filaments most of which did not relate to MT, we aimed at deciphering this localization in detail. We thus examined the localization of KISC components and SFH8 in Arabidopsis roots by total internal reflection fluorescence (TIRF) microscopy, which is suitable for analyzing the PM due to the shallow illumination penetration (decay constant of ∼100 nm). By focusing on lateral cell junction domains (**Fig 4A**; regions 3 and 4; 3-5 days post-germination), we determined that SFH8-mNeon segregates into at least two major populations: (i) immobile filaments that co-localize with KIN7.3- tagRFP and (ii) mobile or immobile KIN7.3-tagRFP-independent cluster-like structures (**Figs 4B-E**). These results are consistent with the observation of SFH8-KISC localization in clusters and filaments (**Fig. 3**). The mobile SFH8 clusters showed little diffusion at the PM and occasionally fused, properties that are reminiscent of cellular condensates that sometimes form through LLPS (**Movies S1-3, Figs 4C and D**). We also observed some small non-diffusing clusters with reduced circularity (**Movies S2, 3**) that may show intermediate phases between the cluster state (droplet-like) and the filamentous state.

**Fig 4.**
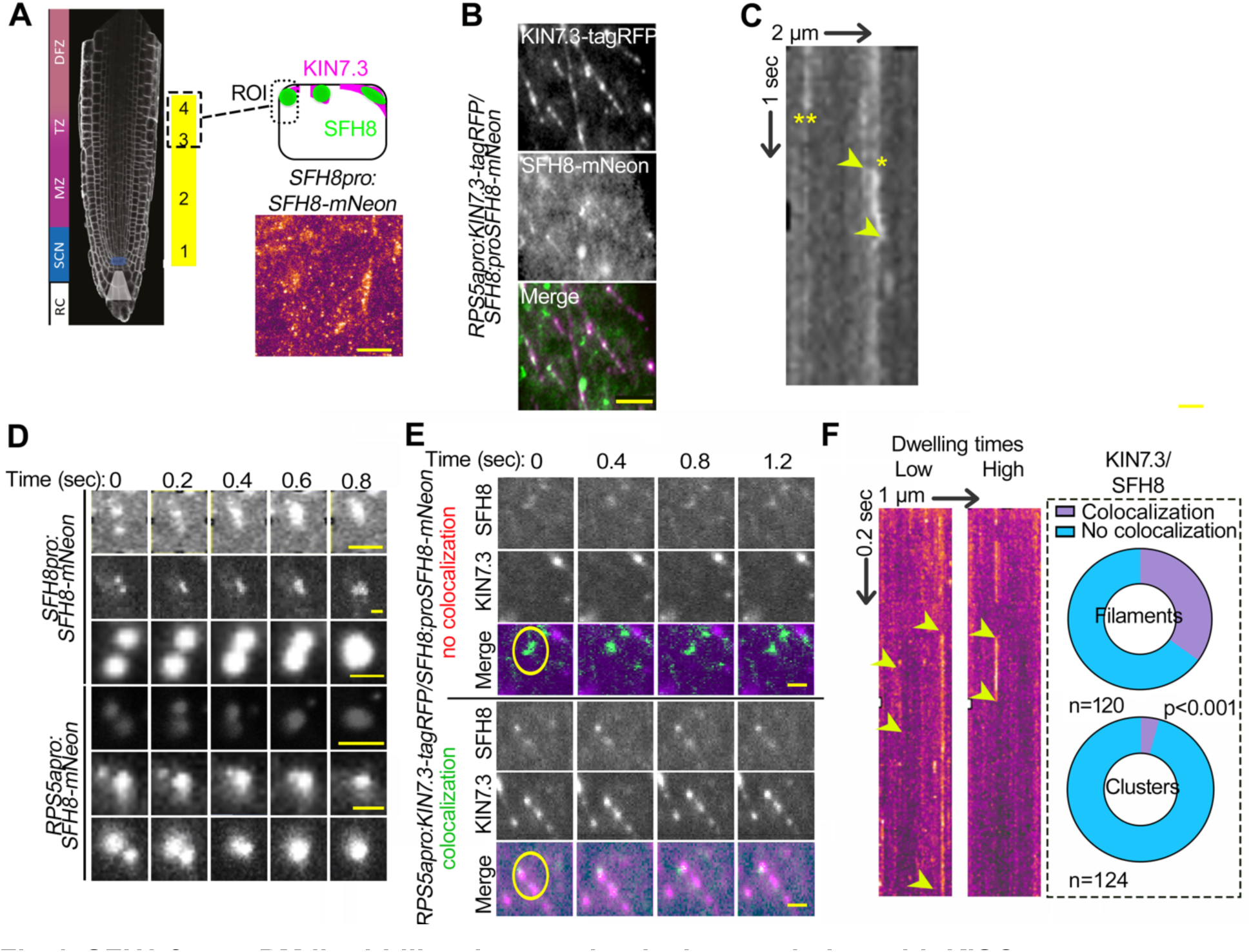
SFH8 forms PM liquid-like clusters that lack association with KISC. **A.** Establishment of a TIRFM setting for visualization of SFH8 at the PM. The model here is showing the region used for imaging, and an SFH8 localization overview (3-5 DAG, region 3). Scale bar, 2 μm. Note that in TIRFM imaging, the focal plane is restricted to the outermost tissues and therefore epidermis of region 1 or 2 cannot be examined (see **Figure 1C** for a root model showing that the epidermis in this region is encapsulated by the root cap). **B.** Example of a dual-channel TIRFM of lines expressing *SFH8-mNeon* and *KIN7.3-RFP*. Micrographs are representative of an experiment replicated multiple times. Scale bar, 0.3 μm. **C.** SFH8 forms clusters that show lateral mobility at the PM plane. Kymograph showing diffusing (*) and non-diffusing (**) clusters. Arrowheads indicate the spatial offset of the diffusing cluster (lateral diffusion on the PM plane ∼200 nm). The spatiotemporal resolution is shown by the bars (2 μm and 1 sec). **D.** SFH8 clusters fuse with one another at the PM plane. Examples of SFH8-mNeon clusters from micrographs obtained by TIRFM fusing on the PM, irrespective of the promoter used (*SFH8pro/RPS5apro*). Note that both promoters produced clusters of similar sizes. Scale bar, 0.3 μm. **E.** Dual-channel TIRFM of SFH8-mNeon/KIN7.3-RFP co-expressing line showing SFH8 clusters and the formation of filaments that do not diffuse. Note the lateral diffusion of SFH8 clusters and the lack of filaments motility (circles). Scale bars, 0.3 μm. **F.** Kymographs show diffusing clusters with low and high dwelling times at the PM. Right: charts show KIN7.3 and SFH8 colocalization percentage in clusters or filaments (three pooled experiments; 5 fields of view, n, as indicated; Wilcoxon).

KIN7.3 and SFH8 co-partitioned in short filaments but not in SFH8-decorated clusters; these clusters showed variable residence times at the PM, unlike filaments that were permanently assembled at the PM (**Fig 4C and E-G**). We wished to determine why the KISC was excluded from the SFH8 clusters; we hypothesized that converting the polybasic charge of the SFH8 nodulin patch to a hydrophobic region would promote the clustered (condensed) state of SFH8. Indeed, replacing six pertinent lysines (K) with alanines (A) in the nodulin patch artificially increased SFH8 clustering in *N. benthamiana* leaves (although also exhibiting a slightly reduced localization at the PM) and decreased its association with KIN7.3 (**S4D-F Figs**). In Arabidopsis, mNeon-SFH8^6KtoA^ showed reduced localization at the PM, reduced filaments, and lacked polarity (**S4G Fig**). This result suggested that hindering the interaction between SFH8 and the KISC by reducing the accessibility to the SFH8 blocked SFH8 filamentous transition, suggesting that filaments are produced through KISC where KISC-SFH8 remain associated. On the other hand, the SFH8 clusters have liquid-like properties.

### The N-terminus of SFH8 Defines its Liquid-like Properties

To address the link between KISC removal of the N-terminus and changes in SFH8 structure at the PM (i.e., the filamentous formation), we first used fluorescence recovery after photobleaching (FRAP). Owning to their increased mobility compared to filaments as shown by TIRFM, we anticipated that cells with liquid-like SFH8 clusters would show increased FRAP rates. Indeed, SFH8-mNeon recovered rapidly at the PM close to the meristem (regions 1 and 2), unlike the distal meristem (regions 3 and 4) in which SFH8 lacked recovery, as it converted to filaments (**Fig 5A**). As this filamentous transition of SFH8-mNeon was also reduced in the KISC mutants (**Fig 5B**; >3-fold), these findings further genetically confirm that KISC mediated the conversion of SFH8 clusters to solid-like filaments. These filaments are more stably attached to the PM as they do not show recovery in FRAP and mobility in TIRFM (see above) and can retain an association with the KISC (**Fig. 5A model**).

**Fig 5.**
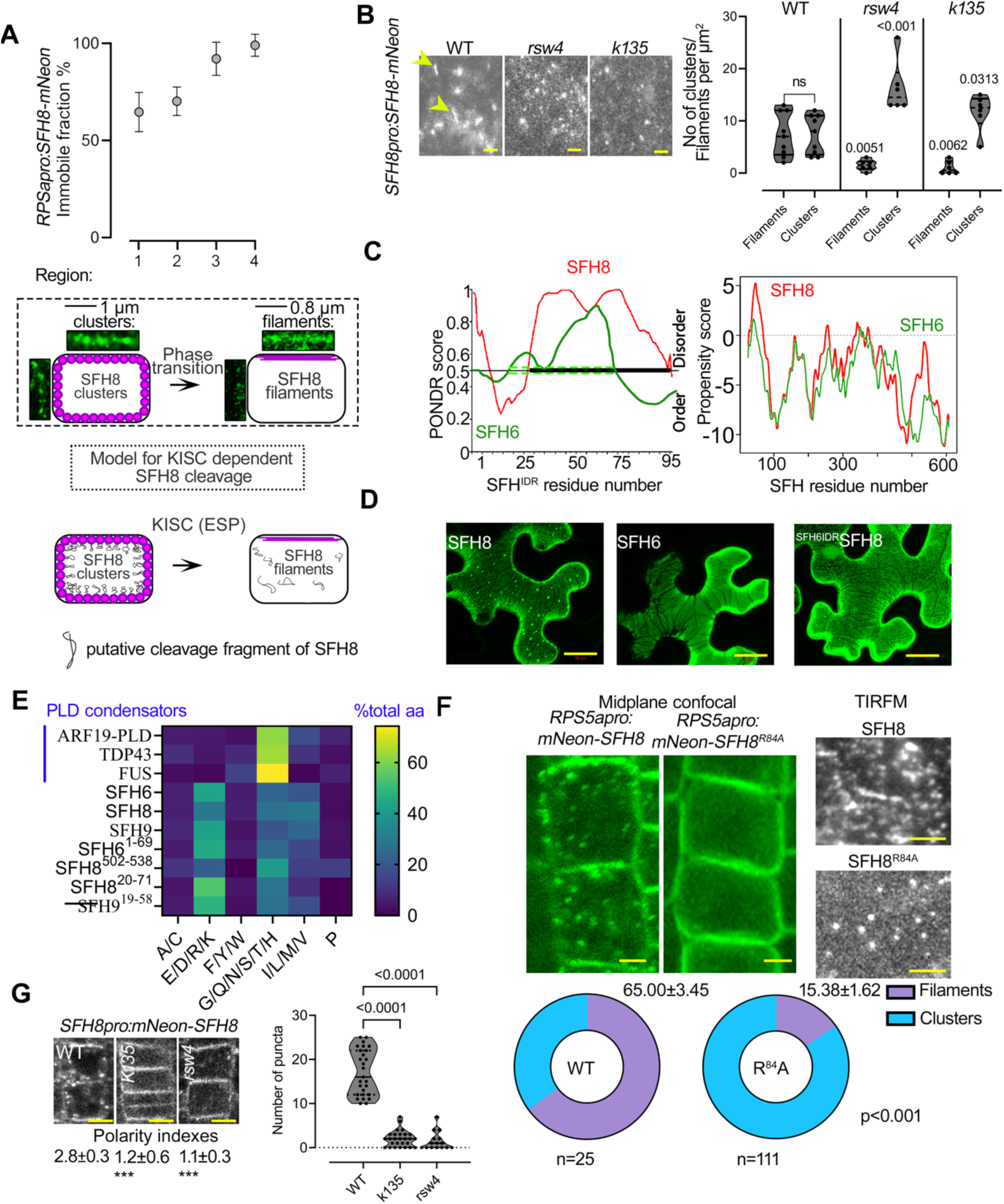
KISC abrogation retains the clustered phase of SFH8 through SFH8 N-terminus. **A.** SFH8-mNeon is immobilized abruptly at region 3 onwards through the formation of filaments as determined by FRAP. Data are means ± SD (N=1, n=10 roots at each point). Bottom: clusters and filaments in two root regions (cortex cells) determined by super-resolution confocal microscopy (120 nm, 40 frames per second, midsection; regions 1 and 4, left and right, respectively) and a model showing the SFH8 clusters-to-filaments conversion. **B.** SFH8-mNeon localization in WT, *k135,* and *rsw4* root cells (5 DAG, region 3, TIRFM; arrowheads indicate filaments). Scale bar, 0.5 μm; quantification of SFH8-mNeon clusters and filaments in WT, *k135,* and *rsw4* (right; n=28, 5 fields of view; Wilcoxon) and a model showing the dependency of SFH8 clusters-to-filaments conversion on KISC. Clusters with circularity below 0.5 were defined as filamentous. **C.** *In silico* predictions of IDRs by PONDR, and phase separation propensity determined by catGRANULE [79] for SFH8 and SFH6. Note the shorter IDR length and lower LLPS propensity score for SFH6. **D.** The N-terminal SFH8 IDR drives phase separation of SFH8 and it is released in the cytoplasm upon cleavage in a heterologous expression system. Reduced puncta formation in a chimeric protein of SFH6^IDR^ and the C-terminal SFH8 in *N. benthamiana* leaves, in the presence of ESP/CyclinD; see also **S2 Fig** for the activation of ESP protein by CyclinD). Scale bars, 20 μm. **E.** Comparative analysis of the SFH8 IDR composition. Each amino acid is assigned to one of six groups on the x-axis, and the fraction of grouped amino acids are shown. For comparison, model “condensators” are shown (ARF19 to FUS). SFH8 IDR is similar to the average IDR signature of disordered proteins in Arabidopsis [19]. The lengths of the IDRs were determined by the fIDPnn [80]. **F.** Localization of SFH8^R84A^ in region 3 epidermal cells. Scale bars, 2 μm. Right: persistence of PM SFH8 condensates in *sfh8 SFH8^R84A^* lines (TIRFM; scale bar, 0.2 μm). Bottom: quantification of mNeon-SFH8 or mNeon-SFH8^R84A^ clusters and filaments (four independent experiments; n as indicated, 2-tailed *t*-test). **G.** Reduced puncta formation and increased mNeon-SFH8 PM (apolar) signal in the *k135* or *sfh8* (7 DAG, epidermis of region 3). Scale bars, 5 μm. Lower: puncta quantification (single representative experiment; n=27 cells from region 3; ANOVA). Polarity indexes of SFH8-mNeon are also shown (“***”: p<0.0001 to WT, Dunnett).

Next, we asked whether the removal of the N-terminal SFH8 fragment can drive a faster cluster-to-filament transition in SFH8. Through *in silico* predictions, we determined that the cleaved N-terminal fragment of SFH8 is an IDR (hereafter SFH8^IDR^; **Fig 5C**), suggesting that it is unlikely to present well-defined structures. We established that this protein architecture is conserved throughout the evolution of SFH-like proteins, which might imply the functional importance of this IDR (**S2 File**). As IDRs are enriched in proteins undergoing LLPS [30], we tested whether full-length SFH8 undergoes LLPS. This test was also justified considering the LLPS features of SFH8 clusters at the PM mentioned above in **Fig 4D** above. SFH8^IDR^ was predicted as an inducer of LLPS through the catGranule algorithm, while the corresponding region of a close SFH8 homolog, SFH6 (SFH6^IDR^), was predicted to exhibit a reduced propensity to undergo LLPS; we verified this prediction in *N. benthamiana* (**Fig 5C and D**). SFH8^IDR^ sequence composition is distinct from that of animal proteins that undergo phase transitions in the cytoplasm but show an amino acid distribution like that of the average IDR profile for Arabidopsis (**Fig 5E**) [19]. We further observed that puncta formed by the SFH8^IDR^ failed to colocalize with vesicular markers, cellulose synthase complex or mitochondria (**S5A and B Figs**) and showed LLPS hallmarks such as droplet-like dynamic morphology with frequent splitting, fusion, and interconnections (**Movie S4, S5C Fig**). FRAP analysis of these produced mNeon-tagged SFH8^IDR^ puncta demonstrated a rapid signal recovery (t_1/2_∼10 s, mobile fraction 40%) and sensitivity to 1,6-hexanediol, which may block LLPS (**S5C Fig**) [31, 32]; 1,6-hexanediol also led to the dissolution of SFH8 clusters on the PM but not of filaments (**S5D Fig**), confirming their more rigid structure.

Since SFH8 clusters displayed sensitivity to hexanediol, rapid FRAP, and fused akin to cellular condensates, we speculated that SFH8 clusters at the PM may form by LLPS much like SFH8^IDR^. *In silico* prediction, using the algorithms PLAAC and CIDER showed that SFH8 can adopt context-specific conformational states with an absolute value of net charge per residue (NPCR) of 0.014, which suggests that is a polyampholyte. This result suggested that the propensity of SFH8 to undergo LLPS may be sensitive to the environments and that SFH8 may represent an ensemble of conformers. The highly electronegative field of the PM where SFH8 accumulates may lead to the unbalancing of opposite charges upon IDR removal (IDR NCPR = 0.071) due to repulsive forces, which would support an extended structure like a filament. Indeed, we established that LLPS of SFH8 relies on the N-terminal IDR, as swapping it with the corresponding region from the SFH6, reduced clustering at the PM and the formation of puncta in the cytoplasm in the presence of ESP (**Fig 5D**). Overall, our analyses suggest that SFH8 behaves like an LLPS polyampholyte at the PM with negatively charged lipids (phosphatidyl-inositols; PIs), likely buffering repulsive charges that would otherwise restrict condensation.

To further examine LLPS of SFH8, we established an in vitro LLPS assay with fluorescently labeled proteins using thiol-reactive maleimide dyes (see **Materials and Methods**). Under conditions that promote phase separation (**S6A Fig** for protein purification), SFH8 or the uncleavable variant SFH8^R84A^ rapidly formed condensates at relatively high concentrations (5 μΜ), while SFH8^ΔIDR^ (for delta IDR, i.e., SFH8 without the IDR) formed filament-like assemblies in good agreement with the in vivo situation (**S6B Fig**). Consistent with the in vivo data, the in vitro filaments did not show recovery of fluorescence in FRAP experiments and showed reduced circularity compared to SFH8 condensates (**S6C Fig**). As the bulk phase separation tests here are likely not relevant to the phase behavior at the PM, we also established a system to test SFH8 phase separation on membranes. We thus used SUPER templates (supported lipid bilayers with excess membrane reservoir) that contain low-tension membranes surrounding a silicon bead [33]. GST-SFH8 formed large droplets on SUPER templates rich in PI lipids (i.e., PIP2, as has been shown that yeast Sec14 binds on these lipids), at lower concentrations compared to the bulk-phase experiments (**S6D Fig**; 0.1 μΜ vs. ≥5 μM in the bulk phase). This result suggested that membranes promote LLPS of SFH8. By contrast, GST-tagged SFH8^ΔIDR^ did not show similar behaviour in this setting and formed oligomers in native polyacrylamide gel electrophoresis; this behaviour could be also observed for SFH8 in the presence of KIN7.3/ESP (**S6E Fig**). Hence, as suggested above, PIs may neutralize electrostatic repulsions via counterion-mediated charge neutralization along SFH8, as suggested for other proteins [34], thereby mediating LLPS. It is worth noting that, consistent with our data, the threshold concentration for LLPS in 2D systems like the PM can be an order of magnitude lower than in 3D systems (for example [35]).

As the N-terminal IDR drives the phase behaviour of SFH8, we speculated that an SFH8 uncleavable variant would fail to undergo a liquid-to-solid transition, as the IDR would be always present. Indeed, in lines expressing the uncleavable variant *RPS5apro:mNeon-SFH8^R84A^*in the *sfh8* background, cells lacked cytoplasmic fluorescent puncta, while SFH8^R84A^ was apolar much like SFH8 in KISC mutants and did not convert to filaments (**Fig 5F and G**). As expected, the mNeon-SFH8 fluorescent protein produced by *SFH8pro:mNeon-SFH8* lines showed higher FRAP rates on the PM (due to cleavage and liquidity of clusters), as expected, unlike the corresponding C-terminally tagged SFH8-mNeon (**S7 Fig**). Altogether, these results suggest that SFH8 clusters exhibit LLPS properties and that SFH8 (initially in LLPS) releases two proteolytic proteoforms upon cleavage by KISC: C-terminal SFH8^ΔIDR^ (converted to solid-like filaments) and the N-terminal SFH8^IDR^ (cytoplasmic liquid-like puncta) that defines liquidity of SFH8.

### SFH8 Phase Separation Enables Delivery of Some Polar Proteins

SFH8^IDR^ is a polyampholyte and may act as an entropic bristle through random movements around its attachment point on lipids which could in theory exclude access of other proteins to PM regions where uncleaved SFH8 resides. This property would also reduce the probability of full-length SFH8 undergoing filamentous transition due to inter- or intramolecular repulsion. We thus aimed to decipher the biological significance of SFH8 phase transitions at polar domains. As a relevant readout here, we used PIN2 because KISC plays a role in PIN2 delivery [20], but this choice is not implying a strict link between SFH8/KISC to auxin signaling. We observed that the PIN2-GFP (or α-PIN2 by immunohistochemistry) signal is lower by about 50% at the PM of *sfh8* or KISC mutants (**Fig 6A and B**), suggesting that SFH8/KISC are required for PIN2 delivery and/or retention on polar domains. Notably, PIN2 accumulated in endosome-like structures in *sfh8*, while KISC or *sfh8* mutants showed reduced PIN2 polar delivery mainly in the cortex (**Fig 6B and C**; regions 3-4). Furthermore, the uncleavable variant of SFH8^R84A^ could not rescue the PIN2 polarity/levels defects of *sfh8* (**Fig 6C**; regions 3-4). We also observed increased localization to endosomes and a reduced delivery for PIN1 in *sfh8* (likely not for other PINs), but not for non-polar proteins (the H^+^-ATPase [AHA1] or PLASMA MEMBRANE INTRINSIC PROTEIN 2a [PIP2a]), discounting a general role for SFH8 in exocytosis (**S8A and C Figs**). These results suggest that cleavage of SFH8 and thus its conversion to filaments is required for the establishment of a subset of polar domains.

**Fig 6.**
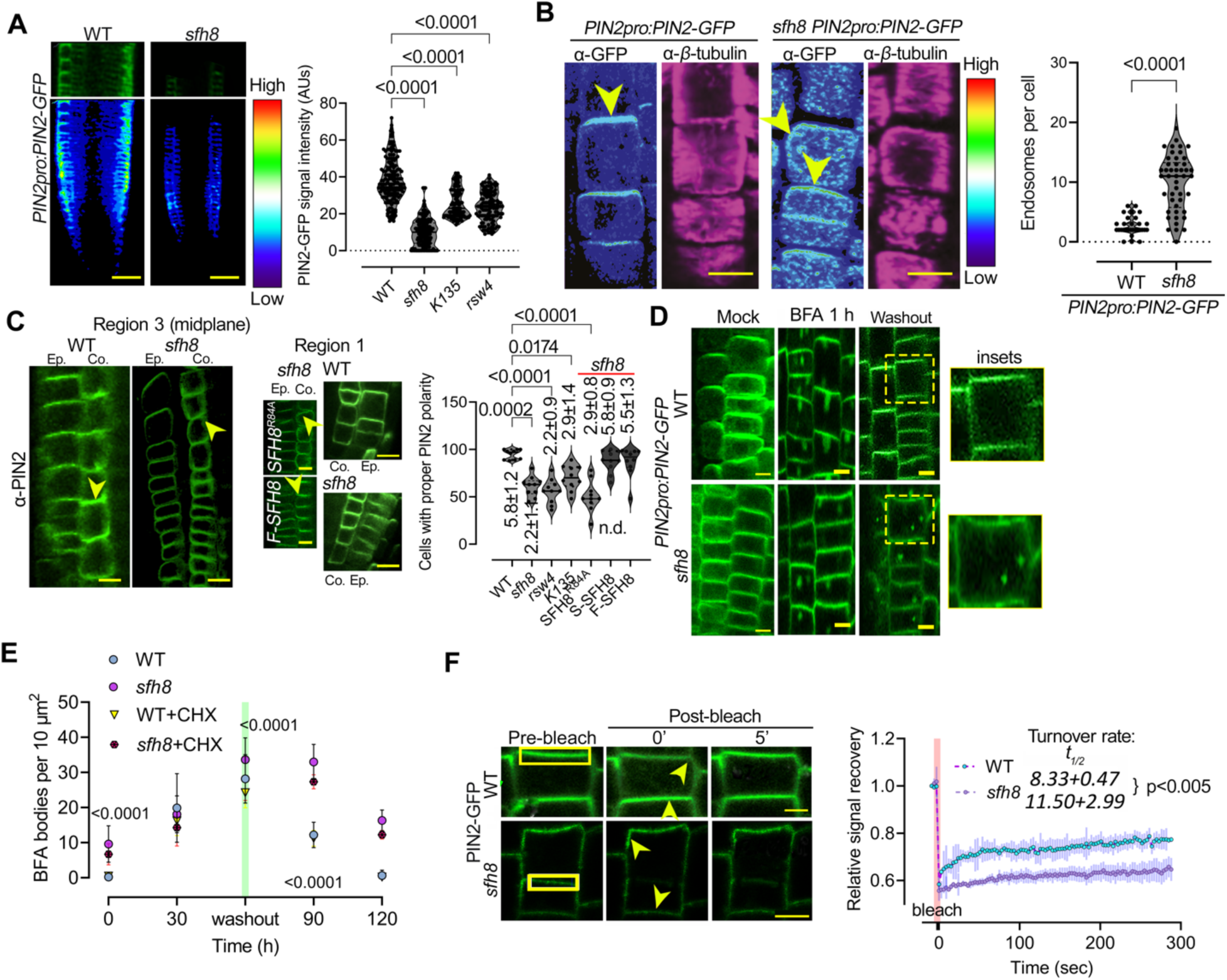
SFH8 can affect PIN2 dynamics at the PM. **A.** PIN2 levels at the PM are reduced in *sfh8*, *k135*, and *rsw4*. PIN2-GFP signal intensity (color-coded) in WT and *sfh8* and quantification (right) of PIN2-GFP on PM of WT, *sfh8*, *k135*, and *rsw4* (7 DAG, region 3; N=2 pooled experiments; n=164-203; multiple comparisons Dunnett). Scale bars, 50 μm. **B.** PIN2 localization in WT and *sfh8* (color-coded; counterstaining with a*-β*-tubulin to show focal plane; magenta; 5 DAG, region 3). Arrowheads indicate PIN2 accumulation maximum, and in the *sfh8* slight polarity offset. Note the high number of PIN2-positive endosome-like structures in the *sfh8*. The *sfh8* signal intensity was adjusted to normalize signal intensity between *sfh8* and WT. Scale bars, 10 μm. Right: quantification of endosomes above the confocal diffraction limit (∼200 nm) in WT and *sfh8* under normal conditions (N=4 pooled experiments; n=123, ordinary ANOVA). **C.** Cleavage of SFH8 is important for robust PIN2 localization. α-PIN2 localization (midsection epidermis and cortex; 7 DAG, region 3) in WT,*sfh8,* and *sfh8* with SFH8^R84A^ (brightness has been adjusted here in sfh8 and *sfh8 SFH8^R84A^*), or HF-SFH8 (“F”). Yellow arrowheads denote PIN2 polarity. Right: PIN2 polarity index and cortex cells with proper polarity in WT, *sfh8* (expressing also *SFH8^R84A^* or *HF-SFH8*), *k135,* and *rsw4* (for the index: means±SD; N=10, n=10, “*”, p<0.0001; 1-way ANOVA, for the number of cells: N=4, n=118, Kruskal-Wallis). Scale bars, 5 μm. **D.** Proper PIN2 polar delivery but not endocytosis is mediated by SFH8. PIN2-GFP localization in WT and *sfh8* (epidermis and cortex; 7 DAG, region 3) treated with 50 μm BFA (±CHX) for 1 h (left) and after BFA washout for 30 min. CHX was added to a final concentration of 30 μM (1 h pre-treatment and retained throughout the experiment). Right: BFA bodies quantification in WT and *sfh8* (three independent pooled experiments; n=15, 5 fields of view; paired 2-tailed *t*-test between WT/SFH8 in the presence of BFA). Scale bars, 5 μm. **E.** FRAP micrographs from polarized PIN2 (region 3) in WT and *sfh8*. Note the offset of PIN2 polarity (yellow arrowheads) in *sfh8*. The rectangular denotes the bleached region of interest (ROI). Scale bars, 3 μm. **F.** PIN2 shows reduced delivery rate to the PM, as has been reported for the KISC mutants (Moschou et al, 2016). FRAP PIN2 signal recovery (normalized integrated density). Data are means ± SD (N=3, n=5). The red faded band parallel to the Y-axis indicates laser iteration time. Numbers next to the genotype, denote recovery half-time±SD (n=15, paired 2-tailed *t*-test).

Next, we asked whether SFH8 promotes the delivery or maintenance of polar domains. To this end, we used the drug brefeldin A (BFA) to induce intracellular agglomerates of PIN2 in so-called BFA bodies (aggregate of trans-Golgi network [TGN] and Golgi). We calculated the endocytosis rate of PIN2 to BFA bodies and the delivery rate from PIN2-positive BFA bodies back to the PM following BFA washout [36]. We further validated BFA experiments with FRAP to measure the rate of PIN2 delivery at the PM. Both assays confirmed that PIN2 delivery to the PM is compromised in *sfh8* and KISC mutants, while PIN2 endocytosis was not, as PIN2-positive BFA-bodies were produced at the same rate in the wild type and the *sfh8* mutant; these effects were independent of *de novo* PIN2 synthesis, as cycloheximide (CHX) treatments did not affect delivery rates or dissolution of the PIN2 endosomes in *sfh8* (**Fig 6D-F; S8A and B Figs**). Furthermore, SFH8 did not colocalize with clathrin and *sfh8* did not show defects in endocytosis of the endocytic tracer FM4-64 or the peptide PEP1 which is internalized by clathrin-mediated endocytosis (**S9 Fig**)[37], excluding the possibility that PIN2 removal from the PM is due to increased endocytosis.

To address the mechanism by which SFH8 might affect PIN2 delivery, we checked whether SFH8 stabilized PIN2 at the PM. In root region 3, SFH8-mScarlet (and KIN7.3), but not mScarlet-SFH8, co-localized with PIN2 and showed similar polarity (**Fig 7A-C**; PCC ∼0.9, and exclusion from lateral PM domains). On the contrary, PIN2-GFP and mScarlet-SFH8 PM puncta showed an anti-correlation of localization, excluding each other in regions 1 and 2, as observed in a super-resolution setting (**Fig 7D**; insets). Thus, SFH8^IDR^ properties in SFH8 clusters may reduce the delivery of proteins like PIN2 at the PM. To address whether the entropic bristle effect is responsible for this exclusion, we evaluated whether the formation of SFH8^ΔIDR^ and the transition to filaments might permit delivery of PIN2, which would likely be manifested as increased SFH8^ΔIDR^ (filaments) proximity to PIN2 (as the two proteins colocalize). Using a specific antibody against SFH8 (C terminus), we observed that SFH8 localization is not affected in a *pin2* mutant, suggesting that SFH8 affects PIN2 delivery, and not the other way around (**S10A Fig**). To test for interactions between PIN2 and SFH8 in root cells, we refined a quantitative proximity ligation assay (PLA; [38]). PLAs use complementary oligonucleotides fused to antibodies to determine the frequency with which proteins of interest find themselves nearby (**Fig 8A**). In PLA, we observed positive interactions between PIN2 and SFH8 in regions 3 and 4 (i.e., in the form of SFH8^ΔIDR^) in epidermal and cortex cells, using two different settings: i) in roots expressing *PIN2- GFP* (PLA antibody combination α-GFP/α-SFH8) or ii) in roots expressing *PIN2-GFP* and a C-terminally HF-tagged SFH8 (PLA antibody combination α-GFP/α-FLAG). We observed significantly lower PLA signals between PIN2 and SFH8 in the N-terminally tagged HF-SFH8 lines (α-PIN2/α-FLAG) as the full-length SFH8 showed anti-correlation of localization with PIN2. We detected no PLA signal for i) SFH8 and a PM aquaporin (PIP2a-GFP; α-GFP/α-SFH8), ii) in the *sfh8* mutant (α-PIN2/α-SFH8), and iii) in the vasculature where PIN2 is absent (**Figs 8B-E; S10B-D Figs**). PLA also confirmed that KIN7.3 interacts with SFH8 exclusively at the PM (**Fig S10C**; region 3 onwards), confirming the result from the rBiFC in **Fig 1**. To follow these results in a live imaging setting, we used Förster Resonance Energy Transfer (FRET) analyses, in which we detected high FRET efficiency for the SFH8-mScarlet/PIN2-GFP pair, indicative of interaction, in epidermal or cortex cells of regions 3 and 4 where SFH8 has been converted to filaments (**Fig 8F**). Using this FRET setting, we also verified the PM interaction between ESP and KIN7.3 in the same regions and cells (**S10E Fig**). Collectively, these results suggest that SFH8 filamentous conversion (SFH8^ΔIDR^) allows the association/delivery of proteins like PIN2 with the PM.

**Fig 7.**
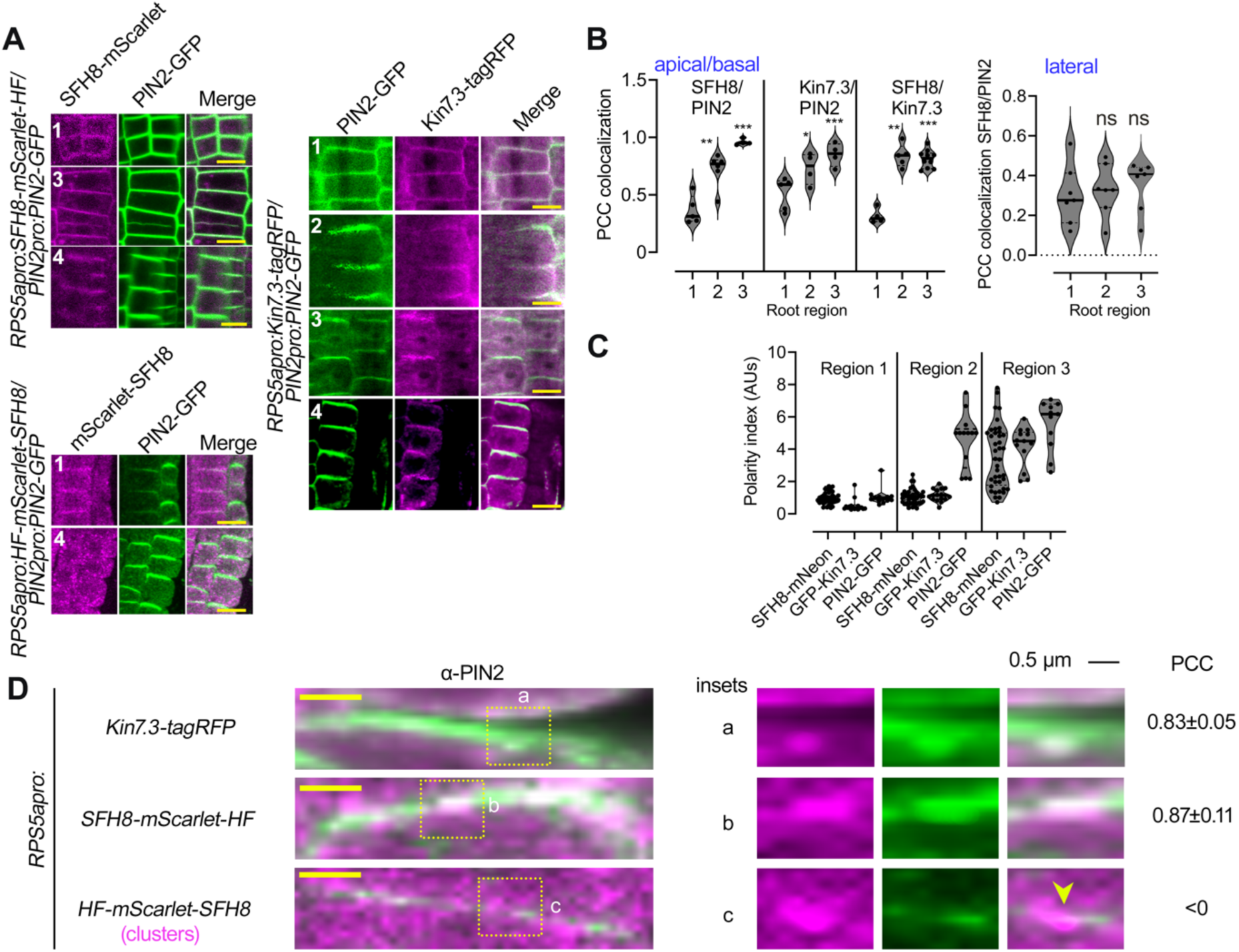
SFH8 restricts PIN2 delivery when it is uncleaved. **A.** Cleaved SFH8 (SFH8^ΔIDR^) can colocalize with PIN2 in regions 3 and 4. Representative images of co-localization of N- or C-terminally- tagged SFH8-mScarlet and HF-KIN7.3-RFP with PIN2- GFP in regions 1-4. Note the lack of colocalization between PIN2/mScarlet-SFH8 in region 4 (epidermal cells). Scale bars, 5 μm. **B.** Quantifications of PCC colocalization between PIN2 and SFH8 or KIN7.3 (apicobasal or lateral domains, for KIN7.3; N=3, n=36; p<0.05, *; p<0.001, **; p<0.0001, ***, nested 1-way ANOVA, compared to region 1). **C.** PIN2/KISC/SFH8 show similar polarity patterns at the PM. Polarity index for SFH8, KIN7.3, and PIN2 in three root regions (three independent pooled experiments; n=36, nested 1-way ANOVA). **D.** The N-terminus of SFH8 restricts PIN2/SFH8 colocalization. High speed (40 frames per second) and super-resolution (120 nm) confocal images with insets showing details of HF- mScarlet-SFH8 cluster/PIN2 exclusion. Note that the two proteins colocalized in non-clustered PM domains. PCC values represent colocalization analyses between filamentous patches of KIN7.3 or SFH8 with PIN2, while clusters of SFH8 (region 2: to observe the signal at the PM) showed anti-correlation (denoted also by the arrowhead). PCC±SD values are from three independent pooled experiments (n=120).

**Fig 8.**
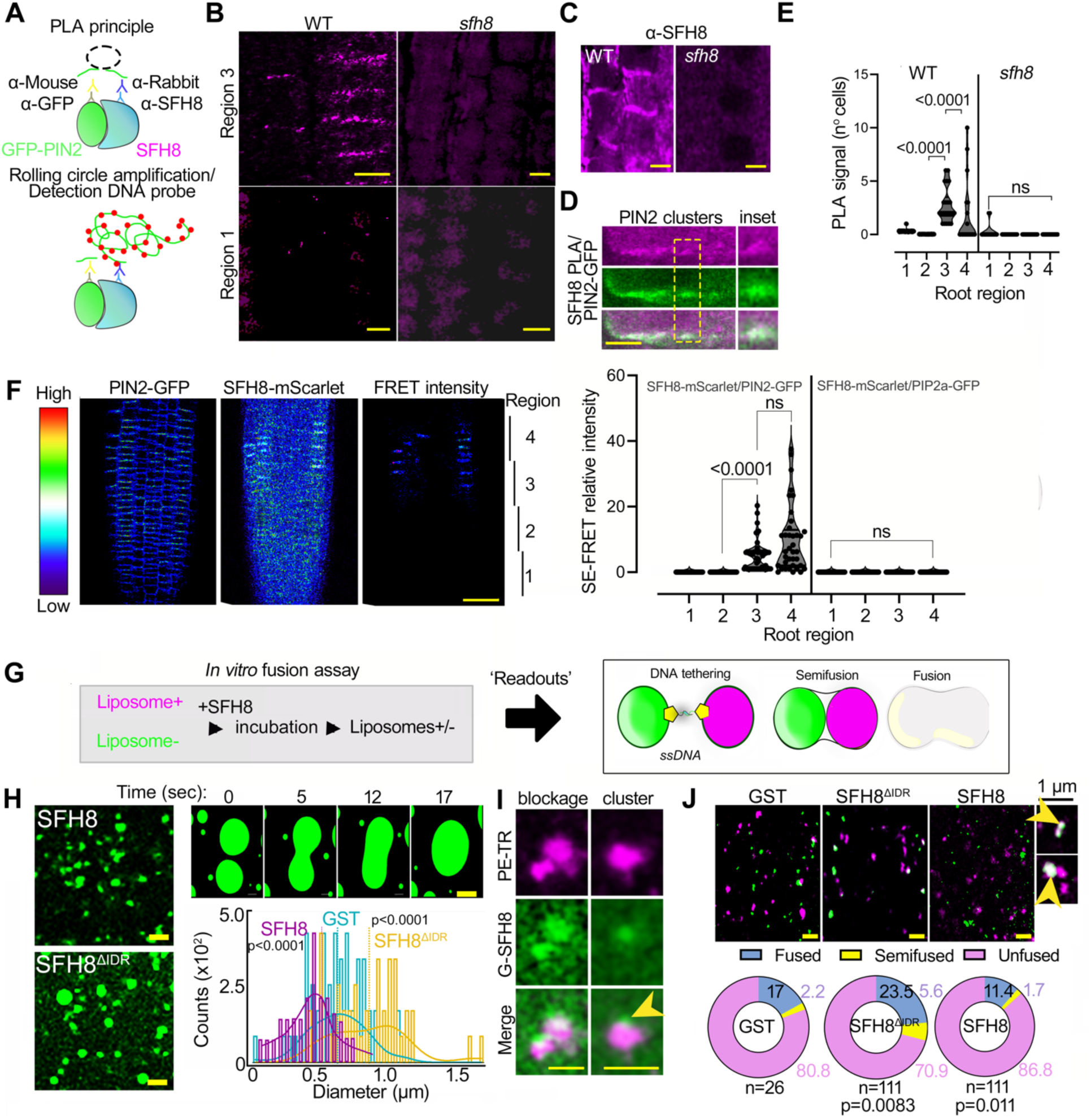
Filaments of SFH8 promote its interactions and establish more accessible interfaces at the PM. **A.** Proximity ligation (PLA) principle. **B.** Cleavage of SFH8 and removal of SFH8^IDR^ is indispensable for interactions with PIN2. SFH8 interacts with PIN2 in its truncated form (region 3 onwards). PLA-positive signal (resembling puncta) produced by the combination of α-SFH8/α-GFP in lines expressing *PIN2pro:PIN2-GFP* in WT or *sfh8*. Scale bars, 5 μm. **C.** Micrographs showing α-SFH8 signal in WT (“*sfh8*” is a negative control with only background signal). Note the lack of specific signalScale bars, 5 μm. **D.** Details of PLA-positive PIN2/SFH8 signal puncta at the PM. Scale bar, 1 μm. **E.** Quantification of PLA-positive cells in 4 regions (five independent experiments; n=20, nested 1-way ANOVA). **F.** SFH8 and PIN2 interact in living root cells as revealed by sensitized emission (SE)-FRET efficiency. PIN2-GFP (PIN2pro) and SFH8-mScarlet (RPS5apro) interact in epidermal and cortex root cells 5 DAG. Scale bar, 50 μm. Right: signal quantification of SE-FRET between the indicated combinations (five independent experiments; n=50, paired 2-tailed *t*-test). **G.** Establishment of a minimal system to detect effects of proteins in stereochemical hindrance. Pipeline of the DNA-zippers approach to test membrane fusion, as a stereochemical hindrance readout relevant to the function of SFH8. The DNA zippers bring together liposomes and promote their fusion. If SFH8 would exert stereochemical hindrance, liposome fusion mediated by complementary DNA zippers would be blocked. **H.** DNA-zipper assay with GST-SFH8^ΔIDR^ or -SFH8 (full length; lumen labeled with fluorescein only). The enlarged micrographs (top) show a time series of the tethering/fusion of two liposomes that converted to GUVs due to SFH8^ΔIDR^. Quantifications of fusions are also shown (three pooled independent experiments, with n=225; means indicated with vertical lines; Wilcoxon). Scale bars, 2 μm. **I.** The increased hydrodynamic radius of SFH8^IDR^ (entropic bristle effect) can restrict fusion and likely impose stereochemical hindrance. Fusion blockage by fluorescently labeled SFH8 clusters on liposomes (stained with PE-texas red [magenta]). Arrowhead denotes an SFH8 cluster formed atop the labeled liposome (images after deconvolution in a super-resolution setting with 120 nm resolution). Scale bars, 2 μm. **J.** Effect of stereochemical hindrance exerted by SFH8 on membranes. Liposome content mixing with lumen stained with fluorescein (green), and lipids stained with PE-texas red (magenta). Scale bars, 2 μm. Insets show a hemifusion event (upper), and a combination of hemifusion with a tethered LUV (lower inset). The arrows show content mixing (pseudo-colored white). Lower: distribution (%) of LUVs among different treatments (three pooled independent experiments; n as indicated, Dunnett for “fusion”).

We then asked whether the observed anti-correlation between SFH8 cluster signal and PIN2 might indeed imply stereochemical hindrance through the entropic bristle effect imposed by full-length SFH8 (by SFH8^IDR^) that could restrain delivery of proteins such as PIN2. We thus established an *in vitro* membrane fusion assay as a proxy of stereochemical hindrance at membranes based on cholesterol-modified DNA zippers (lipid-DNA-zippers; **Fig 8G**) [39]. To test SFH8-mediated membrane fusion, we used assays with low content of labeled phosphatidylethanolamine (PE)-Texas red (peripheral labeling) and fluorescein-containing large unilamellar vesicles (LUVs; 400 nm). Under our super-resolution settings, we resolved three events driven by DNA zippers: membrane tethering, hemifusion (lipid mixing), and fusion resulting in the unification of the lipid bilayer and the intermixing of the volumes. As SFH8^ΔIDR^ converted to filaments in a few minutes, to ascertain that observed effects were not due to differential binding of SFH8^ΔIDR^, we used quartz crystal microbalance with dissipation (QCD-M) to monitor the binding of LUVs in real-time (**S10F and G Fig**). QCD-M can reveal the interaction dynamics between lipids and the sensor surface, translating differential binding of proteins to bilayers in an observable real-time response. The detection is based on measurements of the acoustic ratio of dissipation *versus* frequency change (Δ*_D_*/Δ*_F_*), which depend on real-time mass changes [5], i.e., how much protein would bind on the lipid surface. In QCD-M, both SFH8 and SFH8^ΔIDR^ show similar affinities toward the LUVs, thus excluding differential binding as a potential driver of the changes in fusion dynamics. In the SFH8^ΔIDR^ samples, the average diameter of liposomes was ∼1 μm, in contrast to SFH8 (∼0.5 μm), which was below that of free GST samples (∼0.7 μm) (**Fig 8H**). Content mixing analyses showed that almost 30% of the SFH8^ΔIDR^ samples show semi-fused or fused LUVs (2-fold lower for full-length SFH8; **Fig 8H-J**), while fusion/hemifusion events with SFH8 were even less than those with GST. This result suggests that the N-terminal SDH8^IDR^ when on SFH8 exerts stereochemical repulsion blocking the delivery of proteins likely due to the increased hydrodynamic radius of the IDR. Hence, the uncleaved full-length SFH8 leads to a restriction of protein delivery at the PM, while SFH8^ΔIDR^ promotes delivery.

### SFH8 Cleavage by KISC Mediates Developmental Robustness

Next, we asked whether the KISC-SFH8 link affects development. At the seedling stage, the *sfh8* phenotypes resembled those of KISC mutants (*rsw4* and *k135*) showing both reduced root growth and gravitropism (**Fig 9A; S11A Fig**). As *sfh8-1* and *sfh8-2* were phenotypically similar, we carried out further analyses with *sfh8-1*. We did not observe additive phenotypes at early developmental stages in KISC/*SFH8* mutant combinations (*rsw4 sfh8* and *k135 sfh8*), suggesting some functional convergence between KISC and SFH8 (**Fig 9B**). Adult mutant plants also showed a shorter stature, decreased branching, and smaller cotyledons, or leaves (**S11A-C Figs**). Yet, *sfh8* produced shorter siliques and exhibited a more severe adult phenotype than the one reported for the KISC mutants (**S11A-C Figs**). In *sfh8*, the apical meristem length was ∼2-fold smaller than in the wild type, due to lower mitotic activity as defined by the Cell Cycle Tracking system [40] (**Fig 9C and D**). Although SFH8 is a SEC14-like protein and would therefore be expected to be involved in general lipid homeostasis, its loss of function (or of KISC) did not affect lipid levels at the PM (**S11D Fig**), suggesting that the observed phenotype does not relate to perturbations in general lipid homeostasis. All the constructs used herein encoding N- or C-terminally tagged SFH8 fusions (with tags or fluorophores) and driven by the promoters used in this work (*SFH8pro*, *RPS5apro*, or *KIN7.3pro*; **S11E and F Figs**) all rescued the *sfh8* phenotype.

**Fig. 9.**
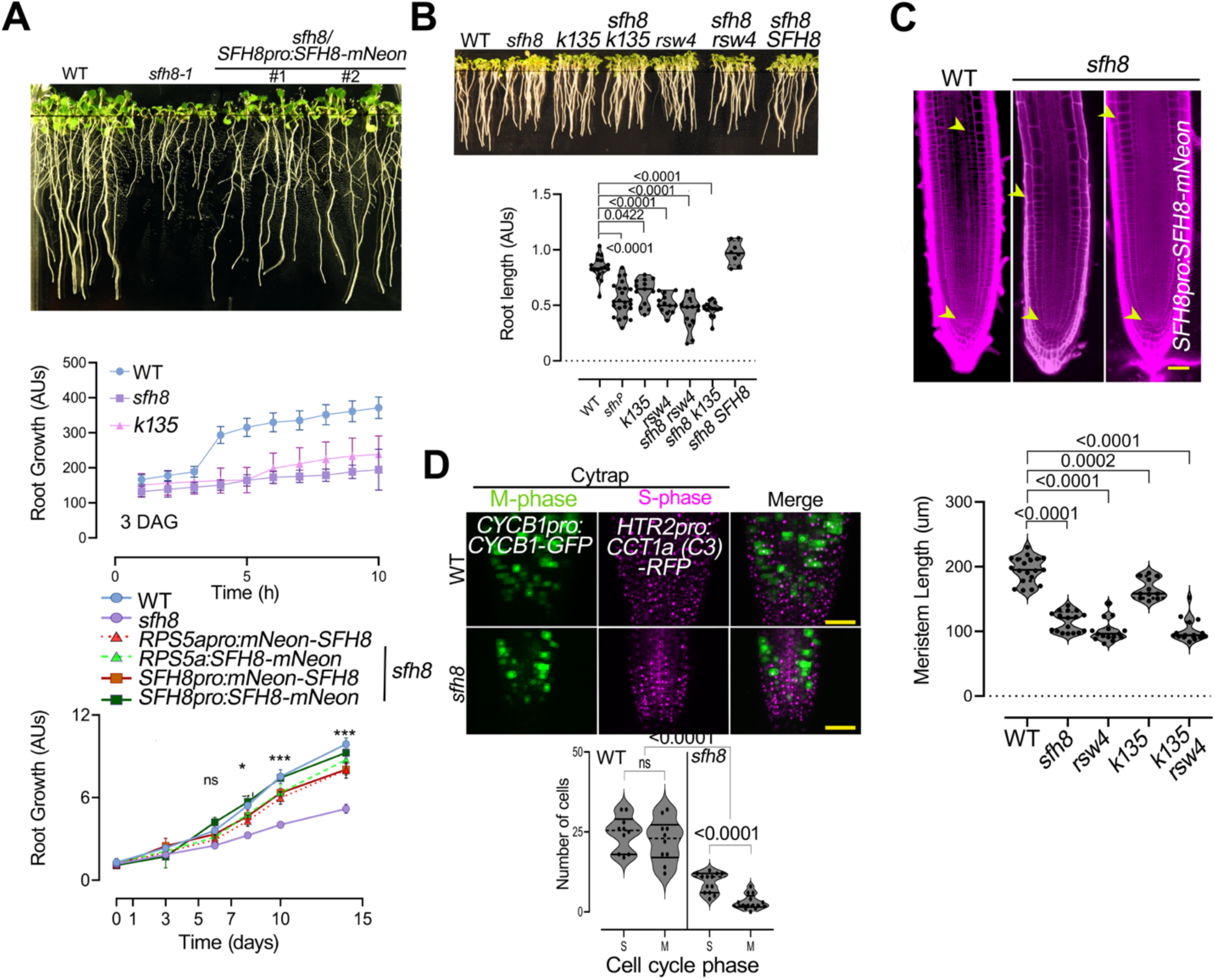
SFH8 modulates development. **A.** Phenotypes of young wild type (WT), *sfh8-1,* and rescued lines expressing *SFH8pro:SFH8- mNeon* (10 DAG on plates). Bottom: kinematic root growth quantification in the order of hours using “SPIRO” (see also Materials and methods; 3 DAG=time 0; at least five measurements (n≥25; 1-sided Dunnett), or in a time frame of days of WT, *sfh8*, *k135*, and rescued lines (with *RPS5a* or *SFH8* promoter, N- or C-terminally tagged). Asterisks: significance at p<0.05 (*) or p<0.001 (***); ns, no significant: WT vs. *sfh8*; *t*-test. **B.** Root length of WT, *sfh8, k135, rsw4*, the double *rsw4 sfh8* and *k135 sfh8* mutants, and the rescued line *sfh8 SFH8* (with *SFH8pro:SFH8-mNeon*, 7 DAG; three independent experiments, n=5 in each; paired *t*-test). **C.** The depletion of *sfh8* leads to a reduced meristem size. Root meristem micrographs (5 DAG; red signal: stained cell walls with propidium iodide); quantifications of WT, *sfh8*, or KISC mutants’ meristem sizes (n=19; ordinary ANOVA). Scale bars, 50 μm. **D.** The reduction of *sfh8* meristem size is due to compromised cell divisions. Micrograph of the “Cycle Tracking in Plants” marker (CyTRAP, tracking S and M phases in root cells) in WT and *sfh8* (7 DAG); quantification of S and M phase (three independent experiments, n=5 in each; paired *t*-test). Scale bars, 50 μm.

We further examined whether the phase transitions of SFH8 (cluster-to-filament) are important for phenotypes. The uncleavable SFH8 variant (SFH8^R84A^) which cannot form filaments failed to rescue *sfh8*, and the same was observed for SFH8^6KtoA^ (**S12 Fig**). On the other hand, deletion of the IDR (*SFH8^ΔIDR^*) also led to a significant loss of SFH8 polarity in roots, a lack of root developmental robustness, and only partial *sfh8* rescue (**S12 Fig**, graph; lack of *sfh8* complementation), suggesting that the initial clustering of SFH8 is functionally important perhaps to . Using the *R2D2* ratiometric auxin input and the *DR5* auxin output sensors (the R2D2 principle is detailed in **Supplemental Info**), we confirmed defects in *sfh8* and KISC mutants with occasional ectopic auxin input or output in distal meristem regions, consistent with perturbations in auxin signalling robustness (**Fig 10A and B**). Reverse transcription-quantitative PCR (RT-qPCR) analyses validated the reduced robustness of selected auxin-related genes in the root meristem of *sfh8* and *k135* mutants (**S13A Fig**). Using imaging, we confirmed that the least robust gene out of the ones tested, *YUCCA2*, shows a surprising variation in its expression levels in the *sfh8* mutant among root cells (**S13B-D Fig**). Overall, these data suggest that SFH8/KISC phase transitions affect developmental robustness.

**Fig 10.**
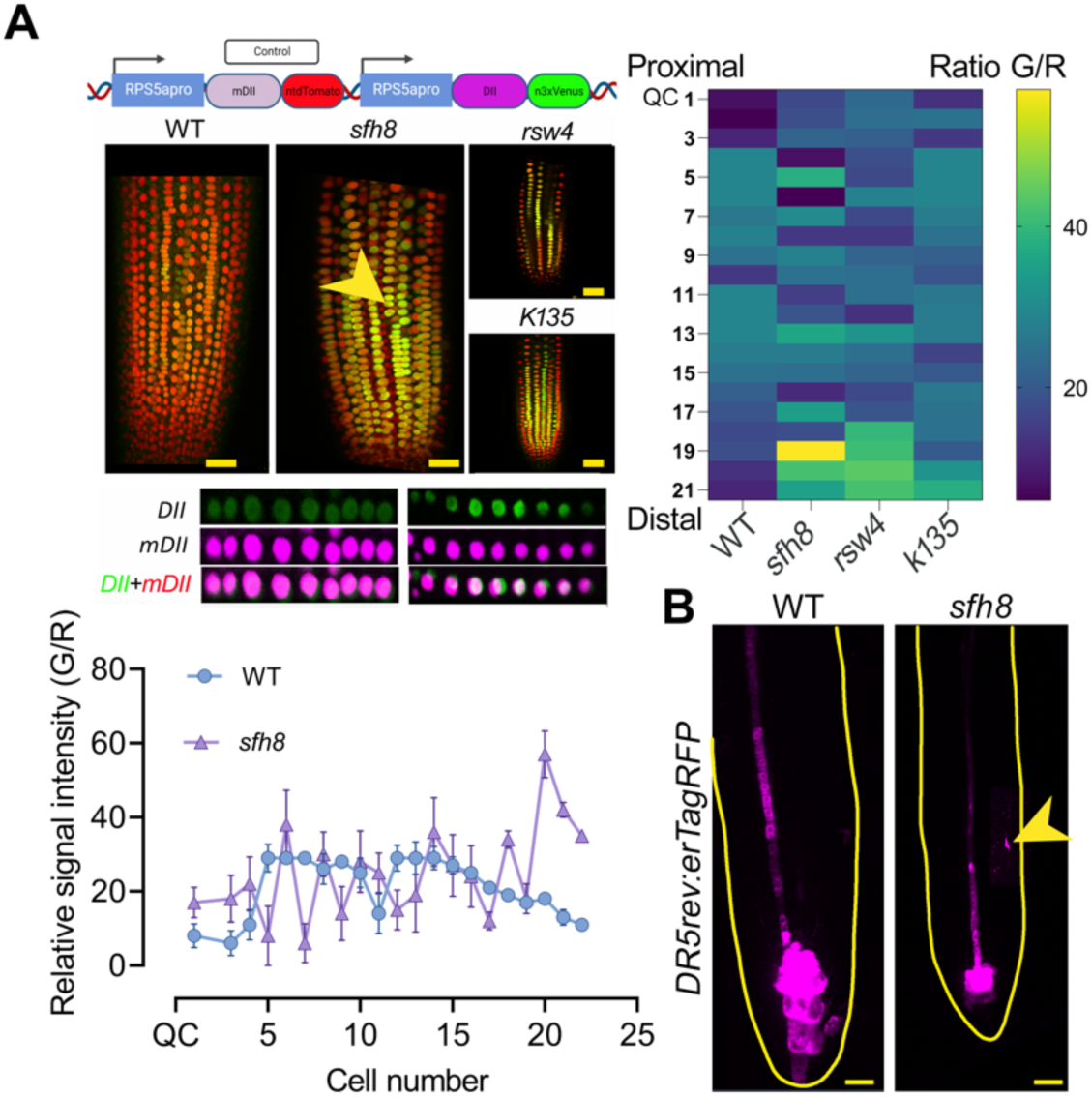
SFH8 can mediate partially the robustness of auxin signaling. **A.** SFH8 affects auxin signaling robustness inputs. R2D2 auxin input sensor (cartoon: transcriptional units *DII-Venus* (“DII”) and *mDII-Tm* (“mDII”)) in WT, *sfh8*, and KISC mutants. The arrowhead denotes a region with perturbed auxin input. Scale bars, 20 μm. Bottom: “DII” and “mDII” auxin input perturbations in *sfh8* (right). The chart shows DII/mDII ratio quantification (QC to 21^st^ cell towards the shoot; left), and the heat map shows DII/mDII ratio in WT, *sfh8*, *k135*, and *rsw4* (QC(1)-21^st^ cell). Data are mean ± SD from three different roots (n=21 cells per root). **B.** Auxin output sensor DR5 expression in WT and *sfh8*. Arrowhead denotes an auxin ectopic maximum in *sfh8*. Yellow outlines denote the roots. Scale bars, 20 μm.

## Discussion

Through KISC-SFH8 discovery, we identified a condensate that interfaces with plant membranes undergoing material properties switches. Collectively, the underlying mechanism comprises an array of events consisting of i) SFH8 LLPS at the PM, ii) reduction of SFH8 clustering and consequent interaction between KISC and SFH8, and iii) proteolytic cleavage of SFH8, which is accompanied by structural modifications that can allow interactions with other membrane proteins, e.g., PIN2. This module contributes to developmental robustness, at least partially through auxin, although we believe that this pathway has general functions in the regulation of PM domains. Intriguingly, SFH8 phase transition is induced by the highly conserved protease ESP, thereby expanding its known target range. So far, ESP targets have mainly been linked to functions in dividing cells. Apart from expanding the targets of ESP, this module also uncovers a novel way to regulate LLPS via limited proteolysis. We further speculate that the released N-terminal SFH8^IDR^ retains features of the LLPS SFH8 state even after its cleavage, suggesting a structural memory for condensates. In this direction, further work will reveal whether other proteolytic fragments can retain condensate memory which might be important for functions.

Condensation is crucial for polarity establishment in biological systems; yet, our work might appear counterintuitive, as it starts challenging or extending these models by showing that LLPS may simply be a mechanism for reducing polarized secretion, conflicting with observations in synapses where condensation exerts an opposite effect to those described here [1]. SFH8 liquid condensates prelude shifts in material properties (from LLPS to likely more solid filaments) that promote a functionality switch for SFH8: from blocker to fusion enhancer. Likewise, Dynamin-related proteins (DRPs) oligomerize into macromolecular ensembles to remodel membranes [41], while coronaviruses (e.g., SARS-CoV-2) co-opt host proteases for structural reconfiguration that prime fusogenic activity of SPIKE resulting in the fusion between the virus and its host cell [42, 43]. Upon IDR removal, SFH8^ΔIDR^ was stabilized, likely due to reduced intramolecular stereochemical repulsion and charge attenuation, which allowed filamentous state conversion and fusion at polar domains. Overall, this mechanism establishes an example of LLPS with importance for PM micropatterning that transcends to polarized patterns.

Furthermore, the dimensionality reduction caused by the PM binding of SFH8 promotes condensation. Cluster formation through LLPS could have a direct effect on PM properties. As has been recently shown, condensates can potentiate lipid clustering [44]. SFH8 could in turn affect lipid clustering, especially of PIs, with important roles in signaling during development. We believe that these functions may also be relevant during stress, given the important roles of PIs in transducing stress signals [45]. The microscale entropic bristle–like properties of SFH8^IDR^ when forming mesoscale condensates in the cytoplasm (the SFH8^IDR^-bodies) suggest that SFH8^IDR^ can restrict interactions with certain proteins when bound to the PM and can likely establish protein/vesicle depletion interfaces. We reconciled this proposition *in vitro* by showing that the removal of SFH8^IDR^ by ESP locally promoted attraction and fusion by alleviating steric hindrance. Alternatively, SFH8 clusters may engulf diffraction-limited vesicles and promote their fusion upon IDR removal. Because SFH8 is a SEC14-like protein, it may render lipids vulnerable to enzymatic modifications [46], regulating local lipid environments at the microscale or nanoscale. These modifications could promote vesicular fusion. SFH8^ΔIDR^ may also reduce the energy barrier required for fusion through an increase in fluidity by its filamentous structure, as has been shown for microtubule filaments [47], or the reduction of the entropic bristle effect.

Addressing how KISC and SFH8 function in development and perhaps stress is an important priority for our future research. The details of structural modifications at the PM for SFH8 upon the removal of the SFH8^IDR^ need further exploration. Intriguingly, as aforementioned, the IDR at the SFH8 N terminus is conserved throughout evolution (**File S2**), suggesting that proteins with similar features should exist in other eukaryotes. We speculate that SFH8 modifications may increase its activity Other pertinent questions are: How do SFH8 clusters form in the first place, and what is the function (if any) of the cytoplasmic “SFH8^IDR^-bodies” condensate? Our work thus provides insights relevant to condensates interfacing with membranes and describes liquid-like PM protein clusters and their transitions.

## Materials and methods

### Arabidopsis backgrounds and ecotypes

All the plant lines used in this study were in the Arabidopsis (*Arabidopsis thaliana*) Columbia-0 (Col-0) accession except the ones as indicated individually, and a detailed description can be found in Materials Table. Primers used for genotyping, RT-qPCR, and cloning can be found in **S1 Table**. The following mutants and transgenic lines used in this study were described previously: *rsw4* mutant [48], *k135* [20], *35Spro:mCherry-MAP4^MBD^*[20], *35Spro:smRS-GFP-TUB6*, *35Spro:smRS-GFP-TUA6* (Nottingham *Arabidopsis* Stock Center-NASC; N6550), *35Spro:GFP- TUB9* (NASC; M84706), *PIN1pro:PIN1-GFP* [49], *PIN2pro:PIN2-EGFP* [50]; *PIN3pro:PIN3- EGFP*, *PIN4pro:PIN4-EGFP* and *PIN7pro:PIN7-EGFP* [51], *DR5v2-ntdTomato / DR5-n3GFP* (R2D2) [52], Cytrap: *HTR2pro:CDT1a(C3)-RFP / CYCB1pro:CYCB1-GFP* [40]. *SNX1pro:SNX1- mRFP* [53], *TIP1pro:TIP1-GFP* [54], *35Spro:GFP-PIP2a*, *35Spro:GFP-AHA1* [55], *WAVE22*, *WAVE27* [56]. *DR5rev:erTagRFP / pTCS:nGFP* [57], In the Wassilewskija background: *CLC2pro:CLC2-EGFP* [58]. The following lines were ordered from GABI or NASC: *sfh8-1*: GABI_55IF03, *sfh8-2*: SALK_006862. Arabidopsis plants were transformed according to [59] using *Agrobacterium tumefaciens* strain GV3101. In all experiments, plants from T1 (co- localization experiments), T2, or T3 (for physiology experiments) generations were used. The Arabidopsis fluorescence marker lines were crossed with *sfh8-1* and corresponding SFH8 transgenic lines to avoid the gene expression levels differences caused by positional effects; F1 (co-localization experiments) and F2 were used in experiments.

### Plant growth condition***s***

Arabidopsis seedlings were surface-sterilized and germinated on ½ strength Murashige and Skoog (MS) agar medium under long-day conditions (16-h light/8-h dark) and were harvested, and treated, or examined as indicated in the context of each experiment. In all experiments involving the use of mutants or pharmacological treatments, the medium was supplemented with 1% (w/v) sucrose. Arabidopsis plants/lines for crosses, phenotyping of the above-ground part, and seed harvesting were grown on soil in a plant chamber at 22°C/19°C, 14 h/10 h light/dark cycle, and light intensity 150 µE m^-2^ s^-1^. *N. benthamiana* plants were grown in Aralab or Percival cabinets at 22°C (or 28-30°C for restrictive temperature treatments in the case of *rsw4*), 16 h/8 h light/dark cycle, and light intensity 150 µE m^-2^ s^-1^.

### Plant material, phenotypic analysis, and drug treatments

For quantification of phenotypes, seeds were surface-sterilized, plated on MS medium and seedlings were grown vertically. Customized Smart Plate Imaging Robot (SPIRO) imaging was done with 15-min intervals of fully automated imaging acquisition (https://www.alyonaminina.org/spiro), in a growth cabinet (Aralab). For a given genotype, differential contrast interference (DIC) images were captured on a Leica DM2500, Leica DM6B, or Leica DM6000. Arabidopsis Col-0 was used as the wild-type control. To define root length, images were captured of the plates using a Leica DM6000 with a motorized stage and computationally compiled together. Root length or size was determined using Image J/Fiji (National Institute of Health). For 1,6-hexanediol treatments, 10% (v/v) aqueous solution was used. XVE-driven expression was activated by transferring seedlings on plates containing various estradiol (in ethanol; light-tight tube aliquots) concentrations, which we determined experimentally in trial experiments (2-100 μM).

### Evolutionary relationships of taxa and sequences analyses

The evolutionary history of SFH8 was inferred using the Neighbor-Joining method [60]. The optimal tree is shown. The percentage of replicate trees in which the associated taxa clustered together in the bootstrap test (1,000 replicates) is shown next to the branches [61]. The evolutionary distances were computed using the Poisson correction method [62] and are in the units of the number of amino acid substitutions per site. This analysis involved 201 amino acid sequences. All positions with less than 70% site coverage were eliminated, i.e., fewer than 30% alignment gaps, missing data, and ambiguous bases were allowed at any position (partial deletion option). There was a total of 611 positions in the final dataset. Evolutionary analyses were conducted in MEGA X [63].

### Transient N. benthamiana assays

For transient assays, *Agrobacterium tumefaciens* strain GV3101 was used. Infiltrations were done as described previously [64]. During treatments, plants were kept in defined growth conditions in an Aralab growth chamber.

### Root gravitropism assays

To observe root gravitropism, seedlings were grown vertically on plates (½ strength MS agar medium under long-day conditions (16-h light/-8h dark) for 12 DAG. The root tip angle change was measured using Fiji.

### Cloning and plasmids

Primer sequences used for amplicons are listed in **S1 Table**. Cloning was performed either by Gateway, restriction enzyme digestion, or In-fusion (Takara). The following constructs were produced in i) pENTR vectors were generated via BP reaction with pDONR/Zeo (Invitrogen) and PCR products: SFH genes (amplicons from RT-PCR using cDNA from 1-week-old seedlings), truncations of SFH8 (amplicons from pDONR/Zeo-SFH8 was used as a template), mutations of SFH8 by mutagenesis PCR using pDONR/Zeo-SFH8. The KIN7.3 and truncations were previously described [20]; ii) In pGWB601: *SFH8pro:mNeon-gSFH8* (g, for genomic), *RPS5apro:mNeon-gSFH8*, *KIN7.3pro:mNeon-gSFH8*, *SFH8pro:gSFH8-mNeon*, *RPS5apro:gSFH8-mNeon, KIN7.3pro:gSFH8-mNeon, RPS5apro:HF-mScarlet-gSFH8-mNeon, RPS5apro:HF-gSFH8, RPS5apro:gSFH8-HF, KIN7.3pro:CFP-cKin7.3-mNeon.* The pGWB601 empty vector was used as a backbone, cut open by XhoI and SacI, then the vectors assembled in four or five fragments using In-fusion cloning (amplicons from Arabidopsis genomic DNA, template for *mNeon*, *mScarlet*, and *KIN7.3pro:CFP-cKIN7.3)*; iii) pGBKT7/pGADT7-gateway-compatible [65] for Y2H: LR reaction with the pENTR vectors of KIN7.3 and truncations, SFH8, SFH8 truncations, and mutations; iv) ratiometric bimolecular fluorescence complementation (rBiFC)-gateway-compatible system [25]: KIN7.3 (pDONR/P3P2-KIN7.3) and SFH8 (pDONR/P1P4-SFH8); v) RPS5apro gateway compatible dual tagged vectors: modified pGWB517 and 560 empty vectors were used as a backbone, cut open by HindIII and XbaI, and then the vectors assembled by two fragments In-fusion cloning (amplicons from Arabidopsis genomic DNA 1.6 kb promoter of *RPS5a* and template for *FLAG* and *sGFP*). The *KIN7.3* (pDONR/Zeo-KIN7.3) and *SFH8* (pDONR/Zeo-SFH8) clones were used to generate the FRET pair and the biosensor, respectively; vi) pGAT4 and pDEST15 gateway compatible vectors for His- and GST-protein production in *Escherichia coli*: full-length or truncations of *KIN7.3* and *SFH8* pDONR/Zeo vectors were used; vii) inducible constructs under the KIN7.3 promoter (XVE): vector was cut with PmeI and MluI and then the vectors assembled by two fragment In-fusion cloning (amplicons from Arabidopsis genomic DNA 1.7 kb promoter of *KIN7.3* and part of LexA); inserts from pENTR-GFP-ESP and pENTR-HA-KIN7.3 tail were introduced by LR reaction from the corresponding pDONR/Zeo vectors; viii) The pDR gateway compatible vector for yeast temperature-sensitive complementation: LR reaction with pDONR/Zeo-SFH8 and the indicated truncations (in pDONR/Zeo). The SYN4 cloning has been described in [29].

### Yeast two-hybrid screening and interactions

The genotype of the strain Y2HGold is MATa, trp1-901, leu2-3, 112, ura3-52, his3-200, gal4Δ, gal80Δ, LYS2::GAL1UAS-Gal1TATA-His3, GAL2UAS-Gal2TATA-Ade2, URA3::MEL1UAS-Mel1TATA-AUR1-C MEL1. The genotype of the strain Y187 is MATα, ura3-52, his3-200, ade2- 101, trp1-901, leu2-3, 112, gal4Δ, gal80Δ, URA3::GAL1UAS-GAL1TATA-lacZ. Transformed yeast cells were incubated at 30°C until OD600=0.8 in a minimal medium (SD) lacking the amino acid tryptophan. To confirm the expression of the baits, the total protein (10 µg) was extracted using alkaline lysis and resolved by SDS-PAGE. Fusion proteins were detected with α-Myc monoclonal antibodies (Roche, Stockholm, Sweden). The following constructs were used: pBKGT7-AtESP domain A (DomA; [20]; pBKGT7-AtESP Domain C (DomC; pBKGT7-AtMau2; [29] pBKGT7-cKIN7.3; pBKGT7-cKIN7.3-motor; pBKGT7-cKIN7.3-tail [20]; The absence of self-activation was verified by a transformation of the baits alone to select on minimal medium (SD) lacking the amino acids leucine, histidine, and adenine. The baits were transformed into the strain Y2HGold and mated with the Universal Arabidopsis cDNA Library (Clontech) in Y187. For pairwise Y2H assays, the Gateway-compatible pGADT7 vector [65] and the yeast Y187 were used, including the following constructs: pGADKT7-SCC2 [29]; pGADKT7-cKIN7.3-tail; pGADKT7-cSFH8; pGADKT7-cSFH8-SEC14 domain (SD); pGADKT7-cSFH8-Nodulin; pGADKT7-cSFH8-Nodulin^6Kto^ ^A^ (or other indicated variants). The “c” corresponds to cDNA.

### Protein production and purification

The pGAT4/PDEST15 constructs were transformed in BL21 (DE3) Rosetta or BL21 (DE3) Rosetta II *Escherichia coli* cells. Bacterial cultures were grown in 800 mL of LB supplemented with 100 mg L^-1^ of ampicillin and 25 mg L^-1^ of chloramphenicol. Protein production was induced at OD_600_=0.5 with 0.05 to 1 mM IPTG (Isopropyl ß-D-1-thiogalactopyranoside). After 3 h the cells were harvested by centrifugation at 2,500g for 20 min at room temperature and frozen overnight at –80°C. Preparation of his-tagged recombinant proteins was performed according to manufacturer instructions (Qiagen). Preparation of GST-tagged recombinant proteins was performed according to manufacturer instructions, using Sepharose beads (GE Healthcare Life Sciences), while the pH of purification was 8.3 (otherwise the SFH8 proteins precipitated). Expression levels of proteins were estimated by CBB staining in PAGE or by Western blots. The proteins were dialyzed overnight in assay buffers (2 L).

### Protein immunopurification

*Agrobacterium tumefaciens* strain GV3101 carrying constructs expressing various forms of KIN7.3, SYN4, SFH8, ESP, or TAP-GFP were infiltrated into *N. benthamiana* leaves. Three-four days later, leaves were ground in liquid nitrogen and resuspended in ten volumes of buffer A (50 mM Tris-HCl [pH 7.5], 5% [v/v] glycerol, 10% [v/v] Ficoll, 0.1% [v/v] Triton X-100, 300 mM NaCl, 5 mM MgCl2, 1 mM EDTA, 1 mM EGTA, plant-specific protease inhibitor cocktail [Sigma], phosphatase inhibitors [Roche], and 1 mM PMSF) and centrifuged at 14,000 x g for 20 min at 4 °C. The supernatant was filtered through four layers of Miracloth (Calbiochem). For TAP-ESP capture, samples were mixed with immunoglobulin G beads and incubated at 4°C for 1 h with gentle rotation. Beads were precipitated by centrifugation at 300 g, washed three times with buffer A, and treated for 4 h with PreScission protease (GE Healthcare). The TAP-ESP beads were used directly, or the supernatant was incubated for 30 min with nickel beads and h-AtESP was eluted with 250 mM imidazole containing buffer A. Protein was dialyzed against 0.1 M PIPES (pH 6.8), 5 mM EGTA, 2 mM MgCl_2_, and 20% (v/v) glycerol buffer. Protein levels were estimated by Western blot.

### SFH8 in vitro cleavage assay

GFP-ESP was extracted from infiltrated *N. benthamiana* leaves as above using GFP-TRAP (CromoTek). GFP-ESP protein lysates were mixed with GST beads carrying GST-SFH8 or the corresponding mutants. The samples were left agitating at 37°C for 1-2 h, resolved by SDS-PAGE, and detected by immunoblot.

### Preparation of liposomes

1,2-dioleoyl-sn-glycerol-3-phosphocholine (DOPC), 1,2-dioleoyl-sn-glycerol-3-phospho-L-serine (DOPS), 1,2-dioleoyl-sn-glycerol-3-phospho-(1’-myo-inositol-4’,5’-bisphosphate) (PI(4,5)P_2_) and 1,2-dioleoyl-sn-glycerol-3-[(N-(5-amino-1-carboxypentyl)iminodiacetic acid) succinyl] (DGS-NTA(Ni)) lipids were purchased from Avanti Polar Lipids (Alabaster, Al, USA). Dipalmitoyl phosphatidylinositol 3-phosphate (PI3P) lipids were from Echelon Biosciences Incorporated (EBI). Lyophilized lipids were dissolved in 1:1 (v/v) chloroform: methanol mixed in glass flasks in the desired amount. The organic solvent was first evaporated under a gentle stream of nitrogen while gently turning the flask to form a thin lipid film onto the wall of the flask. The lipid film was further dried with nitrogen for at least 30 min. Lipid films were hydrated in a buffer containing 10 mM Tris-HCl, pH 7.5 (Merck), and 150 mM NaCl at a final concentration of 2 mg/ml. After vortexing for 30 min, the resulting multilamellar vesicle solutions were extruded 25 times through a polycarbonate membrane with 50 or 200 nm nominal pore diameter (LiposoFast, Avestin) leading to a stock solution of unilamellar vesicles which were stored at 4°C and used for 5 d at most.

### Protein labeling for LLPS

The dyes used for labeling were Alexa Fluor™ 647 C2 Maleimide and Alexa Fluor™ 555 C2 Maleimide (ThermoFischer Scientific). The dyes were dissolved in water-free DMSO (dimethyl sulfoxide) at a concentration of 1 mM. After the protein isolation with GST-affinity chromatography, the proteins were dialyzed against 20 mM Tris-HCl pH 7.5, 150 mM NaCl. Proteins were partially labeled in solution in a ratio of 1 mol protein to 0.1 mol (1:10 molar ratio labeled/unlabeled) of dye for 15 min on ice. The labeled proteins were kept at 4°C and used within the next 3 d.

### Preparation of surfaces for suspended liposomes

Before the experiments, Au-coated AT-cut crystals (QSX-301 Biolin Scientific, Q-Sense, Sweden) were immersed in 2% (v/v) Hellmanex solution and rinsed with double distilled water. The gold sensors were further dried using nitrogen flow and cleaned using UV-ozone cleaner (E511, Ossila, Sheffield, UK) for 30 min. The QCM-D experiments were performed using the Q-Sense E4 (Biolin, Q-Sense, Sweden) instrument and AT-cut quartz disks (5MHz). All the experiments were performed in a buffer solution under a constant flow rate of 50 μL/min at 25°C. Briefly, the gold surface was equilibrated with buffer; followed by neutravidin adsorption (200 μL, 0.2 mg/mL protein solution). 5’-biotinylated, 3’-cholesterol modified DNA was then used (200 μL, 0.075 pmol/μL) as a binding anchor to liposomes of different lipid compositions as previously described [66]. All liposome solutions were used at a concentration of 0.2 mg/ml and a final volume of 70 0μL. Finally, protein solutions of various concentrations were added at a volume of 500 μL.

### Supported lipid bilayer with DGS-NTA(Ni)

Supported lipid bilayers containing 1% (v/v) DGS-NTA(Ni) were formed on SiO_2_ coated sensors (Q-Sense QSX 303) upon the addition of 0.05 mg/mL lipid solution. Final frequency and dissipation changes were Δf=–170 Hz and ΔD = 0.13 x 10^-6^. These values are typical of the formation of a homogeneous supported lipid bilayer.

### Liposome-protein binding assays

For the liposome-protein pull-down assays, we used 25 μl of Small Unilamellar Vesicles (SUVs, 2 mg/ml) combined with approximately 500 ng of the protein of interest. Before binding we centrifuged the proteins at 16,000g, at 4°C, for 5 min, to avoid precipitates [67]. Twenty-five μl of protein mix with 25 μL liposome suspension were combined and incubated on an orbital shaker platform for 30-45 min at room temperature. The binding buffer consisted of 150 mM KCl, 25 mM Tris–HCl pH 7.5, 1 mM DTT, 0.5 mM EDTA. The samples were centrifuged at 16,000g for 30 min at room temperature, we harvested the supernatant (we added 16.7 μL of 4× Laemmli sample buffer to a supernatant fraction) and resuspended the liposome pellet in 300-500 μl 1× binding buffer. The centrifugation was repeated in the same conditions and we resuspended the liposome-protein pellet in 33 μL 4x Laemmli buffer. All samples were incubated at 95°C for 5 min and loaded 10 μL on SDS-PAGE for immunoblot.

### Liposome DNA-mediated fusion assay and SUPER template setting

To establish DNA-zippers, we adjusted concentrations of oligomers-cholesterol and used various temperatures (4-25°C) to achieve DNA-zipper-driven fusion at a percentage lower than ∼10% (noise). Eventually, all liposomes irrespective of the conditions used, given enough time fused to a percentage of ∼50%. The liposomes used were LUVs of 400 nm diameter, with the following composition: DOPC 71.5%, DOPS 27.5%, PE-Texas Red conjugated 0.5% and PI(4,5)P_2_ 0.5% (PS was included for SUPER template experiment to provide required membrane flexibility). We also used LUVs without PE-Texas Red lipid. Instead, they carried the fluorophore fluorescein (cf, 100 mM fluorescein in Tris-HCl pH 7.5). For the fusion of the two different liposome populations, we incubated each with a DNA primer for 45 min at room temperature. The sequence of the single-stranded DNA molecules can be found in **S1 Table**. For every liposome, we used 100 DNA molecules [68]. SUPER templates were prepared, as described previously [33]. Silica beads (monodispersed 4.9 μm in diameter; Corpuscular Inc., Cold Spring, NY) were added to a premixed solution of liposomes in LLPS buffer in a total volume of 100 μL in a 1.5-mL clear polypropylene centrifuge tube. Glass coverslips (Fisher Scientific, Pittsburgh, PA) were cleaned with piranha solution (concentration H_2_SO_4_/30% H_2_O_2_ 4:1, v/v) for 1 h at room temperature and washed extensively with boiling water. The coverslips were allowed to attain room temperature, stored under water, and air-dried before use. To coat the coverslips with PEG-silane, piranha-cleaned coverslips were first dried in an oven at 200°C for 3 h and then treated at room temperature for 30 min with PEG-silane (2%, [v/v]; Gelest Inc., Morrisville, PA) in acetone. The coverslips were later washed extensively with water and air-dried before use. The samples were observed with confocal LIGHTNING SP8 module (high speed and resolution of 120 nm) using an observation chamber and an Apochromat 63x objective (N.A.=1.4). images were deconvoluted by Leica built-in software.

### Phase separation assays

The slides and coverslips used for confocal microscopy were treated overnight with 1-2 mg/mL PLL-PEG, after cleaning with 2% (v/v) Hellmanex for approximately 2 h, at room temperature. The LLPS buffer used was the following: 10 mM HEPES (4-(2-hydroxyethyl)-1-piperazine ethane sulfonic acid), 150 mM NaCl, 0.1 mM EDTA, 2 mM DTT, pH 7.4 with and without 10% (w/v) polyethylene glycol (PEG 3000) as a crowding agent. The amount of the added labeled protein was 1-2 μM for 3 h at room temperature. Some of our samples contained Large Unilamellar Vesicles (LUVs) consisting of DOPC 0.5%: DOPS 99%: DOPE-Texas Red 0.5%. Phase separation was imaged via confocal microscopy.

### Whole-mount proximity ligation assay (PLA) and Immunocytochemistry

Four to five-day-old seedlings were fixed and permeabilized as described [20]. The anti-SFH8 antibody was raised against the unique peptide sequence 517-GNAIELGSNGEGVKEECRPPSPVPDLTET-545 in rabbits using standard immunogenic procedures. Primary antibody combination 1:200 for α-GFP mouse [Sigma-Aldrich, SAB2702197] and 1:100 for α-SFH8 rabbit, 1:100 for α-mNeon mouse [Chromotek, 32F6], and 1:100 for α- SFH8 rabbit or 1:200 for α-FLAG mouse [Sigma-Aldrich, F1804] and 1:200 for α-GFP rabbit [Millipore, AB10145]) were used for overnight incubation at 4°C. Roots were then washed with MT-stabilizing buffer (MTSB: 50 mM PIPES, 5 mM EGTA, 2 mM MgSO_4_, 0.1% [v/v] Triton X-100) and incubated at 37°C for 3 h either with α-mouse plus and α-rabbit minus for PLA assay (Sigma-Aldrich, Duolink). PLA samples were then washed with MTSB and incubated for 3 h at 37°C with ligase solution. Roots were then washed 2x with buffer A (Sigma-Aldrich, Duolink) and treated for 4 h at 37°C in a polymerase solution containing fluorescent nucleotides as described (Sigma-Aldrich, Duolink). Samples were then washed 2x with buffer B (Sigma-Aldrich, Duolink), with 1% (v/v) buffer B for another 5 min, and then the specimens were mounted in Vectashield (Vector Laboratories) medium.

Immunocytochemistry was done as described previously [20]. The primary antibodies used were rabbit α-SFH8 (diluted 1:500), rat α-tubulin YL1/2 (1:200; Santa Cruz Biotechnology), mouse α-FLAG (1:250), and sheep α-PIN2 (1:500). Specimens were washed three times for 90 min in PBST and incubated overnight with donkey α-sheep conjugated Alexa fluor 488, goat α-mouse tetramethylrhodamine isothiocyanate (TRITC), α-rat TRITC, and α-rabbit fluorescein isothiocyanate-conjugated (FITC) secondary antibodies diluted 1:200-250. After washing 3x in PBSΤ, specimens were mounted in Vectashield (Vector Laboratories) medium.

### Ratiometric BiFC and protoplast transformation

The rBiFC assay was done in Arabidopsis root protoplasts via PEG transformation as described previously [69]. After a 16-h incubation following transformation, protoplasts were observed with a Zeiss LSM780 laser scanning confocal microscope using 40×/1.2 W C-Apochromat in multi-track channel mode. Excitation wavelengths and emission filters were 514 nm/band-pass 530-550 nm for YFP, and 561 nm/band-pass 600-630 nm for RFP. Images were processed by Fiji.

### Quantification of fluorescent intensity, FRAP, and FRET

To create the most comparable lines to measure the fluorescence intensity of reporters in multiple mutant backgrounds, we crossed homozygous mutant bearing the marker with either a WT plant (outcross to yield progeny heterozygous for the recessive mutant alleles and the reporter) or crossed to a mutant only plant (backcross to yield progeny homozygous for the recessive mutant alleles and heterozygous for the reporter). Fluorescence was measured as a mean integrated density in regions of interest (ROIs) with the subtraction of the background (a proximal region that was unbleached and had less signal intensity than the signal of the ROI region). FRAP mode of Zeiss 780 ZEN software was set up for the acquisition of 3 pre-bleach images, one bleach scan, and 96 post-bleach scans (or more). Bleaching was performed using 488, 514, and 561 nm laser lines at 100% transmittance and 20-40 iterations depending on the region and the axial resolution (iterations increased in deeper tissues to compensate for the increased light scattering). In FRAP the width of the bleached ROI was set at 2-10 µm. Pre- and post-bleach scans were at minimum possible laser power (0.8 % transmittance) for the 458 nm or 514 nm (4.7%) and 5% for 561 nm; 512 x 512 8-bit pixel format; pinhole of 181 μm (>2 Airy units) and zoom factor of 2.0. The background values were subtracted from the fluorescence recovery values, and the resulting values were normalized by the first post-bleach time point and divided by the maximum point set maximum intensity as 1. The objective used was a plan-apochromat 20x with NA=0.8 M27 (Zeiss).

GFP-AHA1, PIP2a-GFP, PIN2-GFP, mNeon-SFH8, and SFH8-mNeon fluorescence were detected using a water- or oil-corrected 40x objective. During analyses, the FRAP mode of Zeiss Lsm780 ZEN software was set up for the acquisition of one pre-bleach image, one bleach scan, and 40 or more post-bleach scans. The following settings were used for photobleaching: 10 to 40 iterations; 10 to 60 s per frame; 100% transmittance with the 458- to 514-nm laser lines of an argon laser. Pre- and post-bleach scans were at minimum possible laser power (1.4 to 20% transmittance) for the 488 nm and 0% for all other laser lines; 512x512 pixel format; and zoom factor of 5.1. The fluorescence intensity recovery values were determined, then the background values were subtracted from the fluorescence recovery values, and the resulting values were normalized against the first post bleach time point.

For FRET, either acceptor photobleaching (AB) or sensitized emission (SE) modules of the SP8 Leica confocal microscope were used, using standard modules and internal calibration. The acceptor photobleaching experiments were done as described previously [20], using the parameters for FRAP. Fluorescence was detected using a water- or oil-corrected 40x objective. For SE-FRET, The correction factors β, α, γ and δ were calculated with the donor- and acceptor-only reference samples, as described for TMK1-AHA1 interaction [70].

### TIRFM imaging and tracking analyses

TIRF microscopy images were acquired using MetaMorph software on an Olympus IX-81 microscope. The system was maintained at 37°C during imaging. A DV2 image splitter (MAG Biosystems) was used to separate GFP and RFP emission signals. Time-lapse movies were obtained at, 100-ms intervals. For MSD analysis, 30-s-long movies with 100-ms intervals and 200-ms exposure were used. Particle tracking was limited to the amount of time that PM remained in a single focal plane; the median track length was 2,000 frames, corresponding to 3.5 s of imaging. The tracking of particles was performed with the Mosaic suite of Fiji or NanoTrackJ/TrackMate, using the following typical parameters: radius 3 of fluorescence intensity, a link range of 1, cutoff of 0.1%, and a maximum displacement of 8 px, assuming Brownian dynamics.

### LC-MS/MS lipids and proteins analyses and overlay assays of lipids

Lipid overlay assay PI strips were blocked in PBST (0.1% Tween 20 [v/v]) supplemented with 3% (w/v) fatty acid-free BSA (Sigma-Aldrich) overnight at 4°C and then incubated with 0.5 μg/ml GST fusion proteins for 1 h at room temperature. After washes with PBST-BSA, the membranes were incubated with rabbit α-GST for 1 h at room temperature, followed by IRDye 800 goat anti-rabbit secondary antibody for 1 h.

Lipids were extracted from roots after inactivating the tissue with boiling water or from microsomes [71]. Lipid classes were purified by solid-phase extraction, and phospholipids, glycolipids, diacylglycerol, triacylglycerol, and total fatty acids were measured by quadrupole time of flight mass spectrometry (Q-TOF MS/MS) or gas chromatography–flame ionization detection (GC-FID), respectively, as previously described [72].

For proteomic analyses of FLAG-tagged SFH8 immunoprecipitates, samples were analyzed by LC-MS using Nano LC-MS/MS (Dionex Ultimate 3000 RLSCnano System) interfaced with Eclipse Tribrid mass spectrometer (ThermoFisher Scientific). Samples were loaded onto a fused silica trap column Acclaim PepMap 100, 75 μm x 2 cm (ThermoFisher Scientific). After washing for 5 min at 5 μL/min with 0.1% (v/v) trifluoroacetic acid (TFA), the trap column was brought in line with an analytical column (Nanoease MZ peptide BEH C18, 130 A, 1.7 μm, 75 μm x 250 mm, Waters) for LC-MS/MS. Peptides were fractionated at 300 nL/min using a segmented linear gradient 4-15% B in 30 min (where A: 0.2% formic acid, and B: 0.16% formic acid, 80% acetonitrile), 15-25% B in 40 min, 25-50%B in 44 min, and 50-90% B in 11 min. Solution B then returned at 4% for 5 min for the next run. The scan sequence began with an MS1 spectrum (Orbitrap analysis, resolution 120,000, scan range from M/Z 350–1600, automatic gain control (AGC) target 1E6, maximum injection time 100 ms). The top S (3 589 sec) and dynamic exclusion of 60 s were used for the selection of Parent ions for MS/MS. Parent masses were isolated in the quadrupole with an isolation window of 1.4 591 m/z, automatic gain control (AGC) target 1E5, and fragmented with higher-energy collisional dissociation with a normalized collision energy of 30%. The fragments were scanned in Orbitrap with a resolution of 30,000. The MS/MS scan range was determined by the charge state of the parent ion, but the lower limit was set at 100 amu.

### TAMRA-PEP1 and FM4-64 internalization assays

The peptide PEP1 (ATKVKAKQRGKEKVSSGRPGQHN) was labeled with 5′-801 carboxytetramethylrhodamine at the N-terminus (TAMRA-PEP1) with an HPLC purity of 95.24% and molecular weight of 2905.24 (EZBiolab). The peptide was dissolved in water to obtain 1 mM peptide stocks. Further dilutions were done with ½ MS medium. Five-day-old seedlings were dipped into 1 mL ½ MS medium containing 100 nM TAMRA-PEP1 for 10 s, washed five times, and kept in 24-well plates with ½ MS medium for 40 min. Epidermal cells at the meristematic zone were imaged with a Zeiss LSM710 inverted laser scanning confocal equipped with 40×/1.2 W C- Apochromat M27 objective at a zoom factor of 3.5. TAMRA-PEP1 was excited at 559 nm, and fluorescence emission was captured between 570-670 nm, GFP was excited at 488 nm, and fluorescence emission was captured between 500-540 nm. Image quantification was done using Fiji software. To this end, the PM of individual cells was selected with the brush tool with a size of 10 pixels as well as the intracellular space with the polygon selection tool. The average intensity of the top 100 highest pixels for both the plasma membrane and the intracellular space was used to obtain a ratio between intracellular and plasma membrane fluorescence. Five epidermal cells from five plants were quantified.

The FM4-64 experiments were as described in [73]. In brief pulse labeling with FM4-64 (2 μM FM464, Molecular Probes; made from a 2-mM stock in DMSO) was done for 5 min (time 0) and then analyzed for 18 min at room temperature in Zeiss Lsm 780 with excitation at 488 nm and fluorescence emission captured between 540-670 nm. After the pulse with FM4-64, the roots were washed two times in an ice-cold MS medium. Quantifications were done as described for the TAMRA-PEP1 experiments above.

### RNA extraction, RNA-seq, and quantitative RT-PCR analysis

Total RNA from the seedlings was extracted using the RNeasy Plant Mini Kit with Dnase I digestion (QIAGEN). Reverse transcription was carried out with 500 ng of total RNA using the iScript cDNA synthesis kit (Bio-Rad) according to the manufacturer’s protocol. Quantitative PCR with gene-specific primers was performed with the SsoAdvanced SYBR Green Supermix (Bio- Rad) on a CFX96 Real-Time PCR detection system (BioRad). Signals were normalized to the reference genes *ACTIN2* using the ΔCt method and the relative expression of a target gene was calculated from the ratio of test samples to Col-0. For each genotype, three biological replicates were assayed in three qPCR replicates. RT-qPCR primers were designed using QuantPrime (www.quantprime.de) [74]. Primer sequences used for RT-qPCR can be found in **S1 Table**.

### Super-resolution imaging and negative staining

Confocal SP8 Leica confocal microscope LIGHTNING module was used for super-resolution on-the-fly imaging. Confocal images were obtained when indicated at maximum scanning speed (40 frames per second) using a 63x water immersion objective with a theoretical x/y-axial resolution of 120 nm upon deconvolution. Post-acquisition, images were deconvoluted using the LIGHTNING algorithm and water correction was set in the algorithm as the mounting medium. Imaging was done at room temperature in an inverted microscope setting. Negative staining was essentially done as described for the lipoproteins [75].

### Quantification and statistical analysis

Statistical analysis was performed in R studio (R-project.org) or GraphPad Prism (version 9.2.0). Each data set was tested for normal distribution (Gaussian) by the Shapiro-Wilk, D’ Agostino-Pearson, Anderson-Darling, and Kolmogorov-Smirnov normality tests. Lognormal versus Gaussian distributions were also evaluated (not used though herein). For tests involving pairwise comparisons, Student’s t-test, or Wilcoxon tests (or as indicated) were mainly used to define whether differences were statistically significant. The significance threshold was set at *p*<0.05 (significance claim-level alpha) and the exact values are shown in graphs (*p* values <0.0001 are indicated as such). For tests involving multiple comparisons, one-way ANOVA, or the Kruskal-Walli’s test (nonparametric analog of ANOVA) was used followed by Dunnet’s or Dunn’s multiple comparison test to define whether differences were statistically significant (or as indicated). Graphs were generated by using Microsoft Excel, R, or GraphPad Prism 9.2.0. Details of the statistical tests applied, including the choice of statistical method, and the exact number of “n” is indicated in the corresponding figure legend or directly on the graph. In violin plots, upper and lower dotted lines represent the first and third quantiles, respectively, horizontal lines mark the mean, and edges mark the highest and lowest values. Plots were depicted truncated or untruncated for aesthetic reasons. “N” corresponds to biological replicate and “n” to technical.

Image analyses and intensity measurements were done using Fiji v. 1.49 software (rsb.info.nih.gov/ij). The intensity of the fluorescence signal was measured as the Integrated Density in a region of interest. The dwell time rate of tagged proteins in FRAP experiments was calculated by the single exponential fit equations as described by [76]. Co-localization was analyzed using Pearson statistics (Spearman or Manders analyses produced similar results)[77]. Images were prepared by Adobe Photoshop v. 2021 (Adobe). Statistical analyses were performed with JMP v. 9 or 11 (www.jmp.com), GraphPad, or R. Curve fitting was done as we have described in detail for PCs [78]. Time series movies were compressed, corrected, and exported as .avi extension files. The unspecific fluorescence decay was corrected using Fiji v. 1.49 software and default options using the bleaching correction tool. Videos were digitally enhanced with Fiji-implemented filters, correcting noise using the Gaussian blur option and pixel width set to 1.0. Mean intensities in FRAP have been normalized (1 corresponds to pre-bleach signal intensity in arbitrary units).

## Author contributions

P.N.M, conceptualization, project administration, supervision, formal analysis, writing-original draft, funding acquisition, complex identification; C.L., A.M., F.P., A.P., P.R., L.C., investigation, formal analysis, data curation; S.B., P.D., Y.J., E.R., E.G., and G.S., methodology; F.P., and E.G., methodology and establishment of liposome assays, curation; C.L., P.N.M, and A.M., established advanced imaging approaches, visualization. All authors reviewed, edited, and approved the paper.

## Declaration of interests

The authors declare no competing interests.

## Supplemental materials

Figures S1-13 (legends below figures)

Movies S1-4:

1. TIRFM of a diffusive cluster following a directional track or non-directional track (part a and b)
2. Non-diffusive clusters with low dwelling times
3. Immobile patches of SFH8 atop short filaments resembling a “beads on a string” localization pattern
4. SFH8^IDR^ cytoplasmic puncta showing dynamic morphology with frequent splitting, fusion, and interconnections resembling liquid-liquid phase-separated (LLPS) condensates

Files S1-2

1. SFH-like proteins alignments
2. IDR region conservation in various SFH-like proteins Table S1: oligos and their sequences used in the study

Supplemental Information

See below

---

**S1 Fig.**
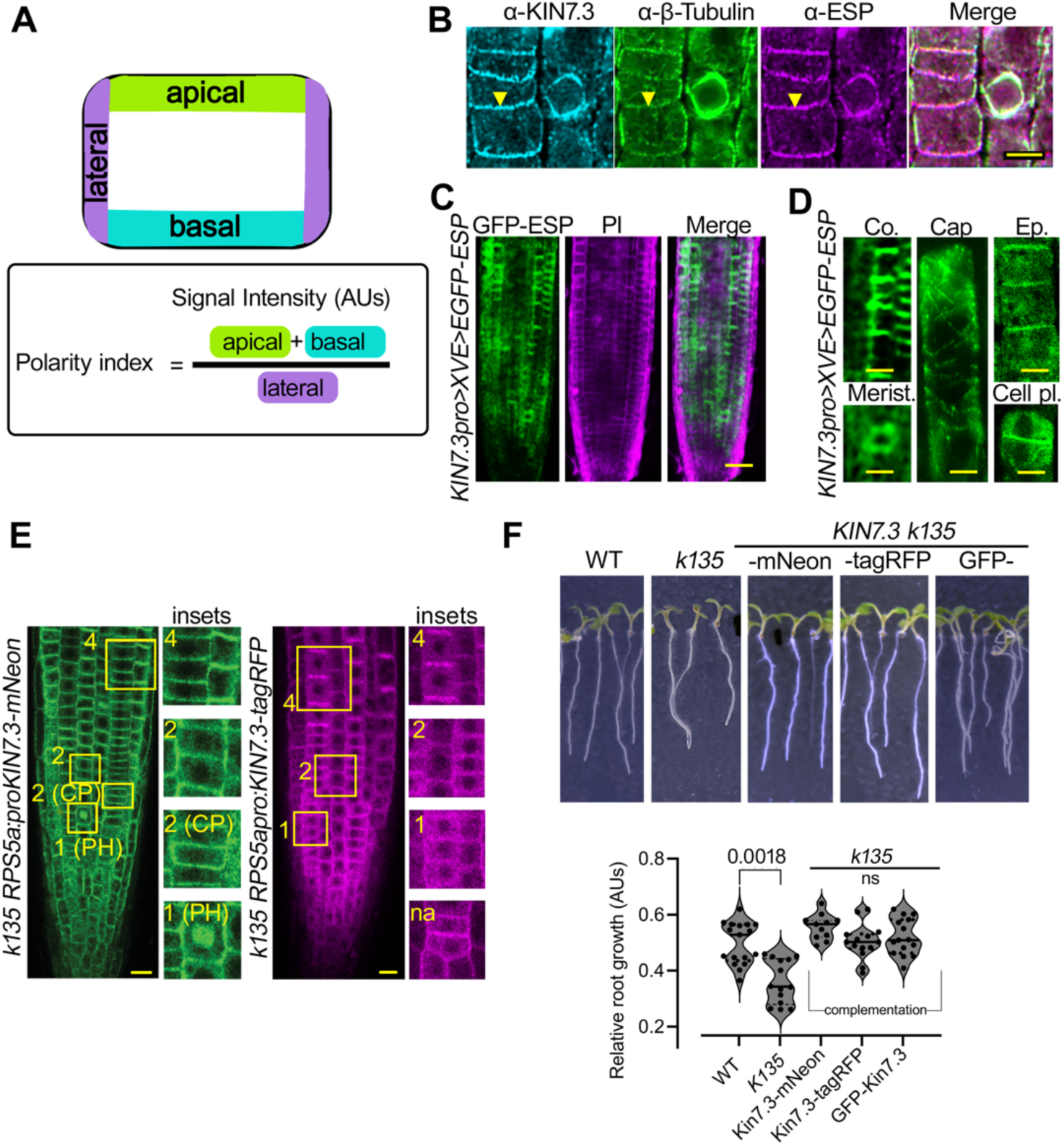
KISC can localize polarly at the PM. **A.** Apical/basal/lateral PM domain nomenclature and polarity index determination. **B.** ESP and KIN7.3 attain similar polarity patterns in root cortex cells. Representative confocal micrographs of α-ESP/α-KIN7.3 localizations at similar polar domains (counterstained with α-*β*- tubulin; region 4, experiment replicated multiple times). Scale bars, 5 μm. **C.** ESP localization in root epidermis or cortex cells. Representative confocal micrograph of the ESP estradiol inducible line (20 μM estradiol, 16 h), driven by the *KIN7.3* promoter (*KIN7.3pro>XVE>GFP-ESP*). *KIN7.3* promoter was used, as the *ESP* promoter was not functional Scale bars, 5 μm. ESP showed polarization only in the distal meristematic region, while in the proximal meristem region ESP was apolar. Images were obtained in low resolution (512x512 with minimal exposure and no averaging), due to photobleaching and low expression levels of ESP (experiment replicated multiple times with variable expression). Scale bar, 50 μm. **D.** Digitally zoomed-in micrographs (deconvoluted) from root tip cells of the *KIN7.3pro>XVEpro>GFP-ESP.* Merist., proximal meristematic cell (note the diffused cytoplasmic signal). Cap, lateral root cap. A cell plate decorated by ESP is also shown (bottom right). Note the MT-binding of ESP in the lateral root cap cell at which MTs are highly bundled (as also described in Moschou et al., 2013; 2016). **E.** KIN7.3 driven by the *RPS5a* promoter retains its localization in root cells. Representative confocal micrographs of 5-DAG seedlings expressing *RPS5apro:KIN7.3-mNeon* in the *k135* background at the indicated regions. Right: representative confocal micrographs of epidermal cells of root tip 5 DAG from seedlings expressing *RPS5apro:KIN7.3-tagRFP k135* (complementation line). CP, cell plate; PH, phragmoplast. Scale bars, 20 μm. **F.** KIN7.3 driven by the *RPS5a* promoter retains its functionality irrespective of the tag used. Rescue of *k135* root phenotype by various KIN7.3 fusions: *KIN7.3pro:GFP-KIN7.3*, *RPS5apro:KIN7.3-tagRFP*, or *RPS5apro:KIN7.3-mNeon*; quantification of root length (right). Data are means ± SD of five independent experiments each containing five measurements (2-tailed *t*- test.

**S2 Fig.**
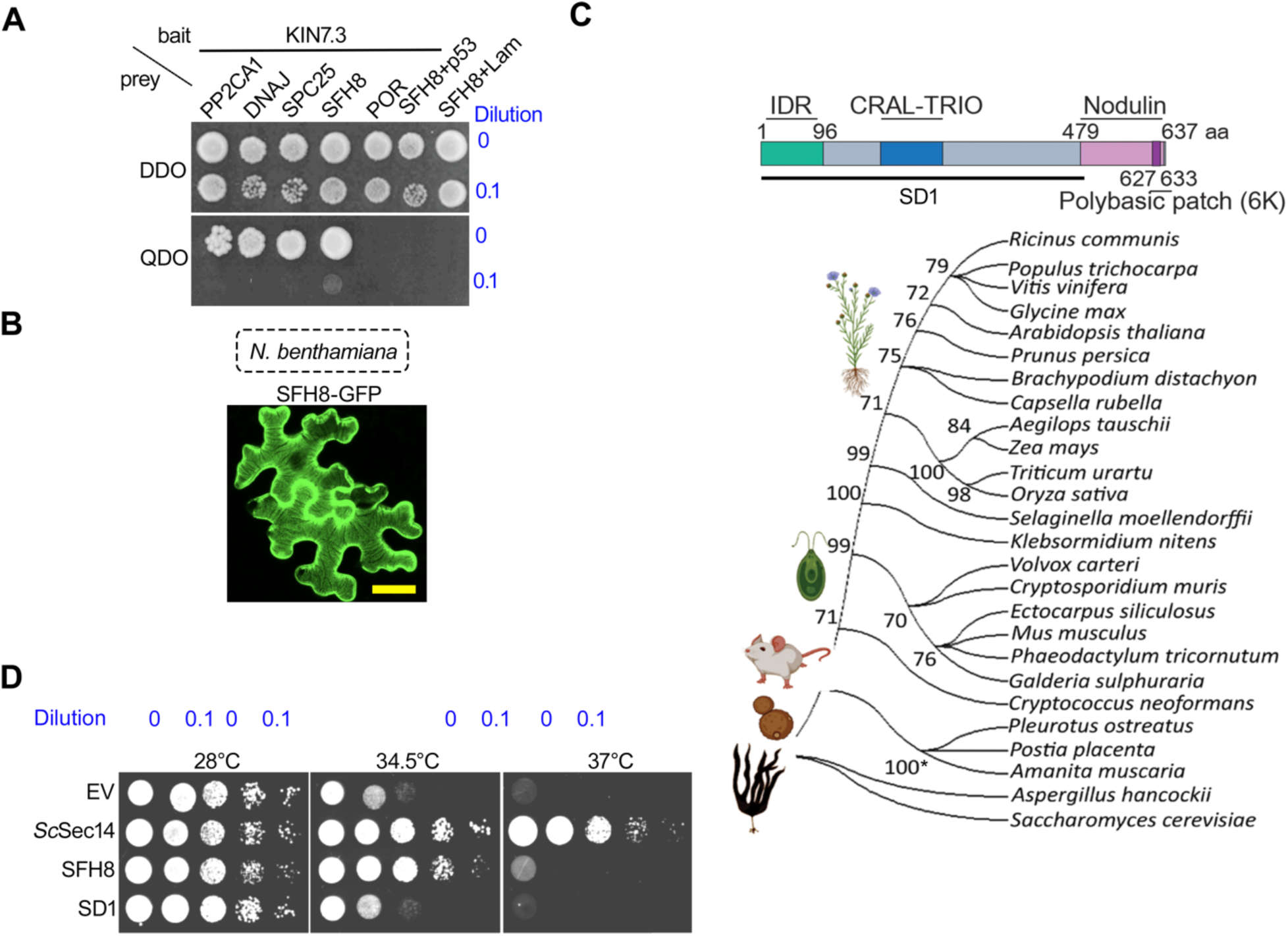
KISC associates with the lipid-transfer protein SFH8 and complements *sec14-1^ts^* yeast mutant. **A.** Identification of AT2G21520 (SFH8) as an interactor of KIN7.3. Y2H between binding domain (BD)-KIN7.3 and four putative interactors (PP2CA1, DNAJ, SPC25, and PORcino; see also **Supplemental Information** for a description of the KIN7.3 interactors which were not followed herein). AD, activation domain; DDO, double dropout (control); QDO, quadruple dropout (selection). Negative controls: p53 and lamin (Lam). Y2H was replicated four times. **B.** Representative confocal micrograph (maximum intensity projection) showing SFH8-GFP localization in *N. benthamiana* leaves (3 dpi, 35S promoter). Scale bar, 20 μm. The micrograph is from a single representative experiment replicated multiple times. **C.** Proposed model of SFH8 architecture (upper). Numbers indicate aa truncations used throughout the paper; SD1 corresponds to amino acid residues 1-478. IDR, intrinsically disordered region (1-96 aa). The CRAL-TRIO domain binds small lipophilic molecules and is named after the cellular retinaldehyde-binding protein and TRIO guanine exchange factor; phylogenetic analysis of SFH8 (AT2G21520) full-length sequence (lower). Numbers indicate bootstrap values with a confidence cutoff of 70. **D.** Yeast temperature-sensitive *sec14-1^ts^* loss-of-function mutant complementation by full-length SFH8 or SD1 (for truncations, see **B**). Note that at 37°C the complementation was moderate, likely due to the instability of the protein at elevated temperatures (37°C is much beyond the physiological Arabidopsis growth temperature which is between 22-28°C). Dilution series:0-10^-4^. The experiment was replicated three times.

**S3 Fig.**
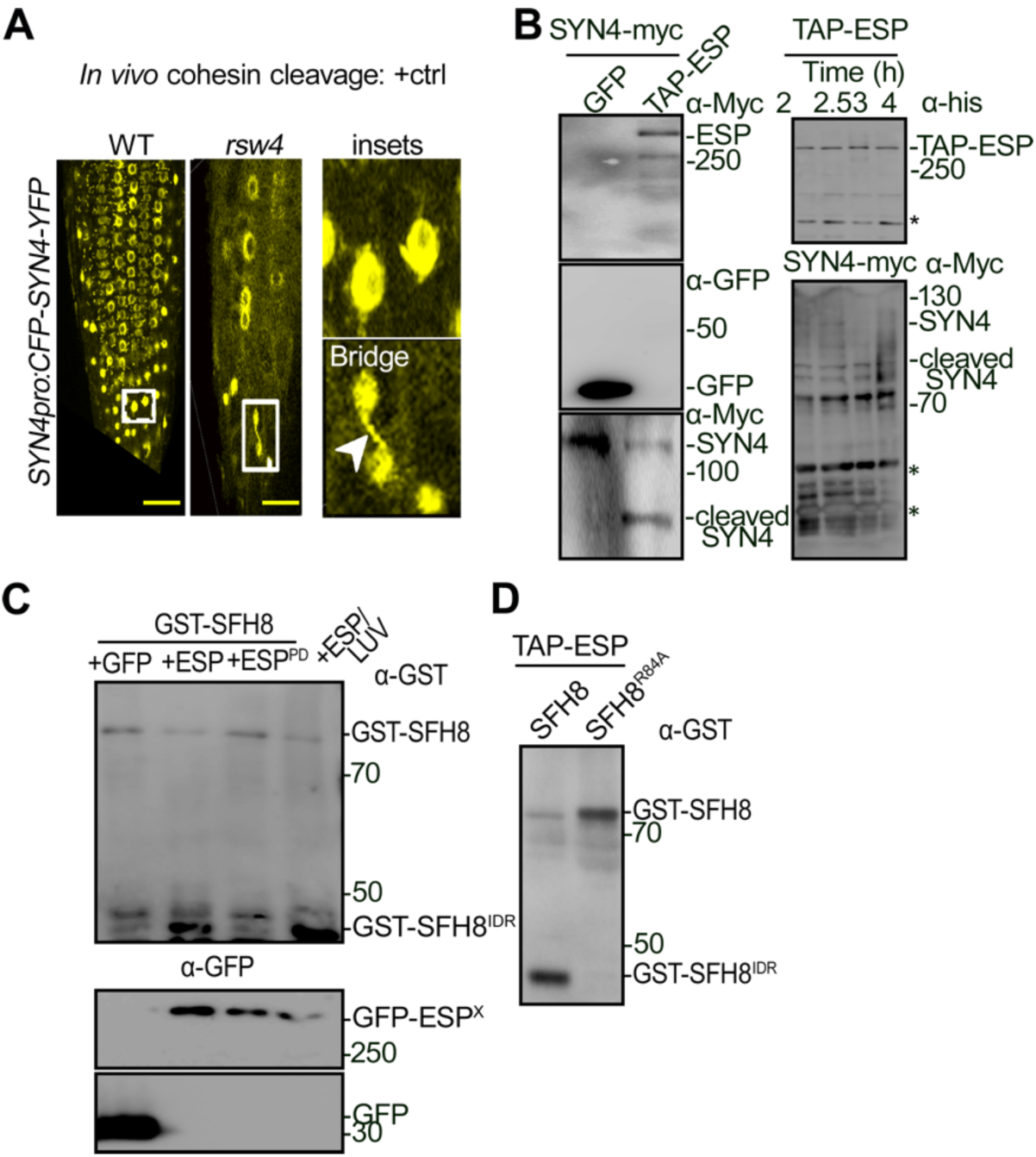
ESP cleaves the cohesin SYN4 and SFH8 in vitro. **A.** Establishment of a positive cleavage control for ESP: the mitotic cohesin SYN4 is an *in vivo* ESP target. Representative images of root cells expressing *SYN4pro:CFP-SYN4-YFP* in WT or *rsw4* (at the restrictive temperature of 28°C for 72 h, 5 -7 DAG). Insets show chromosomal bridges in *rsw4* due to the presumptive lack of the mitotic SYN4 kleisin subunit cleavage by ESP (epidermal cells); the white arrowhead denotes a chromosomal bridge. The experiment was replicated multiple times (with 1-3 bridges evident per root). Scale bars, 40 μm. **B.** SYN4 is cleaved by ESP *in vitro* confirming that purified ESP can be active. *In vitro* cleavage of SYN4 on beads (*35Spro:SYN4-myc*) purified from *N. benthamiana*, in the presence of immunopurified from *N. benthamiana* aTAP-ESP immobilized on beads (aTAP carries c-myc and hexahistidine tags), mitotically pre-activated by CyclinD (see **METHODS** for details on purification). Incubation of the cleavage assay was for 1 h at 37°C. The experiment was replicated three times. Right: increasing incubation time (2-4 h) led to the formation of additional cleavage fragments and smearing of SYN4-myc, suggesting that ESP may cleave SYN4 at multiple sites with differential preferences. Asterisks indicate non-specific bands. The experiments were replicated twice. **C.** Proteolytically inactive ESP fails to cleave SFH8, and cleavage is sustained in the presence of liposomes. *In vitro* GST-SFH8 cleavage by immunopurified from *N. benthamiana* GFP-tagged variants of ESP, ESP^PD^ (protease dead [81], and ESP in the presence of large unilamellar vesicles (LUVs; made from PC and containing 10 mol %PC and PS). The cleaved GST-SFH8^IDR^ is shown (band below 50 kDa). The beads carrying GFP or GFP-tagged ESP variants in the ±LUVs were co-incubated with GST-SFH8 at 37°C for 1 h. As the SFH8^ΔIDR^ was converted to filaments (see below) and due to the observed compromised full-length SFH8 transfer from the SDS-PAGE to the membrane, the immunoreactive signal increment of the SFH8^ΔIDR^/SFH8 (α-GST) ratio upon cleavage was not proportional to the SFH8^IDR^ production. The experiment was replicated four times. Note that the band around 50 kDa was detected in all samples and thus could correspond to a non-specific cleavage product in the GFP-trap setting. **D.** GST-SFH8^R84A^ is not cleaved by aTAP-ESP (similar experimental setting as in **B**). An additional product is due to non-specific cleavage during purification (above the specific cleavage product around 50 kDa). The experiment was replicated three times.

**S4 Fig.**
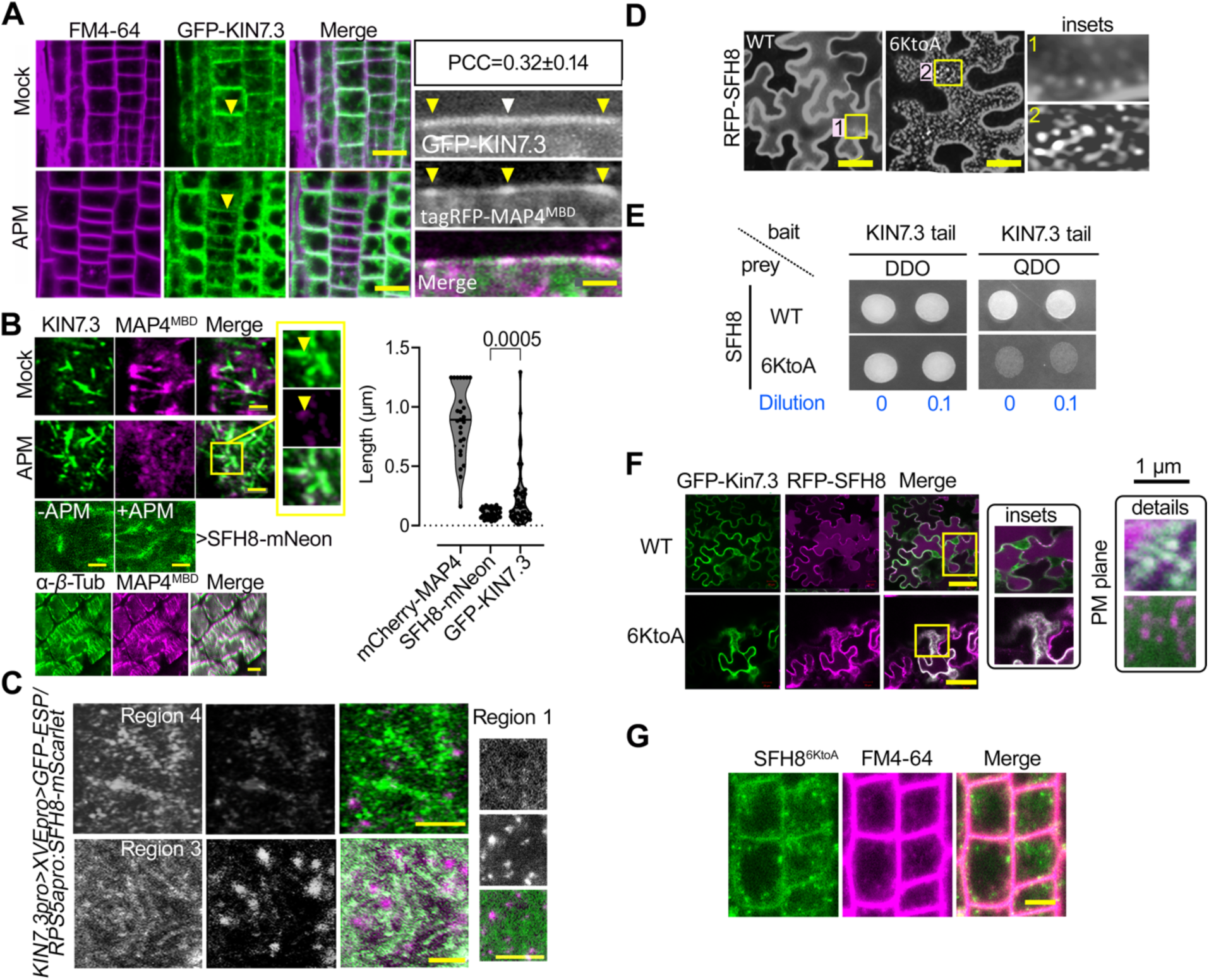
SFH8 liquid-like clusters do not colocalize with MTs or KISC and the positively charged patch of SFH8 is indispensable for its localization and polarity. **A.** KIN7.3 is only partially colocalized with MTs at the PM plane and is excluded from PM-resident MT bundles. Representative confocal images of *KIN7.3pro:GFP-KIN7.3/35Spro:MAP4^MBD^*expressing lines (5 DAG, region 3 epidermal cells) counter-stained with FM4-64, ±APM (10 nM, 1 h; left). Scale bars, 12 μm. Right: representative confocal images of GFP-KIN7.3 at cell contours (5 DAG, region 3 epidermal cells), and cell contour localization of *KIN7.3pro:GFP- KIN7.3/35Spro:MAP4^MBD^*. The box at the top right shows Pearson’s Correlation Coefficient (PCC) calculated at the regions indicated with the arrowheads, which in the case of MAP4^MBD^ denote PM-attached MT bundles. Scale bar, 1 μm. **B.** Short KISC filaments do not show significant signal collinearity with MTs. Representative confocal micrograph showing the partial colocalization of GFP-KIN7.3 filaments with MAP4^MBD^ in root region 3 (defined in **Figure 1**) ±APM (10 nM, 1 h). Note that KIN7.3 filaments are associated with bundled-MT remnants (reminiscent of clusters; insets, APM treatment). Lower: SFH8 filaments and their resistance upon APM treatment (10 nM, 1 h; region 3); colocalization between MAP4^MBD^/α-tubulin signals. Note that overexpression of MAP4 protein induces significant MT bundling. Scale bars, 0.1 μm (for filaments) or 3.4 μm for colocalization (MAP4/tubulin). Right: quantification of KIN7.3/SFH8 filaments’ length at the PM (six pooled independent experiments; n=40, Wilcoxon). Note the high density of SFH8 length counts around the mean, suggesting that SFH8 filaments are mostly short compared to MT filaments (note the MT length distribution) with little variability. KIN7.3 showed more dispersion, as shown by the violin plot (and the micrograph on the left), indicative of KIN7.3 association with MT filaments/bundles, as well. Scale bars, 1 μm. **C.** ESP is also restricted from SFH8 clusters in Arabidopsis roots. Representative confocal micrographs of the line co-expressing *KIN7.3pro*>*XVEpro>GFP-ESP*/*RPS5apro:mScarlet-SFH8*, after estradiol induction (20 μM, 16-24 h; 5 DAG, region 2 epidermal cells, PM surface at the apicobasal junction). Images were obtained at maximum scanning speed (40 frames per second) using the LIGHTNING super-resolution module of the SP8 Leica confocal microscope (63x W with an x/y-axial resolution of 120 nm upon deconvolution). Post-acquisition, images were deconvoluted using the LIGHTNING algorithm and water correction was set in the algorithm as the mounting medium. Similar settings were used in other images as well. Scale bar, 10 μm. **D.** The nodulin motif modulates SFH8 clustering propensity most likely through the repulsion from its lysine (K) residues. Upper: representative confocal micrographs showing the propensity to form clusters of N-terminally RFP-tagged SFH8 variants (SFH8^6KtoA^ under the 35S promoter) in *N. benthamiana* transient expression system. Insets (right) show details of clusters. The experiment was replicated two times. Scale bars, 20 μm. **E.** KIN7.3 tail fails to interact efficiently with SFH8^6KtoA^ (see also **Figure S1**). Y2H between the baits SFH8, or SFH8^6KtoA^ with KIN7.3 tail. KIN7.3 tail was fused to the binding domain (BD) and the SFH8 variants to the activation domain (AD). DDO, double dropout (growth control); QDO, quadruple dropout (selection medium for interaction). Two dilution series are shown (0 and 0.1). The experiment was replicated three times (N=3). **F.** Representative confocal micrographs showing colocalization analyses of RFP-SFH8, and - SFH8^6KtoA^ with GFP-KIN7.3 (*35Spro:GFP-KIN7.3*) in *N. benthamiana* transient expression system (3 days post-infiltration). Note that in cells expressing SFH8^6KtoA^, GFP-KIN7.3 retained a cytoplasmic localization (insets and details on the right), and the lack of GFP-KIN7.3 colocalization with RFP-SFH8^6KtoA^ clusters. Scale bars, 50 μm. The experiment was replicated twice (N=2). **G.** Representative confocal micrographs showing the reduced levels of SFH8^6KtoA^ and lack of polarity of mNeon-SFH8^6KtoA^ (green; *sft8 RPS5apro:mNeon-SFH8^6KtoA^*; epidermal cells 5 DAG, region 3) counterstained with the dye FM4-64 (magenta; PM and endosomes staining) from line #2 shown in **A**. Scale bar, 25 μm.

**S5 Fig.**
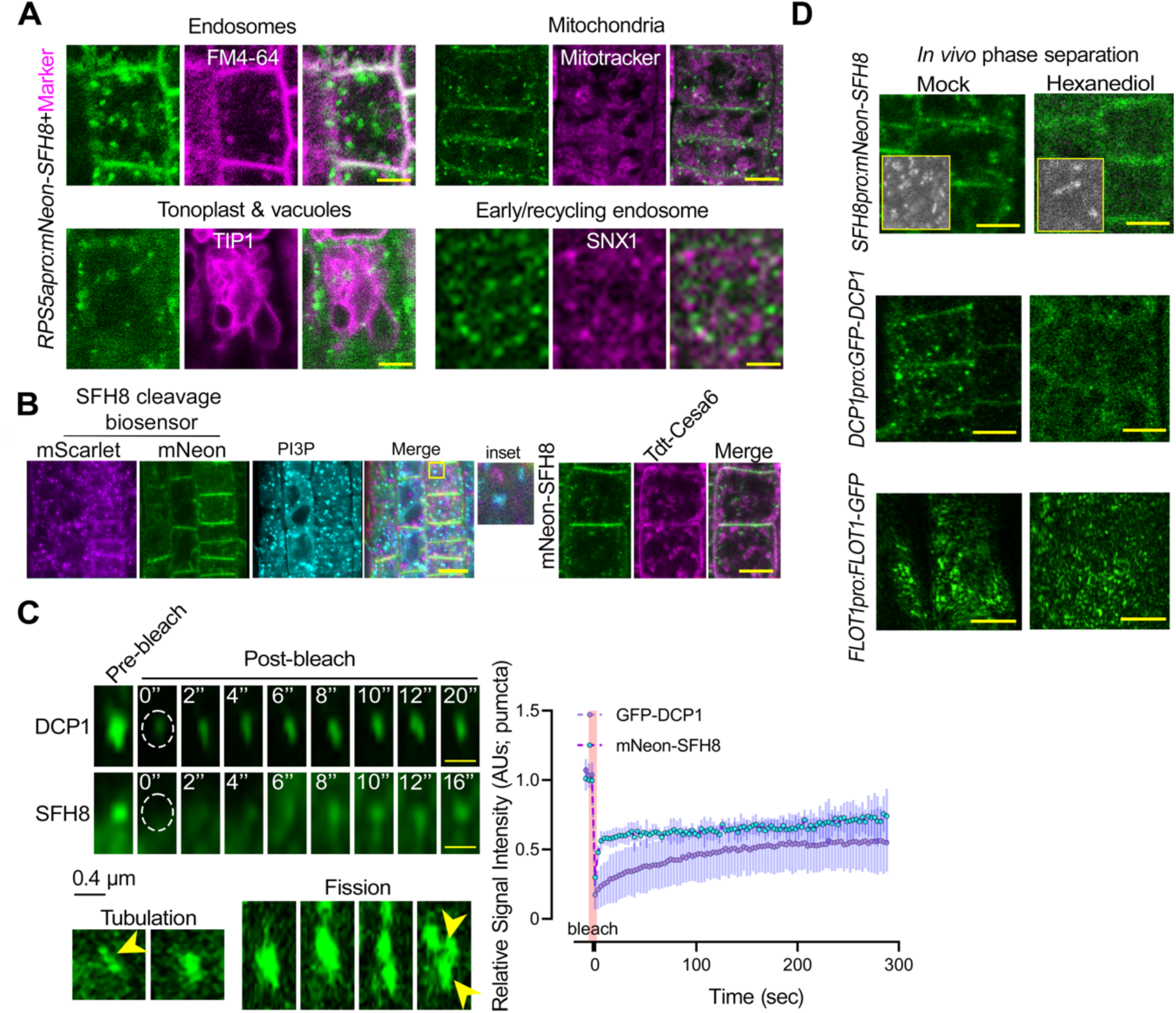
SFH8^IDR^ upon its removal from SFH8 is stable and does not colocalize with membrane markers or the cellulose synthase complex. **A.** Representative confocal micrographs from *RPS5apro:mNeon-SFH8* expressing lines counterstained with FM4-64 (endosomes and plasma membrane) or mitotracker (mitochondria) dyes; lines co-expressing *RPS5apro:mNeon-SFH8* with *35Spro:TIP-RFP* (tonoplast intrinsic protein), or *35Spro:SNX1-RFP* (sorting nexin 1; *trans*-Golgi network). In some cases, the duration of tracking was limited to the amount of time that particles remained in a single focal plane, as the required acquisition rate did not permit the collection of z-stacks (voxels); the median track length was 35 frames, corresponding to 20’’ of imaging. PCC analyses failed to show colocalization of SFH8 cytoplasmic puncta with any of the markers/dyes used (negative values). Scale bars, 2 μm. **B.** Although SFH8 binds to PtdIns(3)P, it does not colocalize with relevant PtdIns(3)P-positive structures *in vivo*. Representative confocal micrograph of lines co-expressing mScarlet-SFH8- mNeon (green; *RPS5apro:6xhis-3xFLAG(HF)-mScarlet-SFH8-mNeon*) with PtdIns(3)P (tagged with CFP and expressed under pUBI10). The inset on the right is digital magnification, showing a lack of colocalization between PtdIns(3)P and mScarlet-SFH8^IDR^. Right: mNeon-SFH8 does not associate with cellulose synthase complex-puncta. Representative confocal micrograph of lines coexpressing *RPS5apro:mNeon-SFH8* with Tdt-CESA6 (CELLULOSE SYNTHASE 6 isomer under the 35S). Note the lack of colocalization between the two proteins. Scale bar, 4 μm. The experiments were replicated three times for various developmental stages (2-7 DAG), and root tissues (regions 1-4). **C.** SFH8 cytoplasmic puncta show features of LLPS proteins. Upper: representative confocal micrographs showing the FRAP signal recovery as a fraction of time in lines expressing *35Spro:GFP-DCP1* (a protein known to form condensates in both plants and animals; [82]) or *SFH8pro:mNeon-SFH8*. The regions of interest were set on mobile cytoplasmic puncta (at the midsection of the cell; 7 DAG, epidermal cells of region 3-4). Note that the duration of tracking was limited to the amount of time that the observed particles remained in a single focal plane, as the required acquisition rate did not permit the collection of z-stacks (voxels). Middle: confocal time-lapse imaging (2 sec time interval) showing that mobile puncta can freely flow and deform around surfaces of other structures, as well as undergo tubulation, fusion, or fission. Arrowheads denote the coalescing puncta. Lower: FRAP signal recovery (normalized integrated density) as a fraction of time. The red faded band parallel to the Y-axis indicates laser iterations time. Data are means ± SD (N=3, n=1). Scale bars, 0.4 μm. **D.** SFH8 puncta and clusters show features of LLPS proteins. Micrographs from 10% (v/v) 1 h 1,6-hexanediol treatment of pDCP1:DCP1-GFP, *SFH8pro:mNeon-SFH8* or *FLOT1pro:FLOT1:GFP* expressing lines (5 DAG cell surface and midsections, epidermal root cells regions 2-3). Note that in the case of DCP1 and SFH8 cytoplasmic puncta were converted to a diffused cytoplasmic signal. DCP1 decorates the LLPS compartment known as processing bodies and was used as a positive control for 1,6-hexanediol efficiency under our conditions. FLOT1 decorates microdomains at the PM [83]. The insets are from the PM surface (TIRFM), showing that 1,6-hexanediol dissolved the SFH8 clusters. Scale bars, 5 μm.

**S6 Fig.**
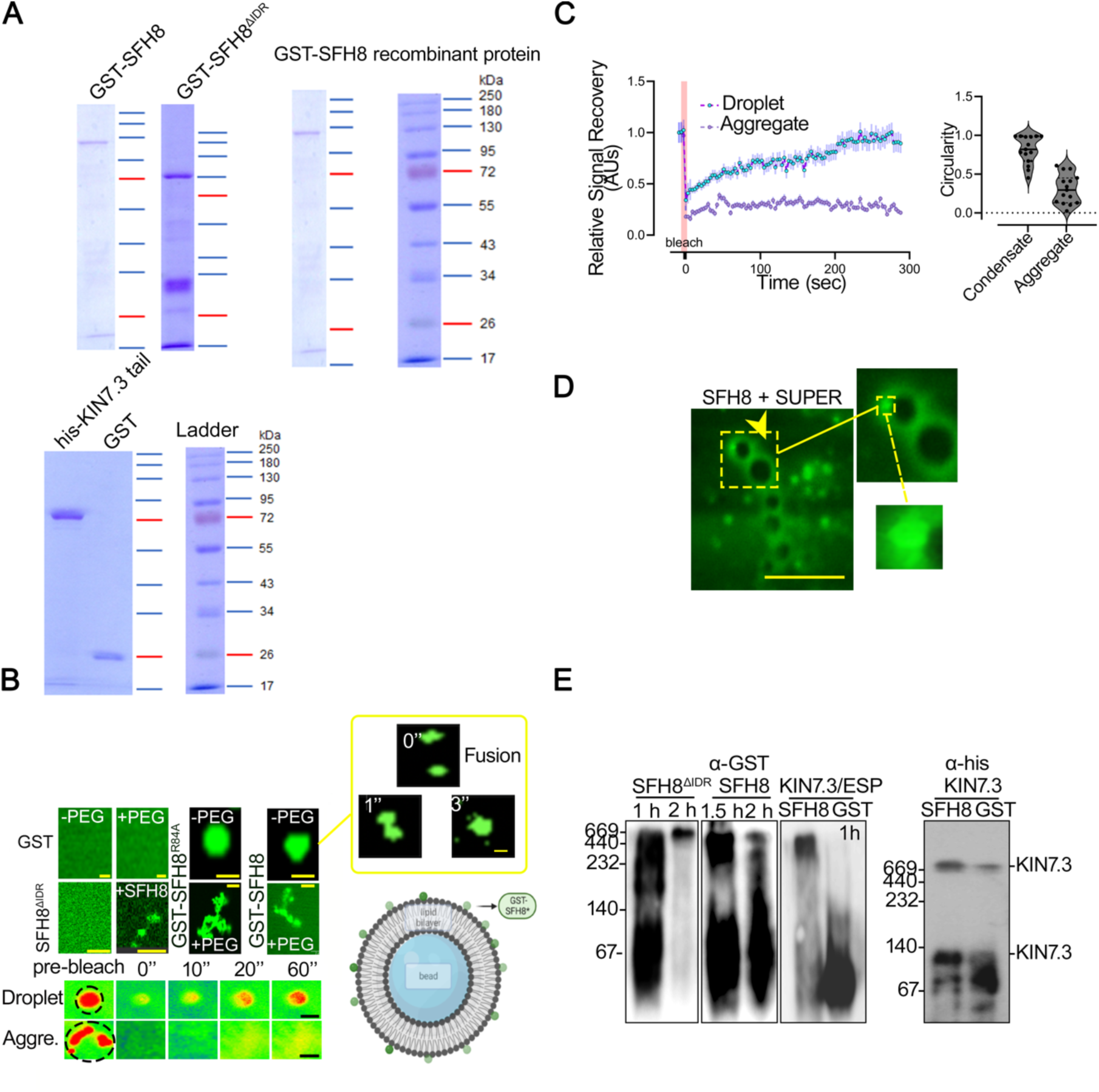
SFH8^IDR^ is functionally important and upon its removal from SFH8 is stable and does not colocalize with membrane markers or the cellulose synthase complex. **A.** Examples of purified proteins for assays. **B.** In vitro LLPS assays for SFH8 variants. Representative confocal micrographs of *in vitro* recombinant GST, GST-SFH8, GST-SFH8^ΔIDR^, and GST-SFH8^R84A^ proteins stained with Alexa638 in the presence or absence of the PEG3000 crowding agent in LLPS conditions visualized in fluidic chambers (see **METHODS**). Note that in the presence of PEG, SFH8 and SFH8^R84A^ switch to agglomerate-like states, while GST-SFH8^ΔIDR^ induced filamentous transition to GST-SFH8. Scale bars, 0.2 μm. Images are from a single representative experiment replicated three times. **C.** representative confocal high-speed micrographs showing droplet and aggregate signal recovery, and quantification of FRAP signal recovery (normalized integrated density) as a fraction of time of GST-SFH8^R84A^ droplets and aggregates. Scale bars, 0.2 μm. The red faded band parallel to the Y-axis indicates laser iteration time. Data are mean ± SD (N=5; n=1-3). Right: GST- SFH8^R84A^ droplet fusion in the same experiment. **D.** SFH8 can phase separate *in vitro* on model membranes more effectively than in the bulk phase. Representative high-speed super-resolution (120 nm) images from a SUPER template experiment show that GST-SFH8 labeled with Alexa638 (0.1 μM of GST-SFH8 in the assay) can form liquid-like droplets on membranes. Furthermore, GST-SFH8 showed an increased propensity to undergo LLPS in the presence of SUPER templates (see arrowhead denoting droplet; some droplets were released in the bulk phase). Data are representative (N=3). **E.** SFH8 conversion from LLPS to high molecular weight assemblies. Native gel electrophoresis of SFH8^ΔIDR^ and full-length SFH8. FL, SFH8. Western blots with α-SFH8 showed the rapid conversion of recombinant GST-SFH8^ΔIDR^ to agglomerates (60 min); very few agglomerations were observed for GST-SFH8. Note that in the presence of Kin7.3/ESP GST-SFH8 converted faster to high-molecular-weight assemblies. Right: detection of Kin7.3 using α-his. Western blots are from a single representative experiment replicated twice.

**S7 Fig.**
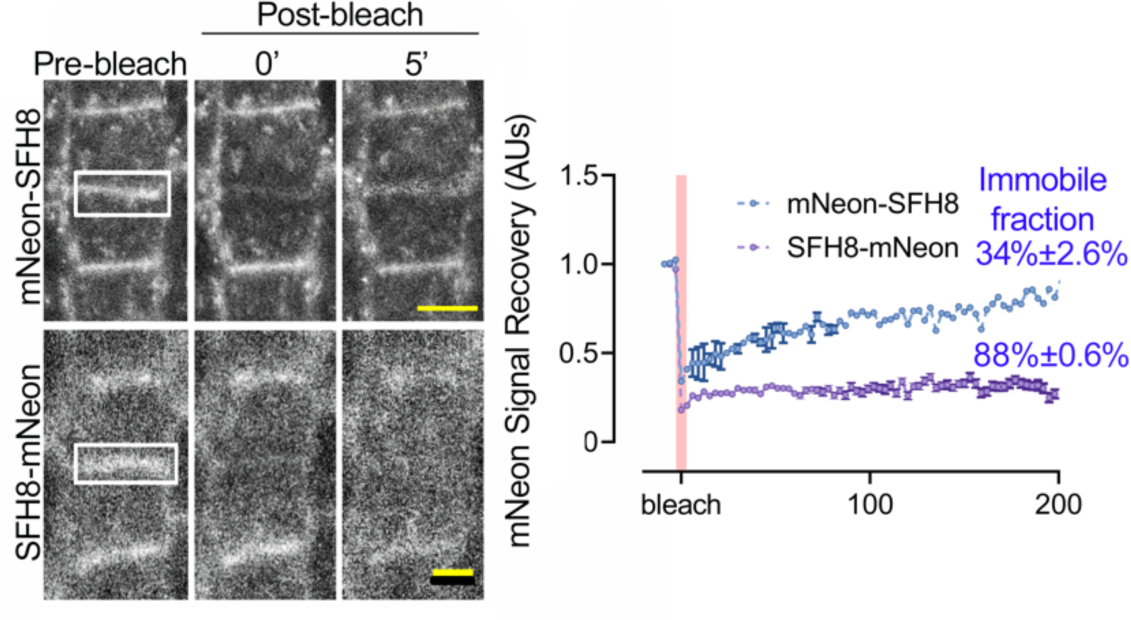
The C-terminus of SFH8 is less diffusive. The C-terminally tagged SFH8 shows a different diffusion rate than the N-terminally tagged. Representative confocal micrograph showing FRAP signal recovery (normalized integrated density) as a fraction of time of the N or C-terminally tagged SFH8 with mNeon on the PM (under the *SFH8* promoter), implying increased diffusion (due to cleavage, liquidity, and dwelling time of clusters). The white rectangular denotes the region of interest that was bleached. Scale bars, 5 μm. The N-terminally tagged SFH8 diffuses from the PM. Quantification of FRAP signal recovery as a fraction of time from cells expressing *SFH8pro:mNeon-SFH8* or *SFH8pro:SFH8-mNeon* lines (in the *sfh8* background) on the PM (5-7 DAG, epidermal cells of region 3; upper graph), Percentages indicate immobile fractions for N- and C-terminally tagged SFH8 with mNeon or N-tagged SFH8. The red faded band parallel to the Y-axis indicates laser iteration time. Data are means ± SD (N=3, n=3). Asterisks: significance at p<0.0001; Wilcoxon.

**S8 Fig.**
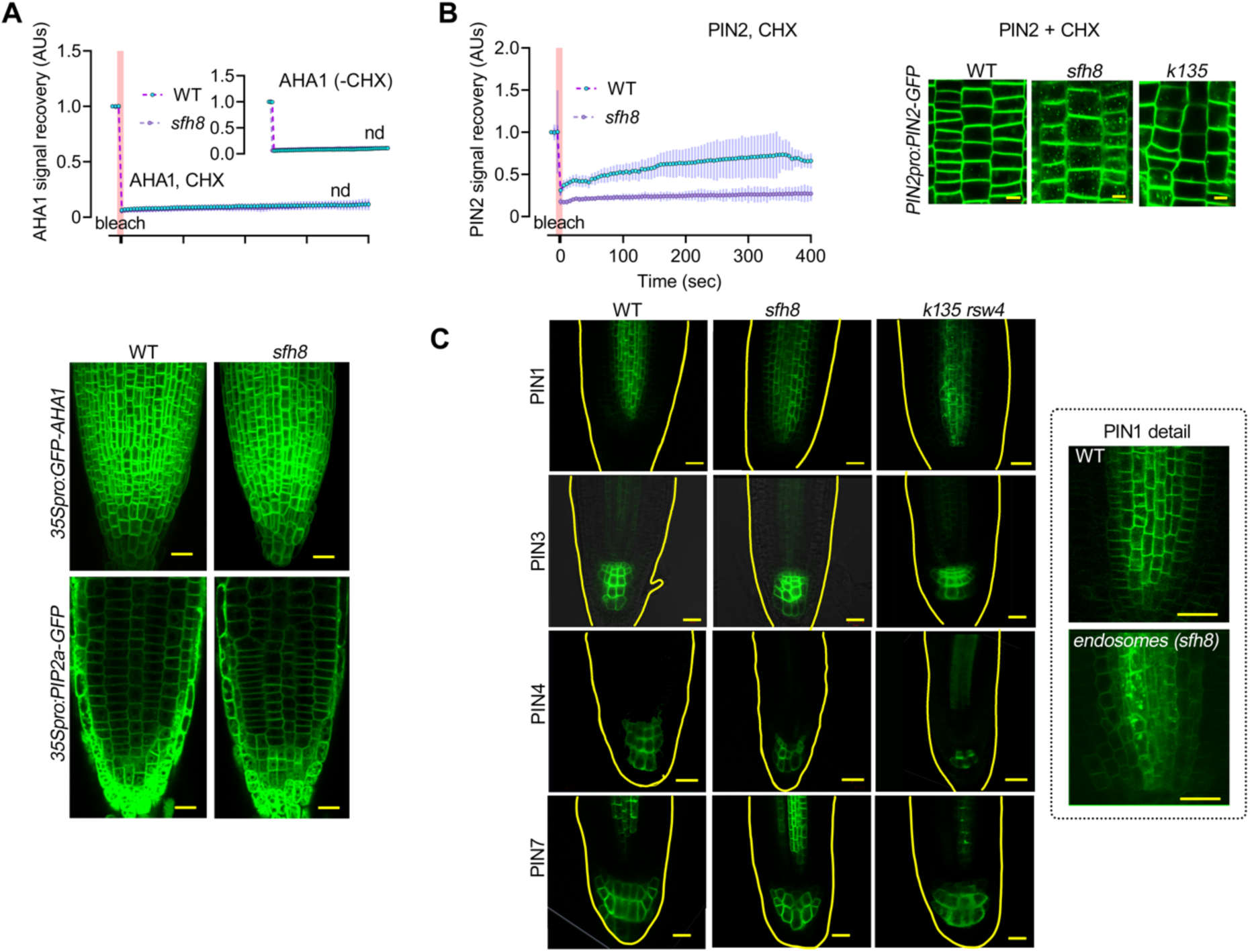
SFH8 may exert effects on fusion of polar proteins. **A.** SFH8 does not seem to be a general regulator of the secretory pathway; the non-polar AHA1 recovery rate at the PM is not affected by *de novo* protein synthesis. Quantification of FRAP signal recovery (normalized integrated density) as a fraction of time from *35Spro:GFP-AHA1* (upper; inset graph without CHX). Lower: representative confocal micrographs of AHA1-GFP in WT and *sfh8* on the PM of epidermal root rip cells (5 DAG, region 3) expressing *35Spro:GFP-AHA1*, or *35Spro:PIP2a-GFP*. Scale bars, 20 μm. Images are representative of an experiment replicated N˃10. **B.** Quantification of FRAP signal recovery (normalized integrated density) as a fraction of PIN2- GFP in WT and *sfh8*. Right: GFP-PIN2 retention in *sfh8* and *K135* endosomes is shown (30 min CHX treatment, right). Note that CHX did not lead to the dissolution of PIN2 endosomes. In **A** and **B** the red faded band parallel to the Y-axis indicates laser iterations time (five or more independent experiments with one replicate each; n=5-8). Ιmages are representative of an experiment replicated two times (N=2). **C.** Representative confocal micrographs showing PIN1, PIN3, PIN4, and PIN7 delivery at polar domains in WT, *sfh8*, and *rsw4* (left) expressing *PIN1pro:PIN1-GFP*, *PIN3pro:PIN3-GFP*, *PIN4pro:PIN4-GFP*, or *PIN7pro:PIN7-GFP*, 5 DAG. PIN1 levels were also reduced in the mutants, while endosomes could also be observed. Note that slight level perturbations were also observed for PIN3, 4, and 7 in the *sfh8* and *k135 rsw4* (24 h at the restrictive temperature 28°C; 5 DAG), but due to different patterning of the columella cells in these mutants these are hard to follow-up. On the other hand, no endosomes were visible for these proteins in the mutant backgrounds, but their signals were reduced. Bottom: localization details for PIN1; note that in some cases PIN1- GFP showed endosomes accumulating in the *sfh8* (mostly) and *k135 rsw4*, suggesting that PIN1 delivery was also compromised albeit to a lesser extent (as also discussed in [73] for the *rsw4* mutant). Scale bars, 20 μm. Images are representative of an experiment replicated N˃10.

**S9 Fig.**
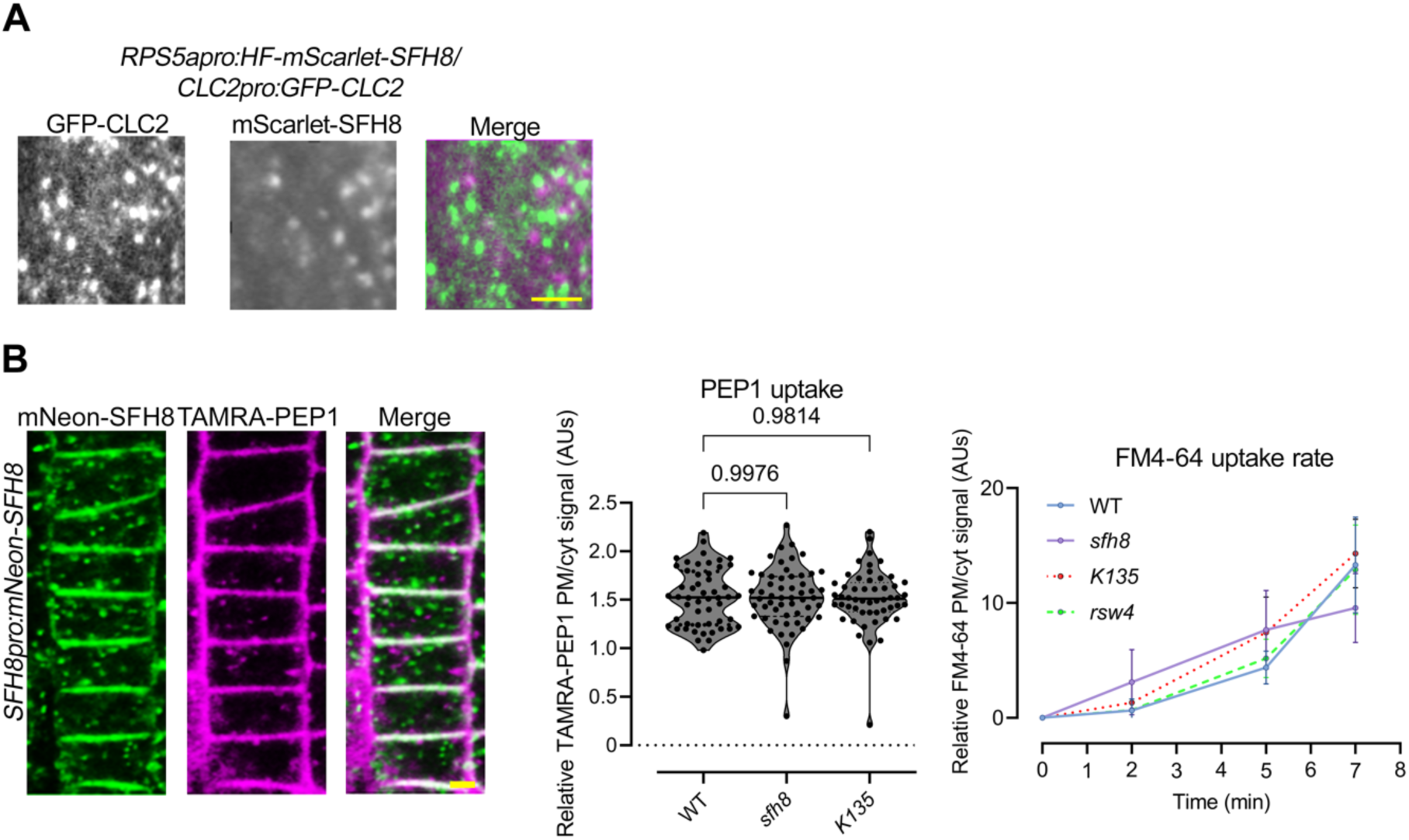
SFH8 and KISC do not significantly affect bulk endocytosis. **A.** Pepresentative dual-channel TIRFM micrographs showing the lack of colocalization between mScarlet-SFH8 (magenta) and CLC2-GFP (green; mock) from *SFH8pro:SFH8- mNeon*/*CLC2pro:CLC2-GFP* expressing lines. Scale bar, 0.2 μm. Images are representative of an experiment replicated multiple times, irrespective of the root region examined. **B.** Lack of colocalization between SFH8-mNeon and TAMRA-PEP1 (left; stains TGN and PM clusters), and quantification of the rate graphs showing TAMRA-PEP1 and FM4-64 uptake in WT, *sfh8*, and *k135* (violin plot and rate plot, respectively; three independent experiments; n=9, Wilcoxon-other tests produced comparable results). Scale bars, 2 μm.

**S10 Fig.**
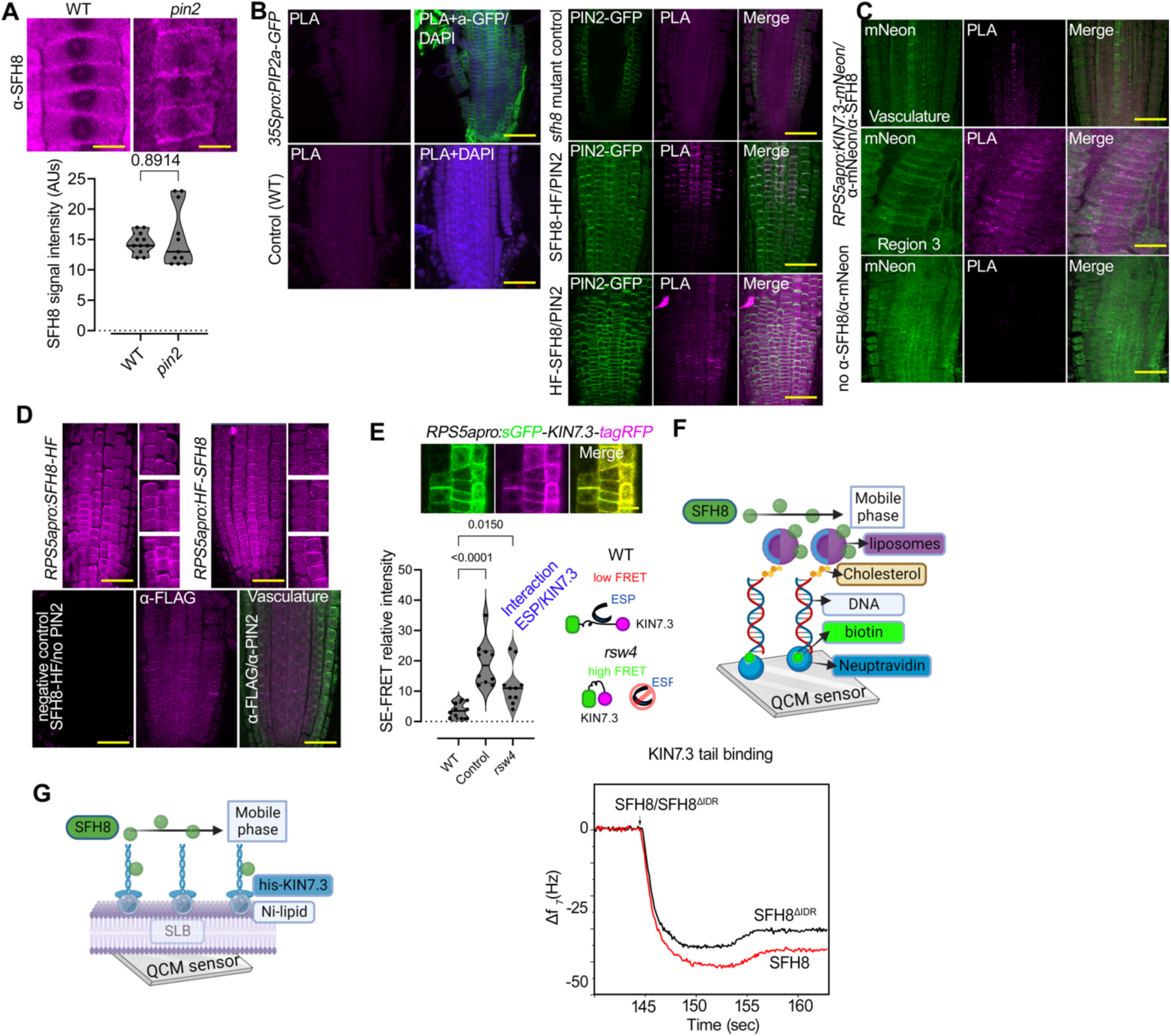
Controls for SFH8/PIN2 interactions and liposome fusion assays. **A.** PIN2 is required for the recruitment of SFH8 at the PM. Representative confocal micrographs showing α-SFH8 signal in WT, and *pin2* mutant background (5 DAG, region 3 cortex cells), and quantification of α-SFH8 signal intensity in WT, and *pin2* mutant (lower; three independent experiments with signal calculations from cortex cells from regions 3-4 with three roots each; n=9, Wilcoxon). Scale bars, 5 μm. **B.** Technical controls testing the specificity of the PLA approach in our settings (PIN2/SFH8 PLA assay), showing also that detected interactions are selective and SFH8 is not promiscuously interacting with other proteins. SFH8 did not interact with PIP2a (α-GFP/ α-SFH8), while SFH8- HF (α-FLAG) interacted extensively with PIN2 (α-GFP; region 3 onwards at the PM, epidermis and cortex cells). Note the significantly reduced PLA-positive signal in HF-SFH8/PIN2 (bottom panel; PLA signal was observed at the cell plate as well). The *sfh8* mutant was used as a negative control for the PLA approach (specificity of α-SFH8 in a PLA setting; sometimes a nuclear, likely non-specific signal was observed in negative controls). Scale bars, 50 μm. Images are representative of experiments replicated five times. **C.** SFH8 and KIN7.3 interact significantly in regions 3-4 (epidermis and cortex cells). PLA of α-mNeon and SFH8 and inset (right) showing the increased interaction strength towards region 4 (for example in the vasculature; upper images). The “no -α-SFH8” corresponds to negative control (bottom) in which only α-mNeon was used and thus did not produce a positive PLA signal. Scale bars, 50 μm (for “region 3” panel, 10 μm). Images are representative of an experiment replicated five times. **D.** FLAG epitope staining confirms SFH8 localization. α-FLAG signal in HF-SFH8 or SFH8-HF expressing lines in the *sfh8* mutant (5 DAG). Insets denote the localization at the corresponding region. Images are representative of an experiment replicated twice. Scale bars, 50 μm. **E.** KIN7.3 interacts with ESP at the PM. SE-FRET experiments at the PM show that N and C termini of KIN7.3 fold over in the absence of ESP (*rsw4*; see also [20] for a similar situation upon MT binding), suggesting also that ESP/KIN7.3 interact at the PM. Scale bar (micrograph), 5 μm. “control”: *35Spro:sGFP-tagRFP* expressing lines (yielding high FRET). Data are means±SD (N=3, n=3 roots; Wilcoxon). **F.** Control experiments show that SFH8/SFH8^ΔIDR^ do not differ in their affinity for LUVs. As liposome co-flotation assays take significant incubation time and thus can be biased towards the increased molecular weight and sedimentation propensity of SFH8^ΔIDR^ due to its filamentous conversion and reduced hydrodynamic radius leading to erroneous affinity determination, we designed a semi-quantitative *in vitro* assay based on Quartz Crystal Microbalance with Dissipation monitoring (QCM-D). We used the QCM device as a sensitive mass sensor monitoring the frequency response (Δf) of the acoustic wave during binding events on the surface of a model membrane [5]. The device surface was first covered with neutravidin (layer 1) which was used to bind specifically a 5’-biotinylated DNA (layer 2). This DNA was also modified at the 3’-end with a cholesterol moiety, further employed to anchor a liposome (layer 3). We prepared 50 nm diameter liposomes with two different lipid compositions: DOPC:PI(4,5)P2 (99:1, n/n) and DOPS:PI(4,5)P2 (99:1, n/n). Finally, proteins (e.g., GST-SFH8) were infused (layer 4) on top of layer 3. **G.** Determination of KIN7.3 binding on GST-SFH8 and GST-SFH8^ΔIDR^. Again, due to the filamentous conversion of GST-SFH8^ΔIDR^ as described above (see **F**), we designed a real-time binding assay with a supported lipid bilayer approach for KIN7.3 tail affinity determination: GST- SFH8 and GST-SFH8^ΔIDR^ protein binding on QCM-D coupled liposomes. Lower: real-time curves showing dynamic binding through the frequency response of QCM-D obtained upon infusion of full-length SFH8 (upper panel) or GST-SFH8^ΔIDR^ (bottom panel) proteins on supported lipid bilayers carrying hexahistidine-tagged KIN7.3 tail immobilized on the supported lipid bilayer via DGS-NTA(Ni) lipid anchor. Only slight, insignificant differences were observed in the binding between the two protein variants. As of note, SFH8/SFH8^ΔIDR^ differ in molecular mass and thus SFH8 should give a slightly higher response in Δf as shown. Data are representative (N=3).

**S11 Fig.**
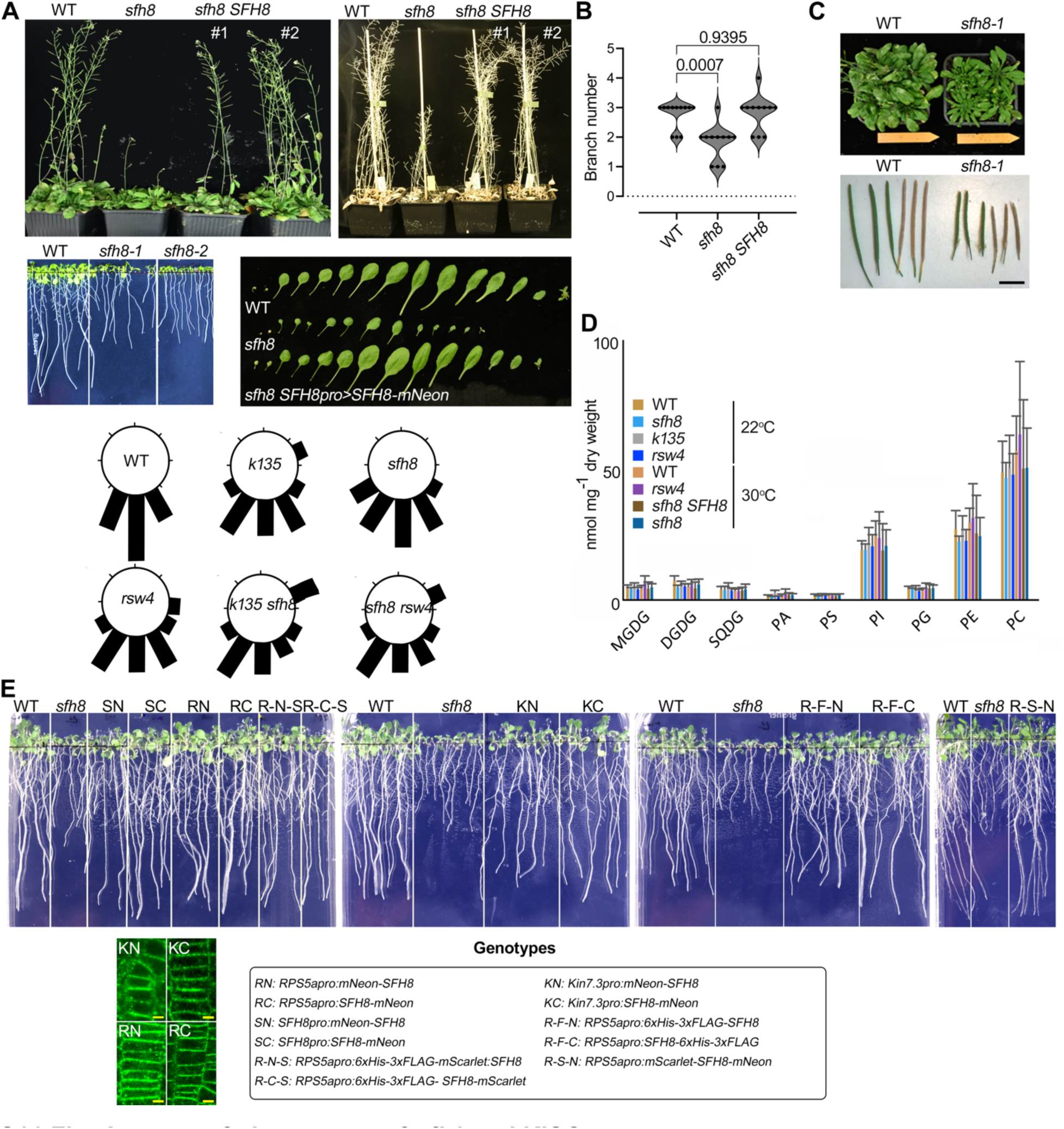
Aspects of phenotypes of *sfh8* and KISC mutants. **A.** Adult plant phenotype (upper, 21 DAG, 4 plants per pot) and representative photos of WT, *sfh8-1,* and *sfh8-2*, and *sfh8 proSFH8:SFH8-mNeon* lines at 10 DAG (lower left, *sfh8 SFH8*). Hereafter, *sfh8-1* is denoted as “*sfh8*”, and due to phenotypical resemblance with *sfh8-2* allele is used throughout the paper. Bottom: circular plots showing moderate gravity perception defects of *sfh8, k135, rsw4*, the double *rsw4 sfh8,* and *k135 sfh8* mutants grown vertically on plates at 7 DAG. Circular plots are from a single representative experiment replicated multiple times. **B.** SFH8 mutation affects some auxin-related phenotypes apart from positive root gravitropism. Quantification of branch number (three independent experiments; n=12, ordinary 1-way ANOVA). **C.** SFH8 mutation impact overall growth and fecundity. Representative images of adult WT and *sfh8* plants in short-day (10-h light /14-h dark) condition (upper), and silique lengths (lower). Length bar, 0.8 cm. **D.** SFH8 or KISC do not affect lipid profiles of the PM. LC-MS/MS lipid species quantitative analysis in WT, *sfh8*, *k135,* and *sfh8 SFH8* complementation line (*sfh8 SFH8pro:mNeon-SFH8*) and the *rsw4* mutant (5 DAG; treated for 24 h at 28°C as the *rsw4* is a temperature sensitive allele; [20]). Data are means±SD (N=5, n=1; significance was tested with various tests). MGDG, monogalactosyldiacylglycerol; DGDG, digalactosyldiacylglycerol; SQDG, sulfoquinovosyl diacylglycerol; PA, phosphatidic acid; PS, phosphatidyl-serine; PI, phosphatidyl-inositols; PG, phosphatidylglycerol; PE, phosphatidylethanolamine; PC, phosphatidylcholine. **E.** SFH8 mutation can be rescued by SFH8 driven by *RPS5a* or *KIN7.3* promoters. Representative images of seedlings 10 DAG expressing various fusions of SFH8 in the *sfh8* mutant background. The representative confocal micrographs show localization of KN/KC, RN/RC (5 DAG, region 3 epidermal cells). Scale bar, 3.5 μm.

**S12 Fig.**
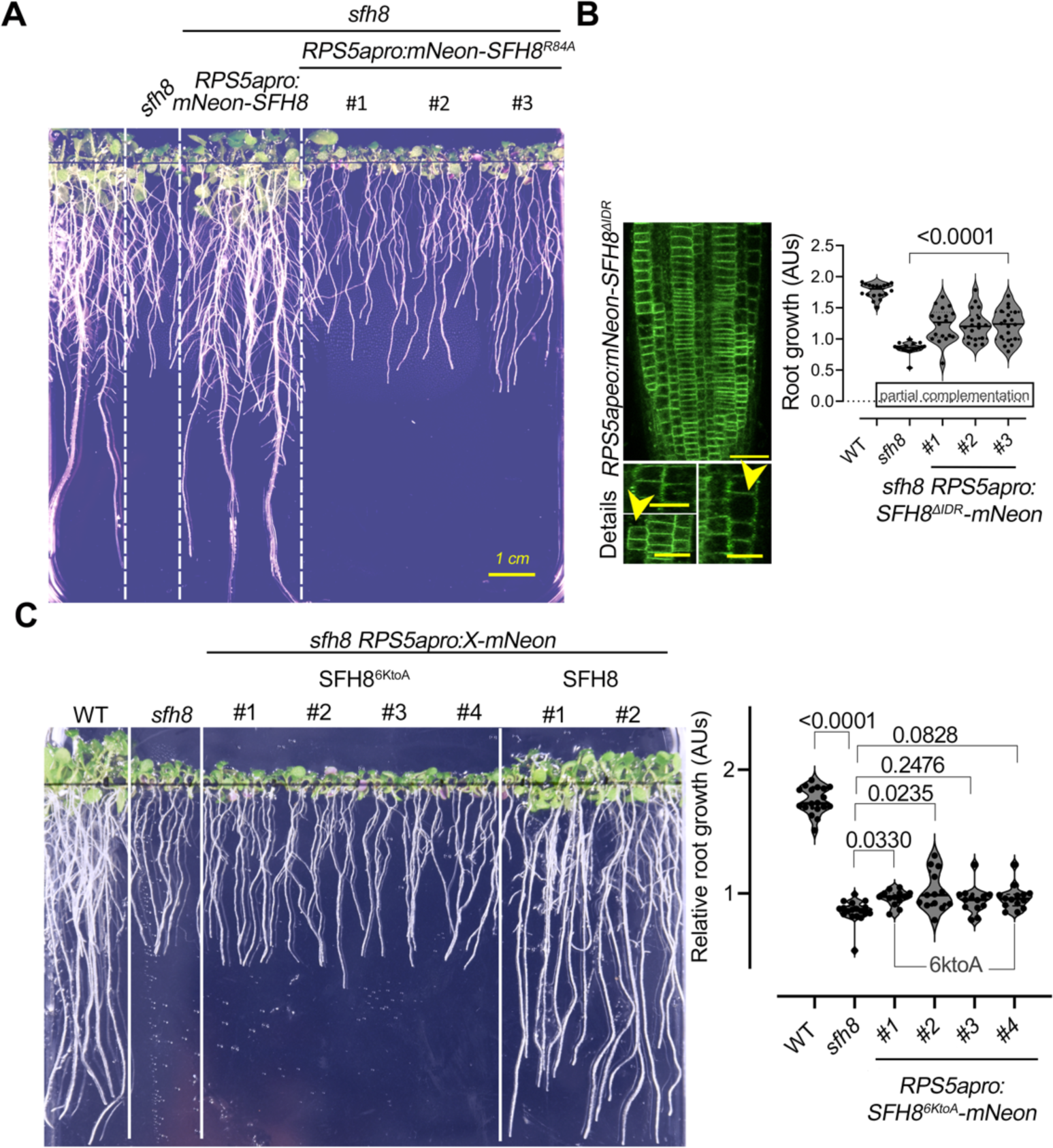
KISC/SFH8 can affect the robustness of auxin signaling. **A.** Reduction of *sfh8* rescue by the *mNeon-SFH8^R84A^* variant. Three-individual lines are shown. **B.** Representative confocal micrographs from meristematic epidermal root cells expressing *RPS5apro:mNeon-SFH8^ΔIDR^* in the *sfh8* mutant background show details of localization with a lack of filaments, polarization, and cytoplasmic puncta formation. Also note the reduced robustness of PM localization, uneven SFH8 levels, and perturbations of growth realized as reduced growth anisotropy (details; middle right cell denoted by arrowhead). The scale bar on the top, 40 μm and all others, 20 μm. Right: quantification of root length of three lines expressing *RPS5apro:mNeon-SFH8^ΔIDR^* in the *sfh8* background (two independent pooled experiments; n=18; ordinary 1-way ANOVA). **C.** Images showing partial complementation of *sfh8* by *RPS5apro:mNeon-SFH8^6KtoA^*in 4 individually transformed lines (10 DAG). Two complementation lines with SFH8 are also shown. Right: quantification of root growth from WT, *sfh8*, and four lines (presented in **A**) expressing *RPS5apro:mNeon-SFH8^6KtoA^* in the *sfh8* background (N=2; n=30, ordinary 1-way ANOVA).

**S13 Fig.**
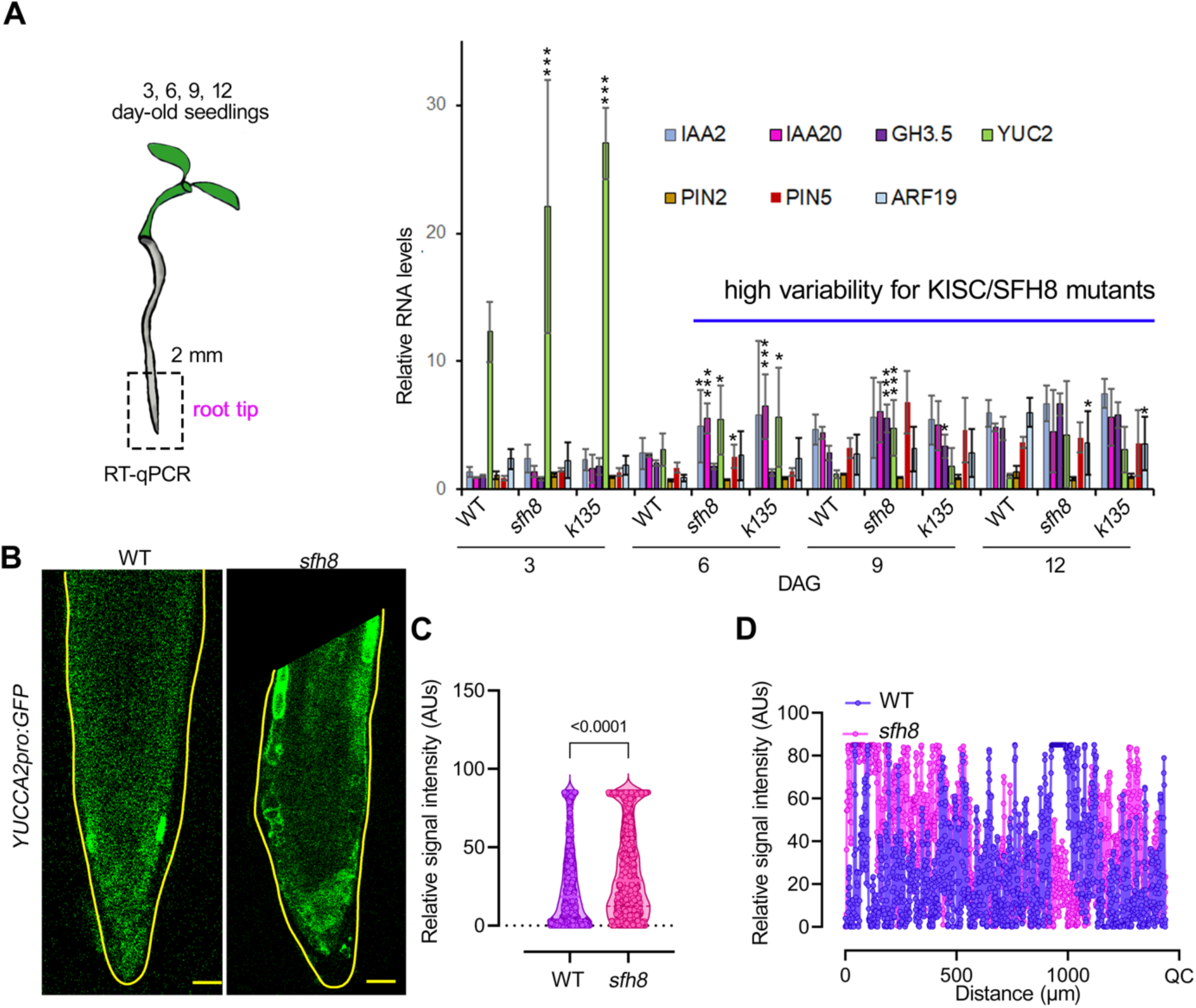
KISC/SFH8 can affect the robustness of auxin signaling. **A.** Reverse transcription quantitative PCR (RT-qPCR) setting for evaluation of changes in auxin-signaling genes (left). Two-millimeter root meristems were cut and processed for RNA extraction. Right: relative expression of the indicated genes normalized against *ACTIN7 and PP2A* (mean±SD from three independent experiments with two technical replicates; 2-tailed *t-*test to test robustness). Note the high variability of gene expression in the mutants (expanded SD). Asterisks: significance at p<0.05 (*) or p<0.0001 (***); *t*-test. **B.** Confirmation of reduced robustness of expression in *sfh8* of the auxin biosynthetic gene *YUCCA2*. Representative confocal micrographs showing *YUCCA2pro:GFP* expression (5 DAG root meristem cells). Images are representative of experiments replicated five times. Scale bars, 20 μm. **C.** Quantification of *YUCCA2pro:GFP* signal distribution (signal integrated density), in WT and *sfh8* (three independent experiments with three roots each; n=1,492 points across the root; 2- tailed *t*-test). **D.** Plot profile showing relative signal intensity along the root in WT and *sfh8*. QC, quiescent center (single representative experiment, replicated three times). Note the aberrant intensity peaks of *YUCCA2pro:GFP* in *sfh8*.

## Supplemental Information

### Localization of KIN7.3 interacting proteins in stable or transient expression

We expressed *PP2C*, *Spc25*, and *POR* in stable Arabidopsis Col-0 lines or *N. benthamiana*. The localization of these proteins is shown (**Fig 1**). Furthermore, we identified homozygous mutants of these proteins, showing a lethal phenotype for *spc25* and *por* mutants confirming previous data [84, 85].

The phosphatase PP2C (AP2C1), an Arabidopsis Ser/Thr phosphatase of type 2C, is a stress signal regulator that inactivates the stress-responsive MAPKs MPK4 and MPK6 [86, 87]. Mutant *ap2c1* plants produce significantly higher amounts of jasmonate upon wounding and are more resistant to phytophagous mites (*Tetranychus urticae*). Plants with increased AP2C1 levels display lower wound activation of MAPKs, reduced ethylene production, and compromised innate immunity against the necrotrophic pathogen *Botrytis cinerea*. Phosphorylation by a MAPK module regulates Kin7 superfamily members Hinkel and Tetraspore, which impinge on cell plate formation [88].

The Spc25 is part of the centromere assembly complex. KIN7.3 is a centromeric protein- E homolog and thus this interaction as well might be biologically relevant, especially during cell division. Spc25, is a component of the NDC80 complex, in mouse oocytes, an outer kinetochore complex comprising four subunits (ndc80/Hec1, nuf2, spc24, and spc25) [89]. The NDC80 constitutes one of the core MT-binding sites within the kinetochore. One hypothesis would be that KISC during cell division is involved in MT stabilization on the centromeres. Among the components of the outer kinetochore complex, the four proteins in the NDC80 complex, including NDC80 [nuclear division cycle 80, also known as Hec1 (highly expressed in cancer1) in humans], NUF2 (nuclear filament-containing protein 2), SPC24 (spindle pole body component 24) and SPC25, play critical roles in connecting spindle fibers to chromosomes. In vertebrates, the C- terminal ends of the NUF2-NDC80 heterodimer associate with the N-terminal coiled-coil domains of the SPC24-SPC25 heterodimer [90]. Globular dimeric heads, containing the RWD (RING finger, WD repeat, DEAD-like helicases) domain of the SPC24-SPC25 dimer, bind to the inner kinetochore components, the KNL1-Mis12 complex. Therefore, the NDC80 complex serves as a central ‘hub’ connecting the kinetochore complexes, which provides the ‘bridges’ between the inner kinetochore complexes and the MTs of the spindle.

The PORCINO protein is a tubulin folding cofactor and its absence results in embryo lethality [84]. The importance of KIN7.3 interaction with PORCINO (POR) remains to be established. The TFC C ortholog, PORCINO belongs to the four *PILZ* group genes encoding orthologs of mammalian tubulin-folding cofactors (TFCs) C, D, and E, and associated small G- protein Arl2 that mediate the formation of α/β-tubulin heterodimers *in vitro*. PORCINO was detected in cytosolic protein complexes and did not colocalize with MTs. Consistently, with previous reports, the *por* mutant was embryo lethal and the POR localized in the cytoplasm.

**Fig 1.**
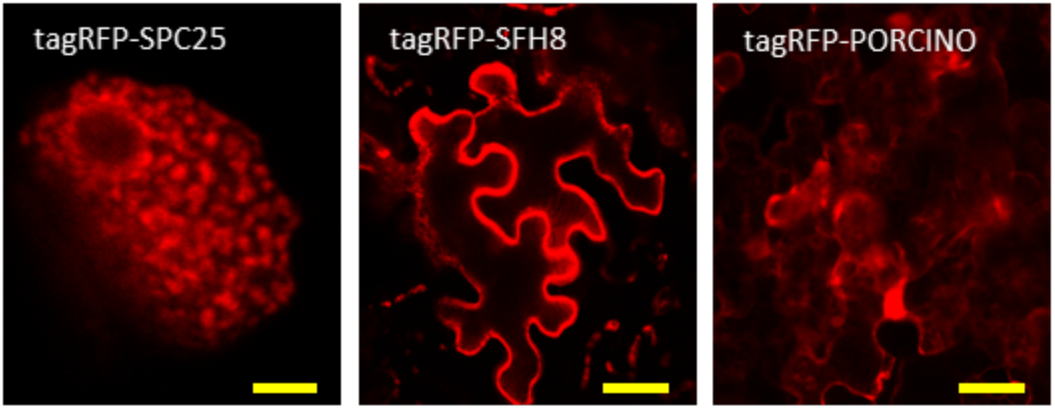
Localization of KIN7.3 interactors. Localization of SPC25, SFH8, and PORCINO tagged N-terminally with tagRPF in *N. benthamiana* transient system. Micrographs are representative of an experiment replicated three times results. Scale bars, 20 μm.

### Cleavage of Cohesin Subunit SYN4 by Separase

Sister-chromatid cohesion depends on cohesin, a 4-subunit protein complex that links the sisters as they are synthesized in the S phase. In anaphase when the protease separase cleaves the Scc1/ Mcd1 subunit of cohesin (or its Rec8 counterpart in meiotic cells), thereby allowing the sisters to be pulled apart by the mitotic spindle. The *Arabidopsis* genome encodes four kleisin subunits: the meiosis-specific ones SYN1 and SYN3, involved in gene expression of meiotic genes, but also expressed in somatic cells, and SYN2 and SYN4, which have been suggested to participate in mitotic cell division [29]. As we have shown previously, plants expressing tagged SYN proteins, except the mitotic cohesin SYN4, were highly susceptible to even mild environmental changes and showed decreased fertility [29].

### Principle of the R2D2 Probe

R2D2 shows a diminishing green but not red fluorescence when auxin levels are high, as the conserved domain II (DII) marker (*RPS5Apro*-driven DII fused to n3×Venus) is rapidly degraded in response to auxin. On the contrary, *RPS5Apro*-driven mutated DII-ntdTomato (red fluorescence) is not responding to auxin, thereby allowing the ratiometric fluorescence quantification of auxin response levels through the R2D2 [52].

## Notes

### Competing Interest Statement

The authors have declared no competing interest.

### Summary of Updates

Note added by authors: in the previous version of this manuscript the fig. S5 not present in this version was labelled inappropriately. The authors apologize for any inconvenience. For further information, please contact the corresponding author.

